# Host-aware RNA-based control of synthetic microbial consortia

**DOI:** 10.1101/2023.05.15.540816

**Authors:** Alice Boo, Harman Mehta, Rodrigo Ledesma Amaro, Guy-Bart Stan

## Abstract

Microbial consortia have been utilised for centuries to produce fermented foods and have great potential in applications such as therapeutics, biomaterials, fertilisers, and biobased production. Working together, microbes become specialized and perform complex tasks more efficiently, strengthening both cooperation and stability of the microbial community. However, imbalanced proportions of microbial community members can lead to unoptimized and diminished yields in biotechnology. To address this, we developed a burden-aware RNA-based multicellular feedback control system that stabilises and tunes coculture compositions. The system consists of three modules: a quorum sensing-based communication module to provide information about the densities of cocultured strains, an RNA-based comparator module to compare the ratio of densities of both strains to a pre-set desired ratio, and a customisable growth module that relies either on heterologous gene expression or on CRISPRi knockdowns to tune growth rates. We demonstrated that heterologous expression burden could be used to stabilise composition in a two-member *E. coli* coculture. This is the first coculture composition controller that does not rely on toxins or syntrophy for growth regulation and uses RNA sequestration to stabilise and control coculture composition. This work provides a fundamental basis to explore burden-aware multicellular feedback control strategies for robust stabilisation of synthetic community compositions.

## Introduction

Over the past decade, there has been a growing interest in the interactions between microbes and their environment, leading to an explosion of publications on the microbiome. In 2020, over 20,000 articles were published on the microbiome ^1^, driven by the quest to uncover the causality between disease and microbiota for the development of microbiome-based therapeutics. Beyond the human microbiome, microbial communities have gained attention for their potential in various fields, including agriculture ^2, 3^, bioremediation, food production and waste valorisation ^4^. Synthetic biology has enabled the engineering of microbial communities, which can be created or manipulated to perform enhanced or new functions by exploiting the strengths and specificities of each microbe. By dividing complex tasks into smaller ones, the energy expenditure of each microbe is minimised, allowing community microbes to grow better and achieve higher production yields of complex compounds than monocultures ^5–10^. It also enables the bioproduction of high-value molecules from waste products such as plastic, methane or waste derived from the agriculture and food industries ^11–13^. Additionally, compartmentalising genetic circuits and pathways across different microbial species increases the modularity and reusability of genetic parts and modules, thus opening new avenues for distributed biocomputing and multicellular control strategies ^14–20^. These features of engineered microbial communities position them ideally to be explored for the development of sustainable innovations as the bioeconomy continues to grow.

However, the principle of competitive exclusion presents a challenge in establishing stable microbial communities, as it limits the coexistence of microbial species ^21^. Current strategies mostly rely on syntrophic relationships ^7, 12, 22–24^, but they lack the flexibility needed to tune population composition, which is critical for optimal metabolic pathway expression. Varying seeding ratios is a common approach to selecting the best initial conditions for optimal coculture yields, but this strategy often offers little control on the dynamics of the population over time, thus limiting its application ^11, 25–30^. There is currently a lack of understanding regarding the potential benefits of dynamically adjusting these compositions over time, which is compounded by the fact that developing reliable gene circuits that can effectively tune and control the population composition of microbial consortia is a significant challenge. Quorum sensing-based systems, which reflect population density, have been vastly used to engineer such circuits to control microbial consortia dynamics ^6, 31–37^ by placing a growth regulation mechanism downstream of a quorum sensing promoter. The use of lysis proteins ^38–40^ as growth regulators has shown robust temporal control of coculture composition and promising results for the development of anti-cancer treatments. However, they are not well suited for bioproduction in large bioreactors as biomass is constantly lysed and regenerated over time. Toxins and antimicrobials ^41–45^ as growth regulators have also demonstrated promising results for controlling coculture dynamics. Recently, novel feedback approaches for controlling microbial consortia have emerged, employing single-strain control strategies through chemical induction ^44^ and optogenetics ^46^ integrated with titration mechanisms for antimicrobial resistance. These studies have demonstrated that engineering a single strain is sufficient to regulate the composition of a two-strain microbial consortium. Optogenetics offers the advantages of rapid control over system dynamics and the ability to easily adjust composition during experiments. Additionally, manipulating the intake of essential amino acids or sugars ^30, 47, 48^ has shown promise in governing coculture ratios. However, it is important to note that the expression of these control circuits themselves can impose a burden on cell growth, leading to reduced achievable ranges of population composition ^47^. This undesirable effect, along with the use of toxins, antimicrobials, and burdensome control strategies, calls for the exploration of new approaches that can be upscaled to industrial production without compromising growth rates, and in turn bioproduction capacity ^49–53^.

This work proposes an RNA-based control system to regulate the population composition in a two-member *E. coli* coculture. The design includes three core modules: 1) a communication module using quorum sensing to track the density of the cocultured strains, 2) a comparator module to evaluate the difference between the quorum sensing signal and a reference density used to compare the ratio of both strains to a pre-set desired ratio, and 3) a growth module that tunes exogeneous gene expression to modulate growth and achieve the desired population composition (**Figure 1**). The study uses STAR ^54, 55^, a cis-acting RNA-based transcription regulation device, to build a novel STAR-based comparator that estimates the difference between population sizes and acts to restore the population composition to the desired ratio when it diverges. RNA-based systems are an attractive alternative to burdensome protein-based designs as they enable predictable RNA-RNA Watson-Crick base pairings, are highly orthogonal, and enable transduction of information at the RNA level ^56–58^. Importantly, RNA systems theoretically use minimal host resources as they do not require translation to proteins, one of the costliest cellular processes in fast dividing organisms ^50, 52^. RNA-based systems hence offer a promising alternative for the design of burden-aware controllers. Overall, this work presents, for the first time, a quorum and RNA-based control system for the autonomous and robust control of population composition.

**Figure 1.**
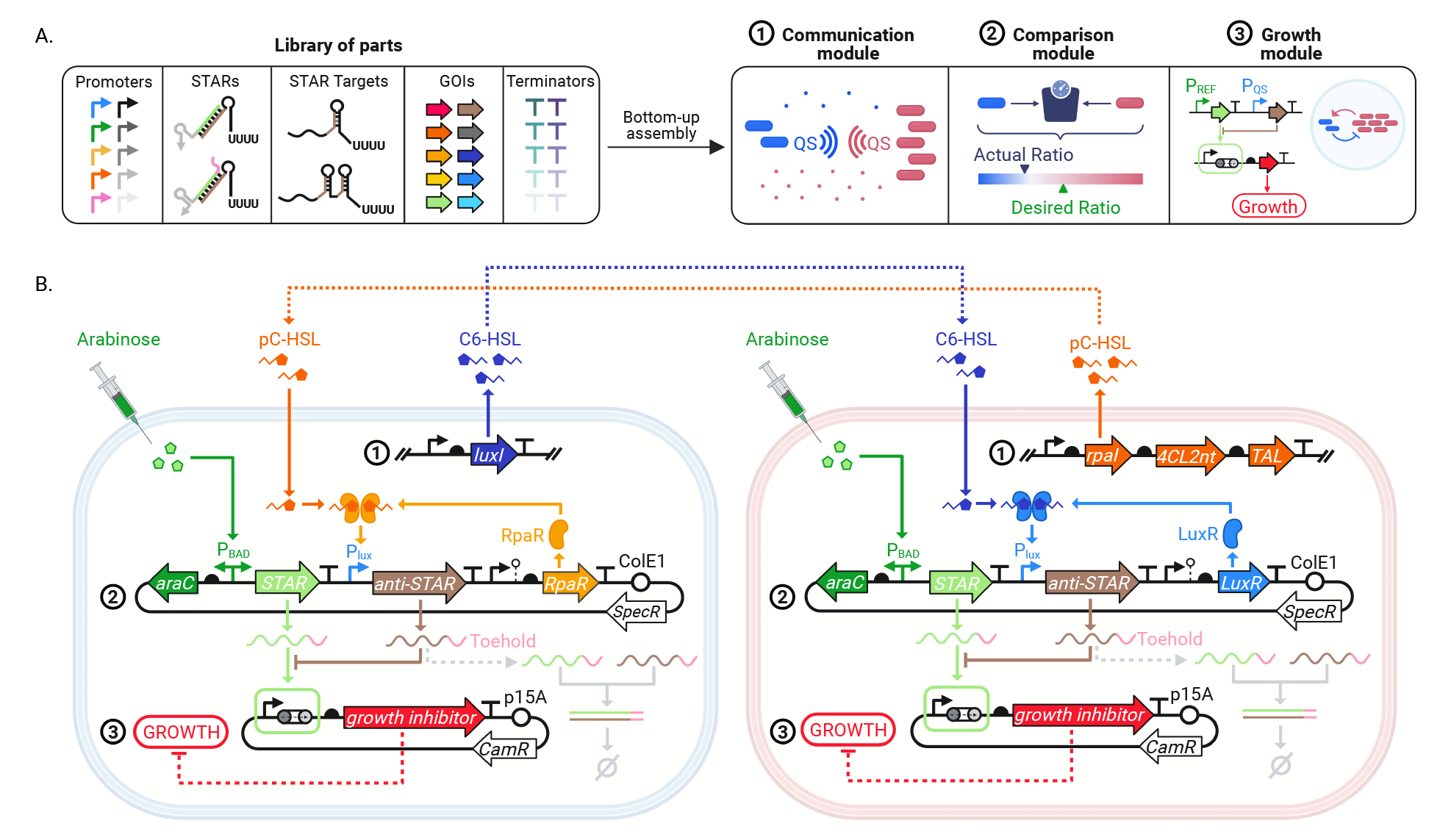
An RNA-based genetic circuit for stabilising population composition in a microbial coculture system. **(A)** Three modules were built using a bottom-up part assembly strategy to engineer a system controlling population composition in a two-strain *E. coli* consortium. The communication module propagates information about the population density of each of the two strains. The RNA-based comparator built from a set of STAR and anti-STAR parts, is designed to compare bacterial population density to an inducible reference signal. The output signal of the comparator is used by the growth module to determine whether the cell density of each strain should be up-or down-regulated to maintain a stable coculture composition. **(B)** Proposed circuit for controlling community composition. Each bacterial population produces a specific quorum sensing molecule (e.g. C6-HSL or pC-HSL) that reflects population density and can be detected by the other population. A synthetic RNA (anti-STAR) is produced upon detection of quorum molecules in each population and is compared to a reference signal (STAR). STAR and anti-STAR are designed to bind to each other and form an inactive complex. Free STAR can bind to the termination hairpin to allow transcription of the growth regulation gene. Anti-STAR acts as a STAR-sequestration buffer, preventing STAR from binding to the termination hairpin.

## Results

### 1. A three-module architecture for tuning population composition in a microbial coculture

Inspired by protein-based sequestration mechanisms such as sigma and anti-sigma factors ^59^, we designed a general architecture using RNA-RNA interactions to control population composition of a two-member *E. coli* coculture. Commonly, synthetic genes circuits are assembled using a bottom-up approach that consists of connecting characterised parts together to form modules, which when combined, form genetic devices ^60, 61^. From a selected set of parts used to express quorum sensing molecules, regulatory RNAs and growth regulators, we assembled three modules: a communication module, a comparison module and a growth module (**Figure 1A**). The communication module relies on quorum sensing molecules to estimate the density of bacterial populations. This information is used by the comparison module to estimate the differences between actual and desired ratios of the two cell types. Using this information, the growth module modulates cell growth in both populations so as to minimise the ratiometric error. In **Figure 1B**, we present the general architecture of the RNA-based sequestration mechanism that can be used to regulate coculture composition. C6-HSL and pC-HSL AI- 1 quorum sensing molecules were selected to convey information about population densities as they were previously reported to be signal orthogonal, i.e. they are orthogonal as long as their cognate regulator proteins LuxR and RpaR are physically compartmentalised ^34, 35, 62^. Additionally, Scott et al. showed that the LLL quorum sensing system, i.e. the system in which C6-HSL binds to the regulator LuxR and activates the lux promoter (pLux), exhibits similar properties to the LRR system, for which pC-HSL binds to the RpaR regulator to activate pLux ^35^. C6-HSL production by C6-HSL sender strains requires the expression of a single enzyme, HSL synthase LuxI from *Vibrio fischeri* ^63–65^, while pC-HSL production by pC-HSL sender strains requires the expression of three enzymes: tyrosine ammonia lyase (*TAL*) from *Saccharothrix espanaensis,* 4-coumarate-CoA ligase from *Nicotiana tabacum* (*4CL2nt*) and HSL synthase RpaI, from *Rhodopseudomonas palustris* ^66^. The comparator relies on the sequestration of two RNA species: STAR and anti-STAR. STAR expression is controlled by an inducible promoter, which sets the desired ratio of the coculture. For example, to stabilise the coculture composition around a 1:1 ratio, we can control STAR expression in both strains by the same inducible promoter, e.g. the arabinose promoter (pBAD). Anti-STAR is controlled by the quorum sensing signal representing the cell density of the opposite strain. Therefore, as anti-STAR sequesters STAR into a complex degraded by the cell’s native RNAse E, less STAR is available to bind to the STAR target, thus reducing the expression of the downstream growth regulator or gene of interest. To minimise the toxicity of the multicellular feedback, we decided to either regulate the cell growth rate through an RNA-mediated essential gene knockdown or by modulating the burden of the gene(s) of interest, thus the circuit did not require an additional gene to regulate growth rate beyond the exogenous gene(s) required to perform the function of interest in the microbial consortium. To validate whether burden could be successfully used for regulating co-culture compositions, we constructed a mechanistic mathematic model describing the three modules and their impact on the growth of two bacterial species sharing a single growth compartment (**Supplementary Figure 1, Supplementary Note 1**). As opposed to previously described co-culture control strategies involving the expression of a toxin that negatively impacts bacterial growth rates, burden slows down growth proportionally to the concentration of the burdensome protein expressed by the bacterial populations ^44, 67, 68^. We found that our strategy successfully stabilised the co-culture composition when different burdens were imposed on the two populations. In addition, the co-culture ratio was tuneable by varying the expression of STAR and anti-STAR in the system.

### 2. Designing an RNA-based molecular sequestration system to stabilise coculture composition

We engineered a set of sender strains that produce either C6-HSL or pC-HSL by expressing *LuxI* or the 3-gene rpa operon respectively under varying promoter and RBS strengths. These genes were genomically integrated to minimise the impact of their expression on the growth of the *E. coli* DH10B strain (**Supplementary Figure 2**). We designed an assay to characterise the transcription activation profile of the quorum sensing lux promoter in response to varying concentrations of homoserine lactone (HSL) molecules produced by the sender strains (**Figure 2A**). To do so, we used dBroccoli, a fluorescent RNA aptamer that was previously used to characterise mRNA expression in *E. coli* cells and that does not impact cell growth upon induction (**Supplementary Figure 3**) ^69^. To quantify C6-HSL when expressing the LuxR regulator (or pC-HSL when expressing RpaR), we created receiver plasmids where dBroccoli is placed downstream of the lux promoter (**Supplementary Figure 4**). To test whether the receiver strains could detect the concentrations of HSL produced by the sender strains, we grew the sender strains separately for 1 to 6 hours and collected their supernatants by centrifugation. We then mixed the supernatant with the LLL and LRR receiver strains and measured the fluorescence emitted by dBroccoli. All receiver strains could detect a significant difference in HSL concentration between the supernatants collected at 1 and 6 hours for all strains, except for *Lux5* and *Rpa6* which already produced saturating amounts of HSL after 1 hour of growth (**Figure 2B**). Temporal responses of the receiver strains to the quorum sensing produced by *Lux1* and *Rpa5* strains are show in **Supplementary Figure 5**. The strain libraries would therefore serve as a basis to tune the strength of the communication signals between the strains grown into a coculture.

**Figure 2.**
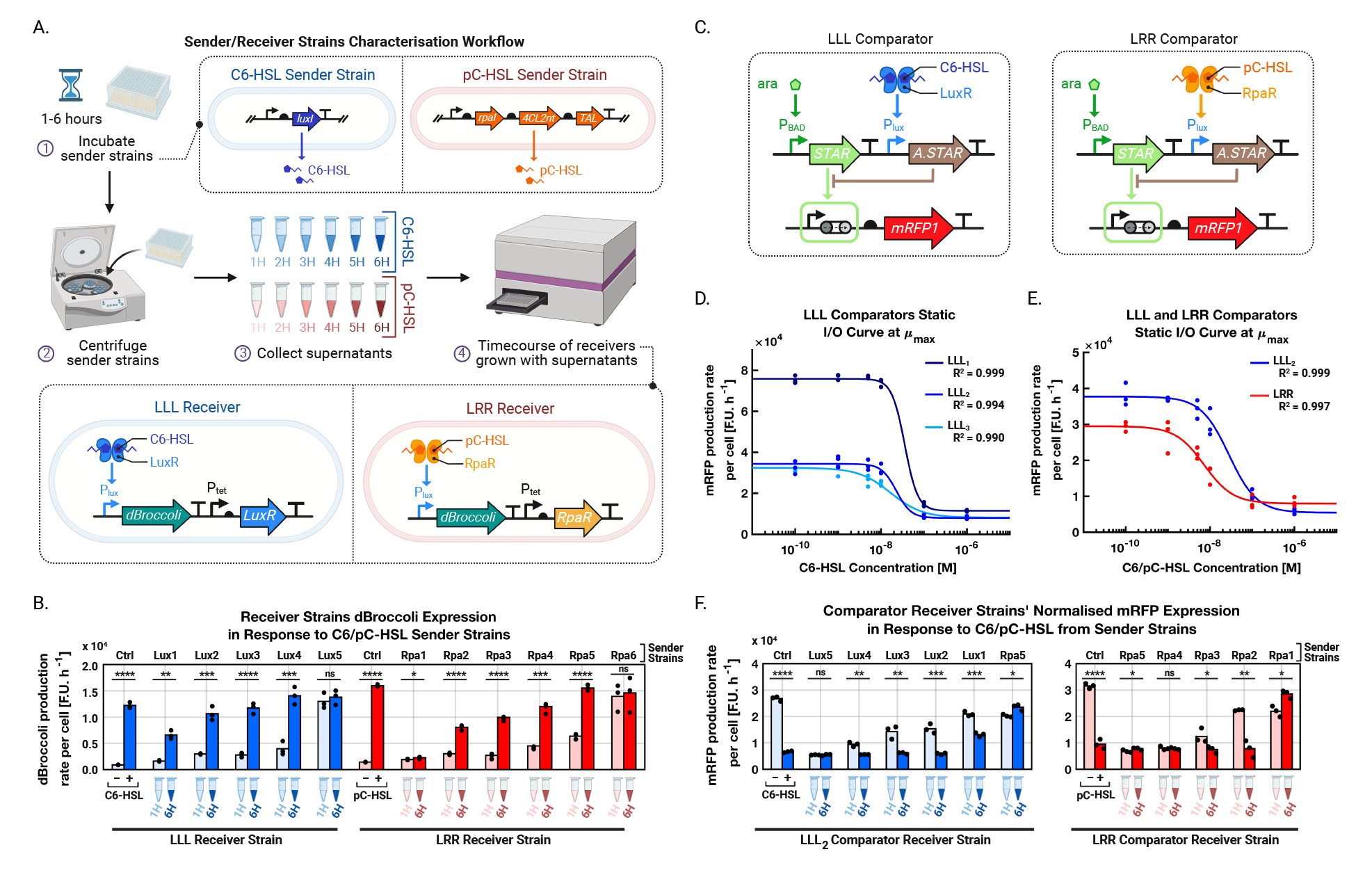
Comparing population density using quorum sensing and a STAR-based comparator. **(A)** Workflow for the characterisation of the HSL sender/receiver pairs. Sender strains producing either C6-HSL or pC-HSL, respectively, are incubated for a period of 1 to 6 hours before being centrifuged for their supernatants to be collected every hour and mixed with the appropriate C6-HSL and pC-HSL receiver strains. The response of the receiver strains to the HSL produced by the sender strains is monitored using a plate-reader assay detecting dBroccoli fluorescence. The LLL receiver detects C6-HSL upon binding with the LuxR regulator, while the LRR receiver detects pC-HSL upon binding with the RpaR regulator. **(B)** dBroccoli expression of LLL and LRR receiver strains in response to sets of C6-HSL-and pC-HSL-producing strains denoted as Lux and Rpa respectively (strains are described in Supplementary Table 1 and plasmids in Supplementary Table 3). The LLL and LRR receiver strains detect quorum sensing molecules produced by HSL-producing strains grown for either 1 hour or 6 hours. Controls are the LLL and LRR sender strains grown in DH10B supernatant supplemented with 0 M of HSL (-) or 10^-7^M of HSL (+). **(C)** Genetic circuits of the LLL and LRR STAR-based comparators (Supplementary Figure 7). The LLL and LRR comparators compare an L-arabinose inducible reference signal to C6-HSL and pC-HSL signals, respectively. **(D)** Input-output static response curves of the LLL comparators (Supplementary Table 3) as C6-HSL is externally added to the culture. The I/O properties of the three LLL comparators depend on the toehold sequence and concentration of LuxR in the cell. The mRFP production rate is displayed at the time corresponding to the maximum growth rate of the host cell (μ_max_). **(E)** Static-input output response curves of the LLL_2_ and LRR comparators as C6-HSL or pC-HSL are respectively externally added to the cultures. **(F)** Response of the LLL_2_ and LRR comparators to the quorum sensing concentration present in the supernatants of the C6-HSL and pC-HSL sender strains collected after 1 hour and 6 hours of growth. Controls are the LLL and LRR sender strains grown in DH10B supernatant supplemented with 0 M of HSL (-) or 10^-7^M of HSL (+). Curves were fitted using MATLAB four-parameter nonlinear regression fit. Data represent the mean values of n = 3 biological replicates. Statistically significant differences were determined using two-tailed Student’s t-test (**** represents p<0.0001, *** represents p<0.001, ** represents p<0.01, * represents p<0.1, ns represents not significant). For all panels, OD and fluorescence data were collected using a microplate reader.

Having established a two-way communication system capable of producing and detecting C6-HSL and pC-HSL, we next explored how to link it to the production of anti-STAR as part of the design of the comparator module. For this, two versions of the comparator were implemented, one producing anti-STAR in response to C6-HSL (LLL comparator) and the other in response to pC-HSL (LRR comparator) (**Figure 2C**). To test the comparator, STAR was placed under the control of the strong arabinose-inducible araBAD promoter and controlled the activation of mRFP1 expression. We built a small library of STAR and anti-STAR with six different toehold sequences from Green et al. ^70^ (**Supplementary Figure 6**). We also tested comparator variants with varying RBS strength driving LuxR. Here after, the LLL_2_ comparator expresses STAR and anti-STAR with toehold 2 (pAB300), LLL_1_ expresses STAR and anti-STAR without a toehold (pAB317) and LLL_3_ is a variant of LLL_2_ with *luxR* expressed with a stronger RBS (pAB545) (**Supplementary Table 3**). We observed that changing the hybridization energy between STAR and anti-STAR changed the ON/OFF properties of the comparator as demonstrated by the LLL_2_ and LLL_1_ comparators results presented in **Figure 2D**. When only expressing STAR from the LLL_1_ and LLL_2_ designs, mRFP production rate was 2-fold higher for LLL_1_ than for LLL_2_ (**Figure 2D, Supplementary Figure 7**). The properties of the comparator can also be tuned by increasing the expression of the LuxR regulator, which changes the slope of the input-output response curve as shown by results from the LLL_2_ and LLL_3_ comparators, which share the same toehold sequence (**Figure 2D**). We found that using a tandem STAR termination hairpin reduced both the OFF and ON states of the comparator (**Supplementary Figure 8**). As the ON state of this design was low, we did not use it further in this work, but we hypothesised that this design could be suitable for specific applications such as for expressing highly toxic products which require very tight and low gene expression. The LRR comparator, which uses the same STAR and anti-STAR design as the LLL_2_ comparator, has a similar operating range as that of LLL_2_ (Figure 2E). However, the deactivation of STAR by anti-STAR was more efficient for the LLL_2_ comparator than for the LRR comparator, with a deactivation percentage of 93% for LLL_2_ compared to 78% for LRR (**Supplementary Figure 9**). It is therefore possible to tune the output of the comparator by playing with the comparator’s RNA-RNA hybridization energy properties as well as by tuning the level of anti-STAR expression by using weaker or stronger HSL producers (**Figure 2F**).

### 3. Controlling growth rates with the output of the RNA-based comparator

Having demonstrated the RNA-based comparator can be used to modulate the level of expression in response to the concentration difference of two signal-orthogonal quorum sensing molecules, we investigated how to control bacterial growth rate while using a minimal amount of host resources. For this, we showed that the comparator can be coupled to the expression of a gene of interest, which in turn impacts cell growth rate (**Figure 3**). We investigated four types of growth regulators: (1) a small heterologous protein: eforRed, (2) a large heterologous protein: VioB, (3) a metabolic pathway to express β-carotene, and (4) CRISPRi targeting *E. coli*’s native leucine operon.

**Figure 3.**
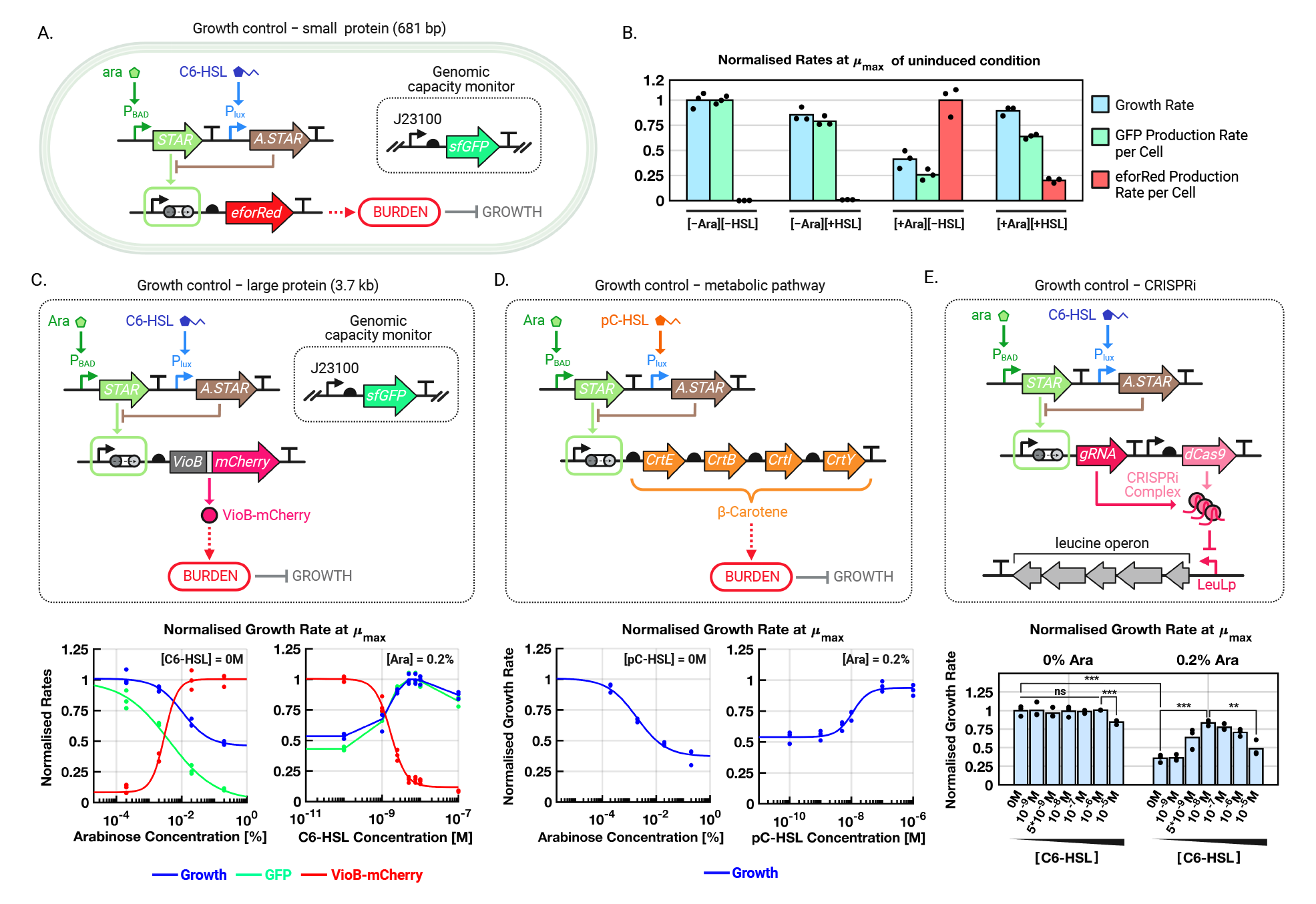
The STAR-based comparator controls growth rate depending on the input concentration of a quorum sensing molecule. **(A)** The DH10B-GFP capacity monitor strain (Supplementary Table 1) is used to monitor gene expression burden caused by eforRed chromoprotein production when the STAR-based comparator activates the growth module. All plasmid descriptions are available in Supplementary Table 3. **(B)** Normalised growth rate, GFP production rate per cell and eforRed production rate per cell of the circuit shown in A at the time corresponding to the maximum growth rate of the host cell (μ_max_). EforRed is expressed in the presence of L-arabinose, which induces gene expression burden observable by a reduction in GFP production rate per cell and growth rate. In the presence of both L-arabinose and C6-HSL, anti-STAR sequesters STAR, thus reducing eforRed production, attenuating gene expression burden and recovering growth rate. L-arabinose and C6-HSL were externally added at the following concentrations: 0% and 0.2% L-arabinose for [-Ara] and [+Ara] respectively, and 0 M and 10^-7^ M of C6-HSL for [-HSL] and [+HSL] respectively. **(C)** The comparator controls growth rate by tuning the expression of VioB-mCherry, a large heterologous protein, in response to L-arabinose and C6- HSL in DH10B (Supplementary Tables 1 and 3). In the left-hand side panel, cultures were externally induced with 0M of C6-HSL and increasing concentrations of L-arabinose. In the right-hand side panel, cultures were externally induced with 0.2% of L-arabinose and increasing concentrations of C6-HSL. Growth rate and GFP production rate per cell values were normalised against the uninduced state of the circuit (0 M of C6-HSL and 0% of L-arabinose). The VioB-mCherry production rate per cell values were normalised against the STAR-induced and anti-STAR-uninduced condition (0 M of C6-HSL and 0.2% of L-arabinose). Data represent the mean values of n = 3 biological replicates shown as individual dots. **(D)** The comparator controls growth rate by tuning the expression of a four-gene metabolic pathway producing β-carotene, in response to L-arabinose and pC-HSL in DH10B (Supplementary Tables 1 and 3). In the left-hand side panel, cultures were externally induced with 0M of pC-HSL and increasing concentrations of L-arabinose. In the right-hand side panel, cultures were externally induced with 0.2% of L-arabinose and increasing concentrations of pC-HSL. Growth rate and GFP production rate per cell values were normalised against the uninduced state of the circuit (0 M of pC-HSL and 0% of L-arabinose). Data represent the mean values of n = 3 biological replicates shown as individual dots. **(E)** The comparator controls growth rate by tuning the expression of a gRNA targeting the LeuLp genomic promoter driving expression of the leucine operon in BW25113 (Supplementary Tables 1 and 3). The cultures were externally induced with either 0% or 0.2% of L-arabinose and increasing concentrations of C6-HSL. Bars represent the mean values of n = 3 biological replicates shown as individual dots. Statistically significant differences were determined using two-tailed Student’s t-test (**** represents p<0.0001, *** represents p<0.001, ** represents p<0.01, * represents p<0.1, ns represents not significant). For all panels, OD and fluorescence data were collected using a microplate reader.

First, we used the capacity monitor strain, an *E. coli* strain genomically integrated with a constitutively expressed GFP previously developed by *Ceroni et al.*^50^, to monitor the burden caused by expressing a small heterologous gene of interest, eforRed. As expression of eforRed increases, cellular resources are pulled away from other processes, thus decreasing GFP expression from the capacity monitor. We observed that inducing eforRed expression reduced GFP capacity by 80% and growth rate by 60%. In addition, when coupling eforRed expression to our comparator, in the presence of C6-HSL, anti-STAR is maximally expressed and GFP capacity is recovered up to 60% of its original value, while growth rate is recovered up to 90% of the original value observed when eforRed is not expressed (**Figure 3B**). We note that GFP capacity is not fully recovered when anti-STAR is expressed and that capacity when anti-STAR alone is expressed is lower than when anti-STAR is not expressed. This reflects the cost of expressing the controller species, STAR and anti-STAR (**Supplementary Figure 10**). To further explore if burden could be used to regulate growth rate, we expressed VioB-mCherry, a large fusion protein previously shown to impose burden on *E. coli* (**Figure 3C**). Cellular growth rate was reduced by 53% when expressing VioB-mCherry. Sequestration by anti-STAR led to restoring the host growth rate up to 90% of the original value measured when STAR is not expressed, i.e. in the absence of the STAR inducer, L-arabinose (**Figure 3C, Supplementary Figure 11**). We note that the comparator could only regulate growth rate by tuning VioB-mCherry expression if enough VioB-mCherry was expressed by the system (**Supplementary Figure 11**) ^50, 71^. The comparator could also tune burden, and by extension growth rate, caused by the expression metabolic pathways such as the β-carotene pathway (**Figure 3D** and **Supplementary Figure 12**) ^72^. Expressing the β-carotene pathway resulted in a maximum decrease in growth rate of about 50 to 60% when inducing the circuit with 0.2% L-arabinose. Expressing anti-STAR by inducing the system with 1 µM pC-HSL was able to restore growth rate up to 90% of the original value measured when no STAR was expressed (0% L-arabinose). Finally, we linked the output of the comparator to a CRISPRi system targeting the leucine operon to build a tuneable amino acid knockdown (**Figure 3E**). We designed a gRNA targeting the native LeuLp promoter driving BW25113’s leucine operon (**Supplementary Figures 13**) and repress cellular growth (**Supplementary Figure 14**). However, repression of the leucine operon with a fully complementary guide sequence to the LeuLp promoter gave rise to extended lag-phases of over 20 hours but not growth rate reduction (**Supplementary Figures 15A-15C**). As an extended lag phase is not a desirable property to build our multicellular feedback controller, we introduced a single base-pair mutation in the guide sequence to tune CRISPRi inhibition level of the leucine pathway and found that a mismatch preceding the PAM sequence could alleviate CRISPRi repression and shorten the lag-phase (**Supplementary Figures 15D-F, 16**). By regulating the expression level of the gRNA, the comparator could reduce cellular growth rate by 64%, and STAR sequestration by anti-STAR was able to recover growth rate up to 80% of its original value when no gRNA was expressed, i.e. in the absence of L-arabinose.

Taken together, these approaches demonstrate that the STAR and anti-STAR sequestration system can successfully up-and down-regulate growth rate following a quorum sensing input. We envision that the controller could be tuned further via the addition of external molecules, such as L-leucine, which inhibits *E. coli* K-12 strains growth in the absence of L-isoleucine (**Supplementary Figure 17**). This would give an additional level of control to temporally adjust the population composition regulated autonomously by the STAR-based controller.

### 4. Stabilising the ratio of an unbalanced *E. coli* coculture using gene expression burden

Next, we connected the three modules – communication, comparison and growth regulation – to demonstrate how an RNA-based sequestration controller can stabilise the coculture composition of an unbalanced coculture system around a desired ratio. For this, we used a coculture expressing VioB-CFP and VioB-YFP under the control of comparators expressing STAR from the rhaBAD and the araBAD promoters, respectively (**Figure 4A**). To assess the performance of our multicellular feedback, we built two versions of the controller: a closed-loop and an open-loop version. In the closed-loop coculture (CL), the CFP strain produces C6-HSL while the YFP strain produces pC-HSL. The open-loop coculture (OL) carries the same circuit as that of the CL, however neither strain produce quorum sensing molecules and, as a result, the controller does not receive information about the strains’ densities.

**Figure 4.**
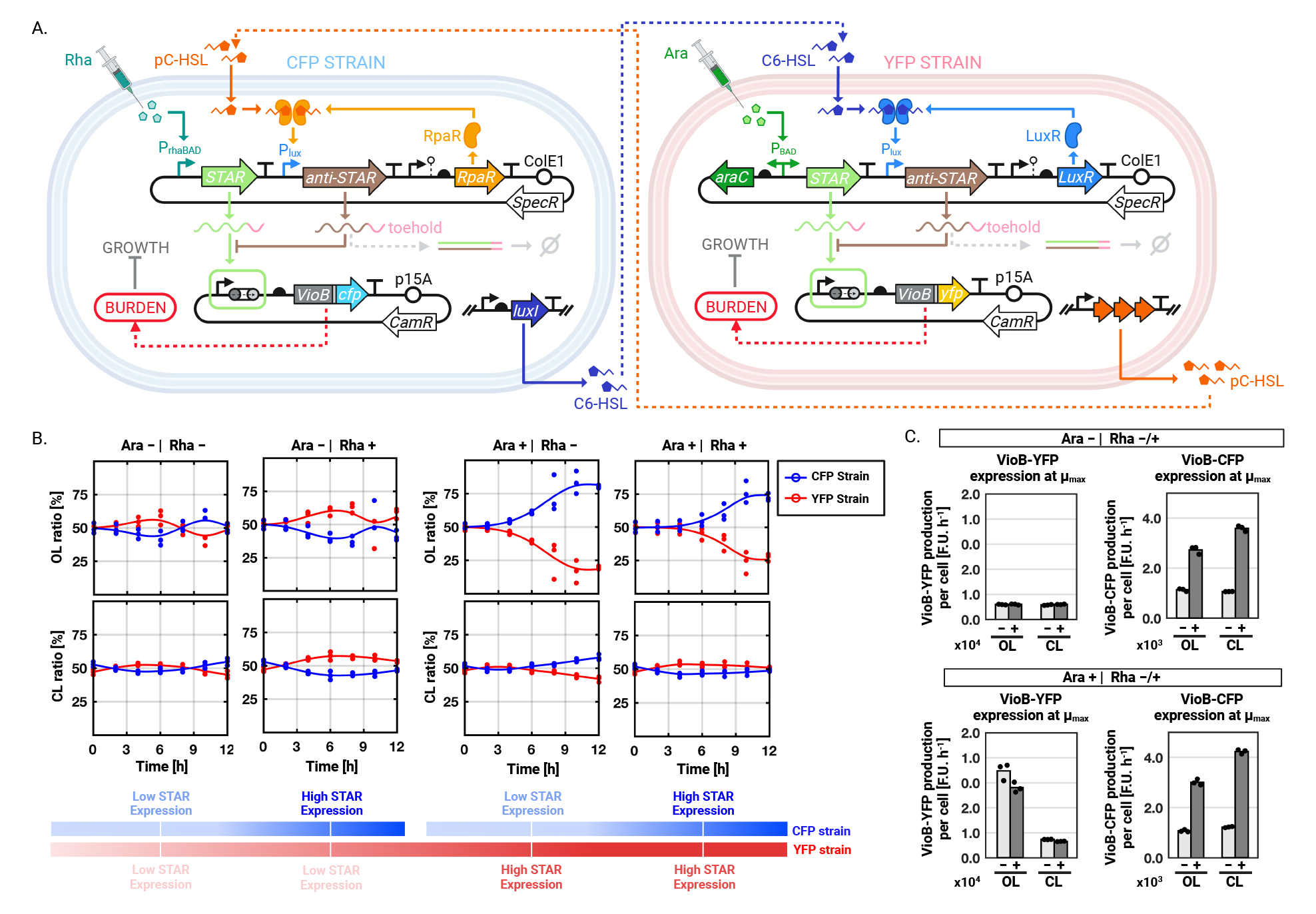
Stabilisation of coculture composition using a burden-driven growth control strategy. **(A)** Circuit of the closed-loop two-member *E. coli* coculture. The CFP strain produces C6-HSL that inhibits VioB-YFP expression in the YFP strain, while the YFP strain produces pC-HSL that inhibits VioB-CFP expression in the CFP strain. Growth rate is controlled by the STAR-based comparator by tuning gene expression burden caused by VioB-YFP and VioB-CFP expression. **(B)** Population composition of the open-loop (OL) and closed-loop (CL) cocultures when induced with combinations of 0% (Rha-) and 0.2% (Rha+) of L-rhamnose and of 0% (Ara-) and 0.2% (Ara+) of L-arabinose. The OL design was composed of DH10B carrying the pAB537 and pAB519 plasmids and DH10B-mScarlet carrying the pAB317 and pAB518 plasmids, and the CL design was composed of *Lux_5_* carrying the pAB537 and pAB519 plasmids and *Rpa_5_* carrying the pAB317 and pAB518 plasmids (Supplementary Tables 1 and 3). In the presence of L-arabinose, the CL coculture corresponds to the circuit depicted in panel A. The OL coculture corresponds to the circuit in panel A in which neither the CFP Strain nor the YFP Strain produce quorum sensing molecules. Data represent the mean values of n = 3 biological replicates shown as individual dots. **(C)** VioB-YFP and VioB-CFP production rate per cell at maximum growth rate μ_max_ of the cocultures from panel B. For all experiments, the concentrations of L-rhamnose and L-arabinose used for induction are: 0% and 0.2% of L-rhamnose for (-Rha) and (+Rha), respectively; 0% and 0.2% of L-arabinose for (-Ara) and (+Ara), respectively. Bars represent the mean values of n = 3 biological replicates shown as individual dots. For all panels, coculture composition was determined by flow cytometry. OD and fluorescence data were collected using a microplate reader.

In **Figure 4B**, we showed that for the OL cocultures, as we externally supply neither, either or both L-rhamnose and L-arabinose to the coculture, expression of VioB in either or both strains drive the coculture composition out of equilibrium. However, in the CL system, exchange of quorum sensing molecules leads to the stabilisation of the coculture composition around a 1:1 ratio. This is achieved through down-regulation of the expression of VioB in the slowest growing strain – here the YFP Strain as, in the CL, VioB-CFP is not downregulated, while VioB-YFP is always downregulated (**Figure 4C**). The YFP strain growth rate is more affected by the expression of VioB than the CFP Strain for two reasons. First, the araBAD promoter is stronger than the rhaBAD promoter (**Supplementary Figure 18**) and, second, VioB-YFP expression is more burdensome than VioB-CFP expression (**Supplementary Figure 19**). The C6-HSL sender strain *Lux_5_* and the pC-HSL sender strain *Rpa_5_* were chosen as host for the CL circuit as weaker production of quorum sensing molecules did not lead to the stabilisation of the coculture composition around a 1:1 ratio (**Supplementary Figures 20 & 21**). We note however that tuning the expression rate of the quorum sensing molecules is a way to stabilise the coculture composition around a wider range of ratios beyond the 1:1 ratio demonstrated in **Figure 4**.

These results demonstrate that our CL circuit can stabilise coculture composition when the expression of two heterologous genes causes different levels of burden, thus differentially affecting the growth of the cocultured strains.

### 5. Demonstrating community composition tuneability and protein yield improvement

Finally, we explored key parameters of the sequestration-based controller that could be tuned to modulate the composition of the two-strain coculture (**Figure 5**). Key parameters of the STAR-based comparator include the production rates of both STAR and anti-STAR, as well as tuning the sequestration dynamics using different toehold domains. As previously mentioned, we explored the impact of HSL production by one of the sender strains on coculture behaviour. We selected the pC-HSL producing strains *Rpa_2_*, *Rpa_4_* and *Rpa_5_* from **Figure 2** to tune sender production of pC-HSL, thus modulating the gain of anti-STAR production in the pC-HSL receiver strain (**Supplementary Figure 21**). The R*pa_2_* strain is designated as a weak pC-HSL sender, *Rpa_4_* as a medium-strength pC-HSL sender and R*pa_5_* as a strong pC-HSL sender. As predicted, increasing the production of pC-HSL by the CFP sender strain, progressively decreased VioB-YFP expression (**Supplementary Figure 21**).

**Figure 5.**
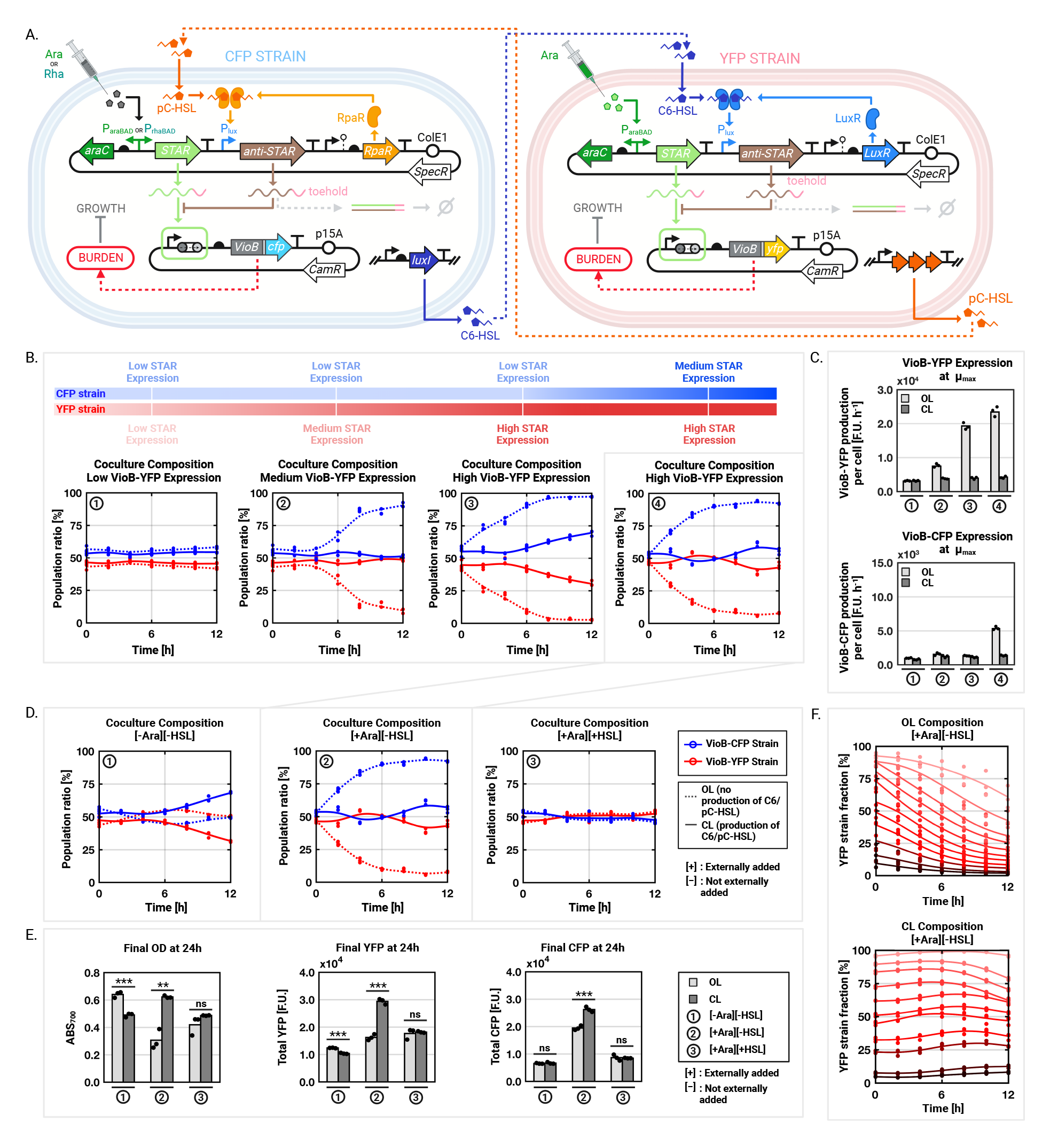
Demonstrating coculture composition tuneability. **(A)** Diagram of the circuit architecture used to demonstrate tuneability of the coculture composition. The coculture composition can be adjusted by tuning the I/O properties of the comparator. This can be done by tuning the expression levels of STAR and anti-STAR as well as using different toehold domains. **(B)** Effect of varying STAR expression in the CFP strain (DH10B for OL or Lux_5_ for CL) and the YFP strain (DH10B-mScarlet for OL or Rpa_5_ for CL) on coculture composition. Plasmids carried by the CFP and YFP strains are described in Supplementary Table 3. (1) The CFP strains carrying pAB519 and pAB537 and the YFP strains carrying pAB518 and pAB300 are externally induced with 0% of L-arabinose and 0% of L-rhamnose such that both strains do not produce either VioB-CFP or VioB-YFP. Both strains carry a medium-activating STAR design based on the toehold domain of the LLL_2_ comparator. (2) The CFP strains carrying pAB519 and pAB537 and the YFP strains carrying pAB518 and pAB300 are externally induced with 0% L-rhamnose and 0.2% L-arabinose such that only the YFP strain produces VioB-YFP in response to a medium-activating STAR design based on the toehold domain of the LLL_2_ comparator. (3) The CFP strains carrying pAB519 and pAB537 and the YFP strains carrying pAB518 and pAB317 are externally induced with 0% L-rhamnose and 0.2% L-arabinose such that only the YFP strain produces VioB in response to a strong-activating STAR design based on the toehold domain of the LLL_1_ comparator. (4) The CFP strains carrying pAB519 and pAB537 and the YFP strains carrying pAB518 and pAB317 are externally induced with 0.2% of L-rhamnose and 0.2% of L-arabinose such that the YFP strain produces VioB-YFP in response to a strong-activating STAR design (based on the toehold domain of the LLL_1_ comparator) and the CFP strain produces VioB-CFP in response to a medium-activating STAR design (based on the toehold domain of the LLL_2_ comparator). Values represent the mean values of n = 3 biological replicates shown as individual dots. **(C)** VioB-YFP and VioB-CFP production rate per cell at maximum growth rate μ_max_ of the cocultures from panel B. Bars represent the mean values of n = 3 biological replicates shown as individual dots. **(D)** Coculture (4) from Panel B tested in three inducer conditions: (1) 0% of L-arabinose, 0M of both C6-HSL and pC-HSL, (2) 0.2% of L-arabinose, 0M of both C6-HSL and pC-HSL, (3) 0.2% of L-arabinose, 10^-7^ M of both C6-HSL and pC-HSL. Values represent the mean values of n = 3 biological replicates shown as individual dots. **(E)** Final OD, VioB-YFP and VioB-CFP fluorescence taken after 24 hours of growing the three cocultures from panel D. **(F)** YFP strain fraction of the open-loop (OL) and closed-loop (CL) circuits from panel B.4. for different initial inoculation ratios. Both the OL and CL are induced with 0.2% of L-rhamnose and 0.2% of L-arabinose such that the YFP strain produces VioB-YFP in response to a strong-activating STAR design (based on the toehold domain of the LLL_1_ comparator) and the CFP strain produces VioB-CFP in response to a medium-activating STAR design (based on the toehold domain of the LLL_2_ comparator). Curves are fitted to the mean values of n = 3 biological replicates shown as individual dots. For all panels, coculture composition was determined by flow cytometry. OD and fluorescence data were collected using a microplate reader.

As a result, coculture composition was brought closer to a 1:1 ratio as pC-HSL production increased, upregulating anti-STAR production in the YFP strains, and thus sequestrating more STAR to prevent the production of VioB-YFP that destabilises the coculture composition. Another parameter that is interesting to tune is the maximal output of the STAR-based comparator by using different toehold domains. To this end, we used the LLL_1_ and LLL_2_ comparator designs from **Figure 2D** to increase the output of the STAR comparator. As the output of the LLL_1_ comparator is 2-fold higher than that of LLL_2_, we used LLL_2_ for medium-strength STAR expression and LLL_1_ for high STAR expression. Doing so, we observed that as STAR expression increases, the YFP strain is more rapidly outcompeted by the CFP strain as VioB-YFP expression increases (**Figure 5B**, **Figure 5C**). If anti-STAR is not present in high enough concentrations, STAR is not fully sequestered, leading to stabilisation of the coculture at a 2:1 ratio, otherwise the composition stabilises around a 1:1 ratio, that is the same as the initial seeding ratio. The results confirm that tuning the input-output properties of the RNA-based comparator by changing the binding affinity of STAR and anti-STAR or by tuning anti-STAR expression, we can modify the composition of a two-member *E. coli* coculture.

Our multicellular controller has the potential to balance burden and production, thus we investigated how our system influences coculture biomass accumulation and product yields when using VioB as a proxy protein as a proof-of-concept. Looking at case 4 of **Figure 5B** for which both VioB-YFP and VioB-CFP are being expressed, we showed that when no L-arabinose is added, the CFP strain does not outcompete the YFP strain as no protein of interest is being produced (**Figure 5D**). Interestingly, the CL coculture diverges from its equilibrium composition after 6 hours, which can be explained by the difference in burden caused by the different anti-STAR designs from the CFP and YFP strains (**Supplementary Figure 8**). Next, in the presence of L-arabinose only, we observed that the OL coculture ratio is driven out of its equilibrium as the CFP strain quickly outcompetes the YFP strain. The CL coculture however can remain around a 1:1 ratio, keeping a stable coculture composition over time. When the system is induced with both L-arabinose and HSLs, the OL and CL cocultures both stabilise around a 1:1 ratio, demonstrating that the comparator can compensate for the difference in density of the two strains when quorum sensing molecules are present in sufficiently large amounts. When looking at the final density of the cocultures, we observe that when externally inducing the system with L-arabinose, the CL coculture achieves a final density that is 2-fold higher than that of the OL coculture and reaches a similar density (∼0.6) than the non-induced OL (Figure 4C). This increase of biomass accumulation for the CL coculture translates to an 81% increase in total VioB-YFP produced and a 35% increase in total VioB-CFP produced after 24 hours compared to the performance of the OL coculture (**Figure 5E**). By repressing VioB-YFP and VioB-CFP expression to balance the coculture composition, the controller effectively allowed both strains to grow better, thus improving biomass accumulation, which in turn led to higher production yields of the protein of interest. When both L-arabinose and HSLs were externally added into the medium, VioB-CFP and VioB-YFP expression were inhibited, resulting in little VioB-CFP and VioB-YFP being produced. Finally, we tested whether the STAR-based controller could stabilise population composition when the initial starting ratio was different from 1:1 (**Figure 5F**). For this, we decided to inoculate our OL and CL cocultures over a range of seeding ratios. After 12 hours, YFP Strain fraction in all OL cocultures had dropped by at least 25% and as much as 55%. For the CL coculture however, the YFP Strain fractions appear to stabilise around their initial seeding ratio, not deviating from it more than 12%. This highlights the ability of our CL system to robustly stabilise coculture composition around the initial seeding ratio of the coculture.

## Discussion

Here, we present a multicellular control strategy using molecular sequestration to stabilise the composition of an engineered microbial consortia. *E. coli* was engineered to express three modules for the bottom-up assembly of microbial consortia: (a) a quorum-sensing-based communication module to obtain information about the cocultured strains densities, (b) an RNA-based comparator module to compare the population density of two strains grown in a coculture, and (c) a growth module, which modulates expression of a growth regulator to tune cellular growth and thereby the desired coculture composition. The RNA-based comparator, the first of its kind, can modulate growth rates via burden regulation of either a single protein or a metabolic pathway, but also through essential gene knockdown using CRISPRi. As a result, the genetic circuit, split across the two microbial species, is able to stabilise population composition when their respective protein productions impose a different burden on each host. We used the burden caused by the expression of heterologous genes of interest (GOI) to control the growth of the two cocultured bacterial strains. As such, the growth control mechanism does not rely on consuming additional cellular resources to produce mutagenic toxins that kill the hosts rather that slowing down their growth. Our multicellular gene circuit could stabilise population composition over time compared to a coculture deprived on the controller circuit. In addition, by modulating the burdensome expression of the GOIs, we found that the multicellular controller improved the total production yields by 81% in the slowest growing strain and by 35% in the fastest growing strain. We identified several parameters that can be used to tune community composition: quorum sensing production rate, transcription rate of the GOI, the transcription rate of the reference signal promoter and the binding affinity of the STAR to its target and to its antisense specie, anti-STAR. As such, the platform we developed could provide a means to balance heterologous expression burden and production, leading to better biomass accumulation and production yield in engineered microbial communities. It paves the way to the development of host-aware complex multicelullar systems for synthetic biology. Overall, this study provides valuable insights to shift the current efforts to improve product yields by solely optimising individuals strains onto optimising consortia of strains working in concert as specialised entities for a complex and common goal. Further studies investigating the control of both production and coculture composition over time, will contribute to assess the importance of dynamic multicellular feedback systems for the improvement of yields and functions in bioreactors and spatially separated environments. If successful, these host-aware and multicellular control strategies could also provide an important method to improve productivity, especially while tackling the issue of competitive exclusion in engineered microbial consortia.

## Material and Methods

### Bacterial strains and plasmids

DH10B (K-12 F- λ- araD139 Δ(araA-leu)7697 Δ(lac)X74 galE15 galK16 galU hsdR2 relA rpsL150(StrR) spoT1 deoR ϕ80dlacZΔM15 endA1 nupG recA1 e14- mcrA Δ(mrr hsdRMS mcrBC)) were obtained from the National BioResource Project Japan. BW25113 (K-12 acI+rrnBT14 ΔlacZWJ16 hsdR514 ΔaraBADAH33 ΔrhaBADLD78 rph-1 Δ(araB–D)567 Δ(rhaD–B)568 ΔlacZ4787(::rrnB-3) hsdR514 rph-1) and JW5807 (BW25113 ΔleuB) were obtained from the Keio Collection. pC-HSL and C6-HSL producing strains were built by integrating *luxI* and the rpa operon (*TAL, 4CL2nt, rpaI*) into the λ phage attachment locus of DH10B by CRIM integration ^73^. We also inserted *mScarlet-I* and *sfGFP* as fluorescence markers for tracking coculture ratios. Genes were cloned into the CRIM integration vector plasmids pAH63 and propagated in *pir-116* electrocompetent *E. coli* cells (Lucigen). Integration, curation and validation of the integrated strains were done following the protocol from Haldimann et al. ^73^. The bacterial strains used in this work are detailed in Supplementary Table 1. The plasmids created in this study are detailed in Supplementary Table 2. All plasmid maps will be made available online on Zenodo. Polymerase chain reactions, Gibson and Golden Gate assemblies were used to build those plasmids. All plasmid sequences were verified using Sanger sequencing.

### Time-course fluorescence assays

Time-course experiments were performed in clear flat-bottom 96-well plates (Costar) with three biological replicates using a Tecan Spark microplate reader. Cells transformed with the constructs of interest and control plasmids were inoculated with 5 mL of M9 (with casaminoacids unless otherwise stated) supplemented with the appropriate antibiotics and grown overnight at 37°C with aeration in a shaking incubator. In the morning, cultures were diluted by 1:4 with fresh M9 in 1 cm cuvettes to measure the OD700 of each sample in the spectrophometer (WPA Biowave II). Each sample was diluted to OD700 0.01 in 2 mL of fresh M9 supplemented with the appropriate antibiotics (kanamycin: 50 μg/mL; ampicillin: 100 μg/mL; spectinomycin: 50 μg/mL; streptomycin: 100 μg/mL; chloramphenicol: 34 μg/mL; tetracycline: 10 μg/mL). For dBroccoli measurements, DFHBI-1T dye (Bio-Techne) was added to a final concentration of 100 μM unless otherwise stated. 200 μL of each sample was then transferred into a sterile 96-well plate and covered with a Breath-Easy membrane (Sigma). The plate was placed into a microplate reader and incubated at 37°C for 1 h (Tecan Spark: Double orbital shaking, 1.5 mm amplitude). Measurements of OD700 and fluorescence (sfCFP: 430(20) nm ex./465(35) nm em.; sfYFP/sfGFP/dBroccoli: 485(20) nm em./535(25) nm em.; mScarlet-I/mRFP1/mCherry/mKate: 560(20) nm ex./ 620(20) nm em.) were taken every 15 minutes. 1 hour into the incubation, we briefly removed the microplate from the plate-reader, carefully removed the Breath-Easy membrane and added the appropriate inducers to each well in the appropriate concentrations. We then covered the microplate with a new Breath-Easy Membrane and introduced it back into the plate-reader and set this time point as "time 0" by creating a “new plate” in the experiment. OD700 and fluorescence (Table 7.12) were taken every 15 minutes. Cells were grown for 6 to 24 hours depending on the experiment. Data were exported into an Excel Spreadsheet and analysed using MATLAB.

OD and fluorescence raw data were first subtracted with the mean of M9 media well replicates over time. Data were then smoothed using MATLAB smoothingspline function (smoothing parameter: 0.8648426188005848). Growth rate and fluorescence production rate per cell was calculated as described in Ceroni et al ^50^: Growth rate at t2 = (ln(OD(t3)) – ln(OD(t1)))/(t3 – t1), GFP production rate at t2 = ((Total GFP(t3) – Total GFP(t1))/(t3 – t1))/OD(t2), and mCherry production rate at t2 = ((Total mCherry(t3) – Total mCherry(t1))/ (t3 – t1))/OD(t2), where t1 corresponds to 0 min after induction, t2 to 15 min after induction and t3 = 30 min after induction. Mean values and standard deviations were calculated from the three biological replicates of each sample. For figures representing input-output response curves, we used the nlinfit and fitnlm MATLAB functions to fit a four-parameter logistic regression model (4PL model) to the data and determine the R-squared and p-value of the model: a +((b-a)/(1+(x/c)^d)).

### Flow cytometry assays

Flow cytometry was used to measure coculture composition by counting the number of red fluorescent cells and non-fluorescent cells. Cell fluorescence was measured in the Attune NxT (Thermo Scientific) flow cytometer using the following parameters: FSC 660 V, SSC 500 V, violet laser VL1 (405 nm ex./440(50) nm em.) 420 V, blue laser BL1 (488 nm ex./530(30) nm em.) 450 V, yellow laser YL2 (561 nm ex./620(15) nm em.) 560 V. The following threshold were used: AND FSC 0.5×1000, AND SSC 4×1000. 5 mixing cycles were used between each sample. 10,000 cells were collected for each sample and data were analysed using FlowJo and plotted with MATLAB. In FlowJo, we first gated E. coli cells from dust and cell debris by plotting SSC-A vs FSC-A. Then we gated the E. coli population to find the singlets population by plotting FSC-H vs FSC-A. Finally, the red and non-red singlets populations were determined by plotting BL1-H vs YL2-H. For graphs representing fluorescence intensity, points represent the mean of the median fluorescence of three biological samples. Deactivation percentage of the STAR-based comparator was calculated using the following formula: deactivation % = ((F_STAR_ - F_STAR,anti-STAR_)/(F_STAR_ - F_neg._))*100, Where F represents average median fluorescence obtained by flow cytometre of three biological samples. F_STAR_ is the fluorescence of the comparator when STAR is expressed, F_STAR,antiSTAR_ is the fluorescence of the comparator when both STAR and anti-STAR are expressed, and F_neg._ is the fluorescence of the comparator when neither STAR nor anti-STAR are expressed.

### Quorum sensing sender-receiver assay

Sender strains and receiver strains were inoculated in 1 mL of rich M9 medium supplemented with the appropriate antibiotics in a 2 mL deep-well 96-well plate (VWR), covered with a “Breathe Easier” membrane (Sigma), and incubated overnight at 30°C in a plate shaker incubator (Infors HT Multitron) shaking at 700 rpm overnight. In the morning, sender strains were diluted to 1:4 in fresh M9 medium in a 1 cm cuvette and OD700 was measured in the spectrophotometer. Cells were diluted to OD700 0.05 in 1 mL of fresh M9 medium with antibiotics in a new 2 mL deep-well 96-well plate, covered with a “Breathe Easier” membrane (Sigma), and incubated at 700 rpm, 30°C in the plate-shaker incubator. Every hour for 6 hours, the deep-well plate was taken out of the incubator and new wells were inoculated with 1 mL of OD700 0.05 of each culture. Receiver strains were diluted 1:200 in fresh M9 with antibiotics in a new deep-well plate and incubated at 700 rpm, 30°C in the plate-shaker incubator. At 6 hours, 250 μL of each sample from the sender strains deep-well plate was transferred to 1 cm cuvette to measure OD700 of each sender strain culture. The sender strains deep-well plate was then centrifuged at 4000 rpm for 5 minutes (Centrifuge Eppendorf 5810R) and the supernatants were transferred by pipetting in a new 2 mL deep-well plate, diluted 1:1 with fresh M9 with antibiotics and the pellets were discarded. The OD700 of the receiver strains were measured in the spectrophotometer by diluting 1:2 with fresh M9 in 1 cm cuvettes. Receiver strains were then diluted to OD700 0.01 in 1 mL of the diluted sender strains’ supernatants. 200 μL of each sample was transferred to a clear flat-bottom 96-well plate and inducers (and dye if dBroccoli was expressed) were immediately added to the appropriate wells. The microplate was covered with a “Breathe-Easy” membrane (Sigma) and incubated in the Tecan Spark for 12 hours. OD and fluorescence were measured every 15 minutes as previously described in the above “plate-reader assay” section.

### Coculture assay

Cells carrying the open-loop and closed-loop versions of the composition controller circuits were grown as monocultures in 1 mL of rich M9 medium supplemented with the appropriate antibiotics in 2 mL 96-well deep-well plates (VWR) at 30°C in a plate shaker incubator (Infors HT Multitron) shaking at 700 rpm overnight. In the morning, cells were diluted 1:4 with fresh rich M9 medium and transferred to 1 cm cuvettes to measure OD700 in the spectrophotometer. Subsequently, each sample was diluted to OD_700_ 0.01 in a volume of 2 mL (adjusted according the experiment’s needs). In a transparent flat-bottom 96-well plate (Costar) which we will refer to as the "experiment 96-well plate", 100 μL of each monoculture that will compose the two-strain coculture are mixed in the appropriate wells (total volume of 200 μL in each well). Cocultures and monoculture controls were immediately induced with the appropriate inducers. The 96-well plate was covered with its plastic lid, inserted into a Tecan Spark plate-reader and incubated at 37°C with shaking. OD700 and fluorescence was measured every 15 minutes. Every 2 hours for 12 hours, we paused the TECAN Magellan program running the plate-reader experiment and, under sterile condition, we transferred 1-6 μL of cells (0-2 hours: 6 μL; 4-6 hours: 3 μL; 8-10 hours: 1 μL, 12-24 hours: 0.5 μL) into a clear round-bottom 96-well plate (Costar) pre-filled with 200 μL of 1X PBS (Sigma) supplemented with tetracycline (10 μg/mL). We call this plate the "flow plate" as it will be used to measure cell fluorescence in the flow cytometer to determine coculture composition. The "experiment 96-well plate" was then covered with its lid and placed back into the plate-reader where the TECAN Magellan program was resumed. The "flow plate" was then immediately stored on ice in the fridge at 4°C. When the "flow plate" was entirely filled with coculture samples diluted in PBS, it was then run in the flow cytometer as previously described in the above “flow cytometry assay” section.

### β-Carotene assay

The protocol was adapted from Borkowski et al ^72^. For quantification of β-Carotene production, cells carrying the β-Carotene producing plasmids and controls were grown in 5 mL of rich M9 media in 15 mL culture tube overnight at 37°C in shaking conditions. In the morning, cells were diluted 1:4 in fresh M9 media and 1 mL of diluted cultures were transferred to 1 cm cuvettes to measure OD700 in the spectrophotometer. Each sample was diluted to OD700 0.01 in a volume of 5 mL in a sterile 15 mL culture tube and induced with the appropriate inducers. After 6 hours of growth at 37°C in the shaking incubator, 0.5 mL of each culture was diluted 1:1 with fresh rich M9 media and its OD700 was measured in the spectrophotometer. The remaining 4.5 mL of each culture were spun down at 4000 rpm for 10 minutes (Centrifuge Eppendorf 5810R) and remaining supernatant was discarded by pipetting. Pellets were resuspended in 300 μL of acetone in 1.5 mL Eppendorf tubes, homogenised by vortexing for 10 minutes and incubated at 55°C (Eppendorf ThermoMixer C) for 15 minutes. Tubes were centrifuged for 1 minute at 10,000 rpm (Thermo Scientific Heraeus Fresco 17 Centrifuge). 100 μL of the supernatants were collected by pipetting and transferred to a new 1.5 mL Eppendorf tube. A volume of 100 μL of water was added to the 100 μL of supernatants in each tube and mixed by pipetting. The total 200 μL were then transferred to a clear flat-bottom 96-well plate (Costar) and OD450 of the microplate was measured in the plate-reader (Tecan Spark). To compare production of β-Carotene between the different samples, OD450 of each sample was divided by its corresponding OD700. We note that β-Carotene does not absorb at OD700.

## Acknowledgments

Figures were created using BioRender.com. We thank J. Lucks and M. Verosloff for plasmids pJBL4826, pJBL4882, pJBL5938, pJBL5939, pJBL5945, pJBL5946, pJBL6063. G.-B.S. gratefully acknowledges support from the U.K. Royal Academy of Engineering through the Royal Academy of Engineering Chair in Emerging Technologies for Engineering Biology [CiET 1819\5] and of the H2020 FET-OPEN project 766840 (COSY-BIO). R.L.A. received funding from BBSRC (BB/R01602X/1), (19-ERACoBioTech- 33 SyCoLim BB/T011408/1) and (BB/T013176/1), British Council 527429894, Newton Advanced Fellowship (NAF\R1\201187), Yeast4Bio Cost Action 18229, European Research Council (ERC) (DEUSBIO - 949080) and the Bio-based Industries Joint (PERFECOAT- 101022370) under the European Union’s Horizon 2020 research and innovation programme

## Author contributions

A.B., R.L.A., G.-B.S designed the study; A.B. performed the experiments and collected the data; A.B. analysed the data; A.B., H.M. and G.-B.S. developed the mathematical model; A.B., H.M., R.L.A. and G.-B.S discussed the results, wrote and edited the paper.

