## Supplementary Materials for "Host-aware RNA-based control of synthetic microbial consortia"

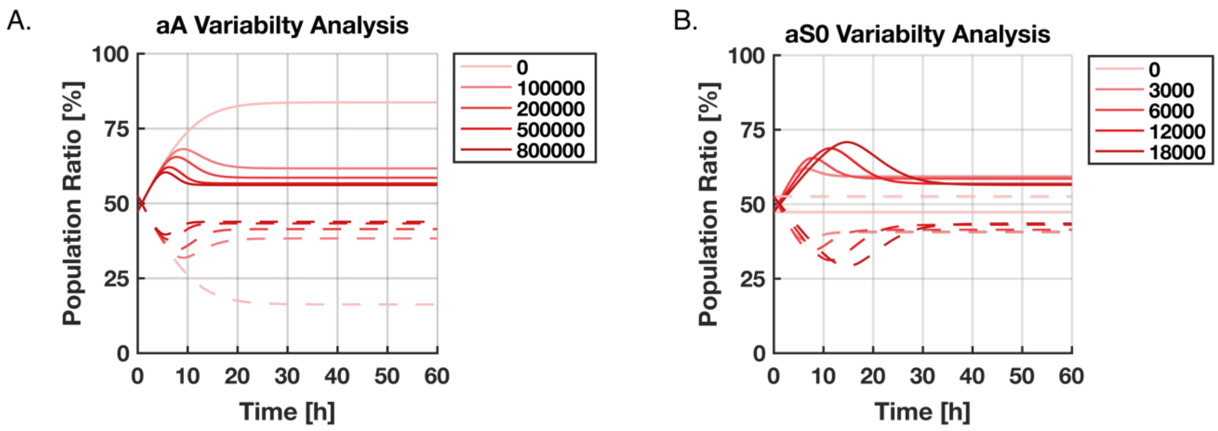

**Supplementary Figure 1:** **Simulations of the microbial composition.** Plots are simulations of the mathematical model from **Supplementary Note 1** that describes the RNA-based comparator that controls growth rate through the burden caused by expressing VioB. (A) Tuning the anti-STAR rate of transcription. The rate of STAR transcription is set as aS0 = 8,000. When aA is set to 0, no anti-STAR is expressed in the cell, and as a result the growth controller is inactive. As the values of aA increase, more anti-STAR is expressed in the cells. As a result, more STAR is sequestered which prevents the expression of the burdensome proteins that slow down cellular growth rate. (B) Tuning the STAR transcription rate. The rate of anti-STAR transcription is set as aA = 200,000. When aS0 is set to 0, no burdensome protein is expressed in either strain 1 or strain 2 of the microbial consortia, thus both strains grow at the same rate. However, as the transcription rate of STAR, aA, increases, each strain expresses more of their respective burdensome protein. As the burdensome proteins have different burden, they impact growth rate differently and the population ratio diverges from the initial inoculation ratio (50:50). The divergence from the initial inoculation rate is more pronounced as the rate of STAR increases as it, in turn, increases production of the burdensome proteins. Simulations were performed in the MATLAB Simbiology toolbox.

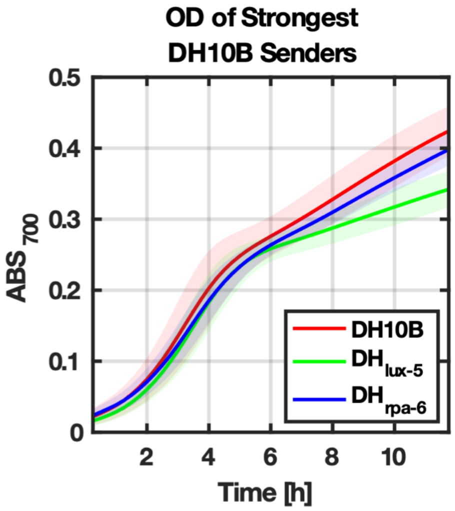

**<u>Supplementary Figure 2:</u>** **Growth of quorum-sensing-producer strains.** OD of the strongest C6-HSL-producing DH10B strain ($Lux_5$) and the strongest pC-HSL-producing DH10B strain ($Rpa_6$) compared to the OD of the wild-type DH10B strain (Supplementary Table 1). OD measurements were collected in time-course plate-reader assay. Data represent the mean values of n = 3 biological replicates ± s.d.

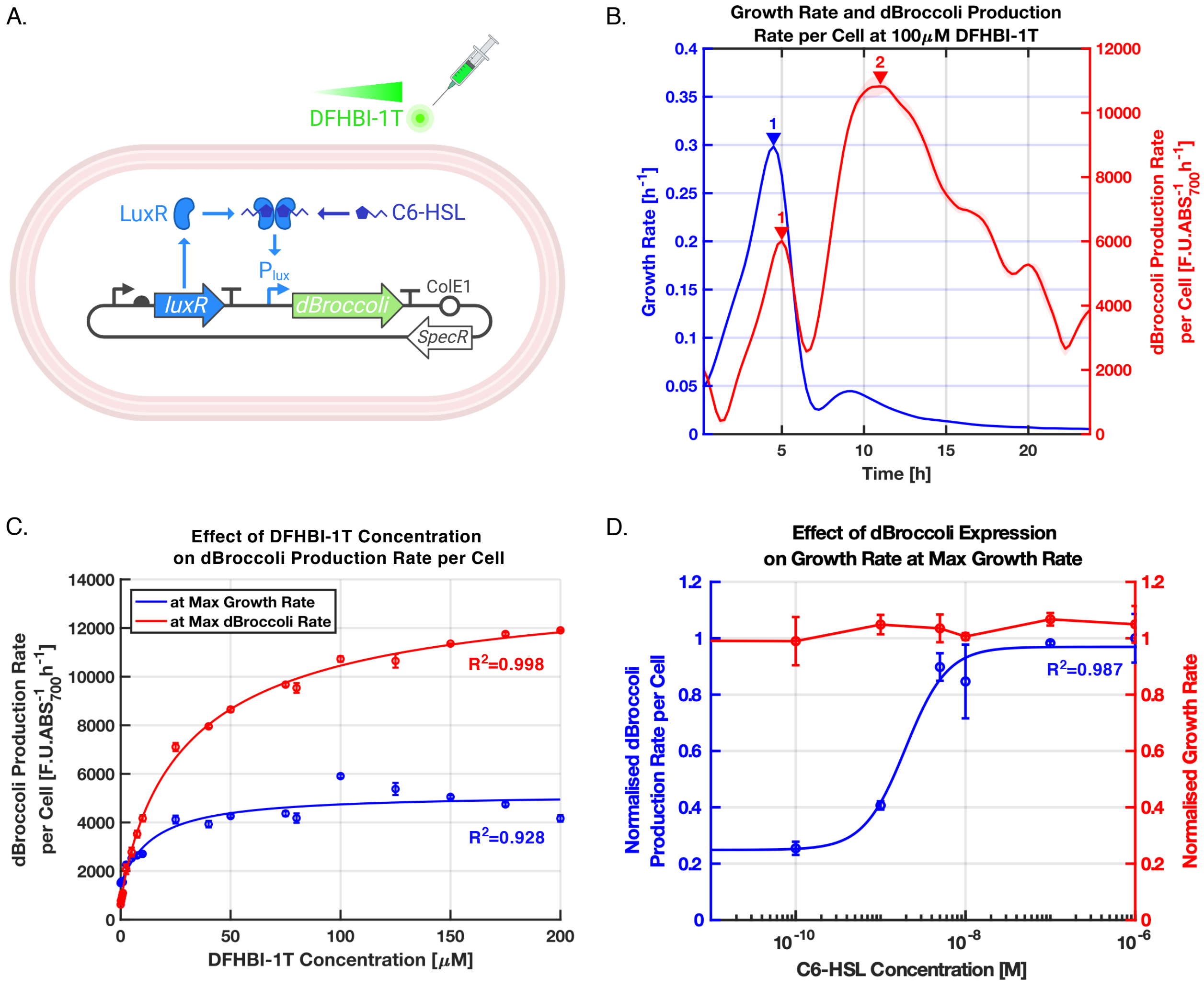

**<u>Supplementary Figure 3:</u> Effect of DFHBI-1T dye concentration on dBroccoli signal strength and host growth rate.** (A) Different concentrations of DFHBI-1T dye are tested to study the impact of DFHBI-1T concentration on dBroccoli signal strength. dBroccoli is expressed under the LLL quorum sensing system, with $10^{-7}$M of C6-HSL in DH10B carrying the pAB420 plasmid (Supplementary Table 3). (B) dBroccoli production rate per cell has two maxima: one coinciding with the maximum growth rate of the cells during exponential phase, and the other when the culture is in stationary phase. (C) Effect of DFHBI-1T dye concentration on dBroccoli production rate per cell at both dBroccoli production maxima. Cells are induced with $10^{-7}$M of C6-HSL. (D) Effect of dBroccoli expression on growth rate at the time of maximum growth rate. Cells are mixed with 100 µM of DFHBI-1T dye. dBroccoli fluorescence was monitored in a time-course plate-reader assay. Curves were fitted using MATLAB four-parameter nonlinear regression fit. Data represent the mean values of n = 3 biological replicates ± s.d.

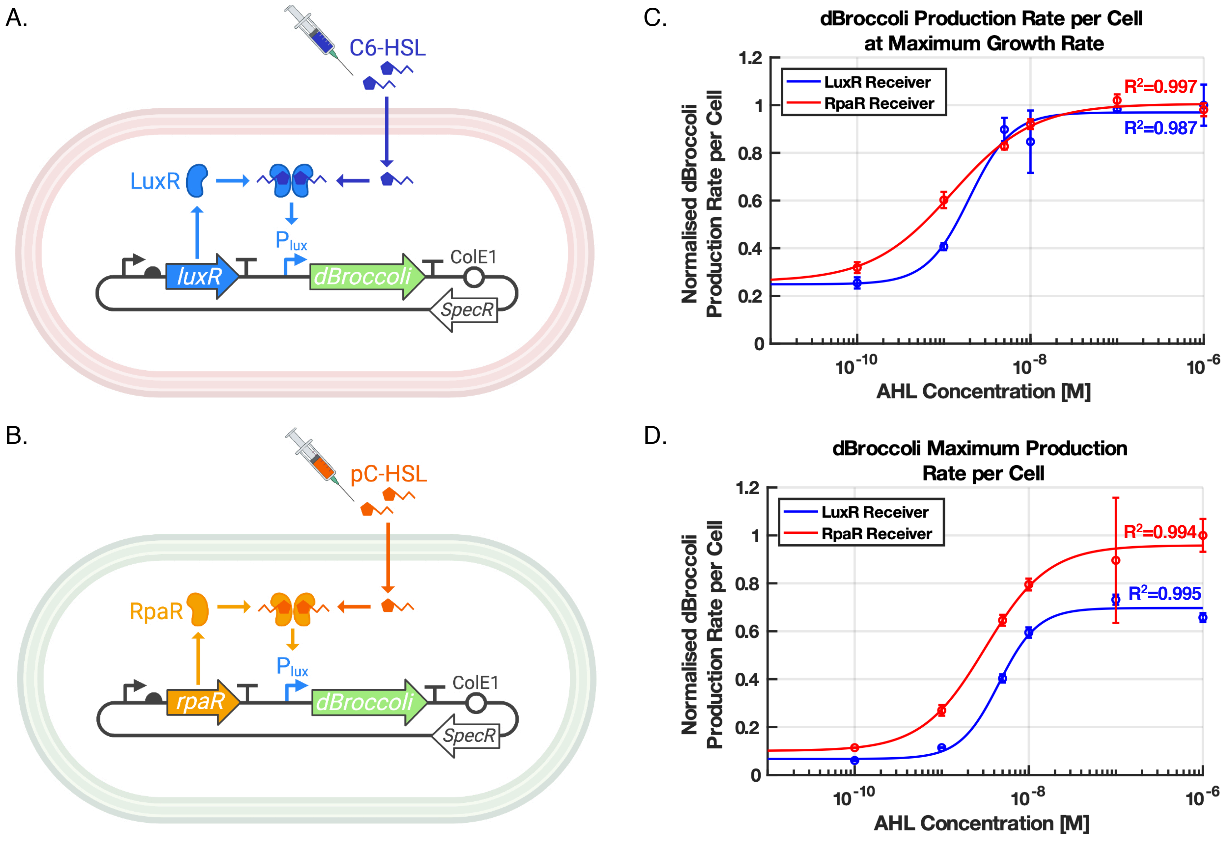

**<u>Supplementary Figure 4:</u>** **Response of the lux and rpa systems to HSL gradients.** (A) The LuxR Receiver circuit is composed of the LLL driving the expression of dBroccoli in DH10B carrying the pAB420 plasmid (Supplementary Table 3). (B) The RpaR Receiver is composed of the LRR system driving the expression of dBroccoli in DH10B carrying the pAB421 plasmid (Supplementary Table 3). (C) dBroccoli production rate per cell at the time of maximum growth rate, for different concentrations of C6-HSL and pC-HSL added to the LuxR Receiver and the RpaR Receiver respectively. (D) dBroccoli production rate per cell at the time of maximum dBroccoli production, for different concentrations of C6-HSL and pC-HSL added to the LuxR Receiver and the RpaR Receiver respectively. dBroccoli fluorescence was monitored in a time-course plate-reader assay. Curves were fitted using MATLAB four-parameter nonlinear regression fit. Data represent the mean values of n = 3 biological replicates ± s.d.

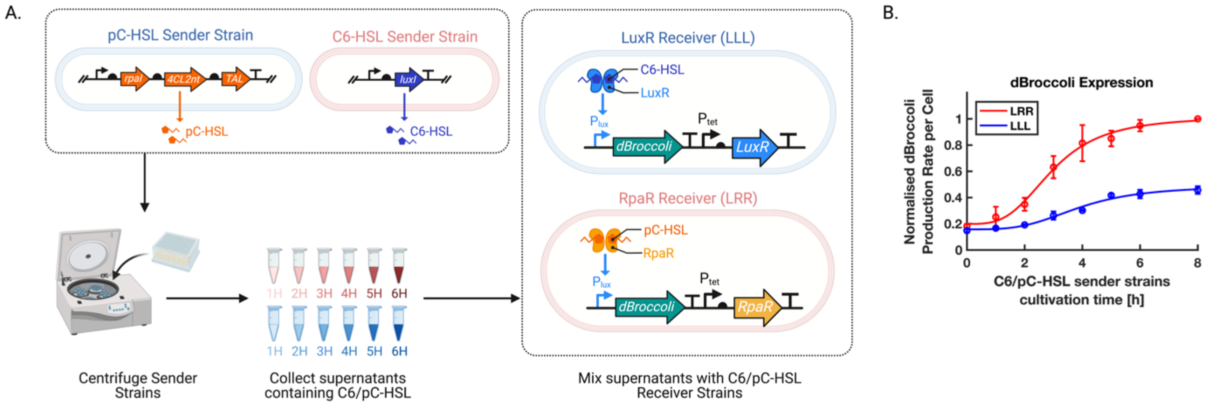

**<u>Supplementary Figure 5:</u> Quorum sensing production by the pC-HSL and C6-HSL producer strains.** (A) Sender strains producing either C6-HSL or pC-HSL, respectively, are incubated for a period of 1 to 8 hours before being centrifuged for their supernatants to be collected every hour and mixed with the appropriate C6-HSL and pC-HSL receiver strains. The response of the receiver strains DH10B carrying either pAB420 or pAB421 (Supplementary Table 3) to the HSL produced by the sender strains is monitored using a plate-reader assay detecting dBroccoli fluorescence. (B) C6-HSL and pC-HSL production by $Lux_1$ and $Rpa_5$ sender strains respectively (Supplementary Table 1) increases over time. C6-HSL and pC-HSL are detected by the LLL and LRR receiver strains cultivated in the supernatants of the $Lux_1$ and $Rpa_5$ stains. dBroccoli fluorescence was measured in a timecourse plate-reader assay. Data represent the mean values of n = 3 biological replicates ± s.d.

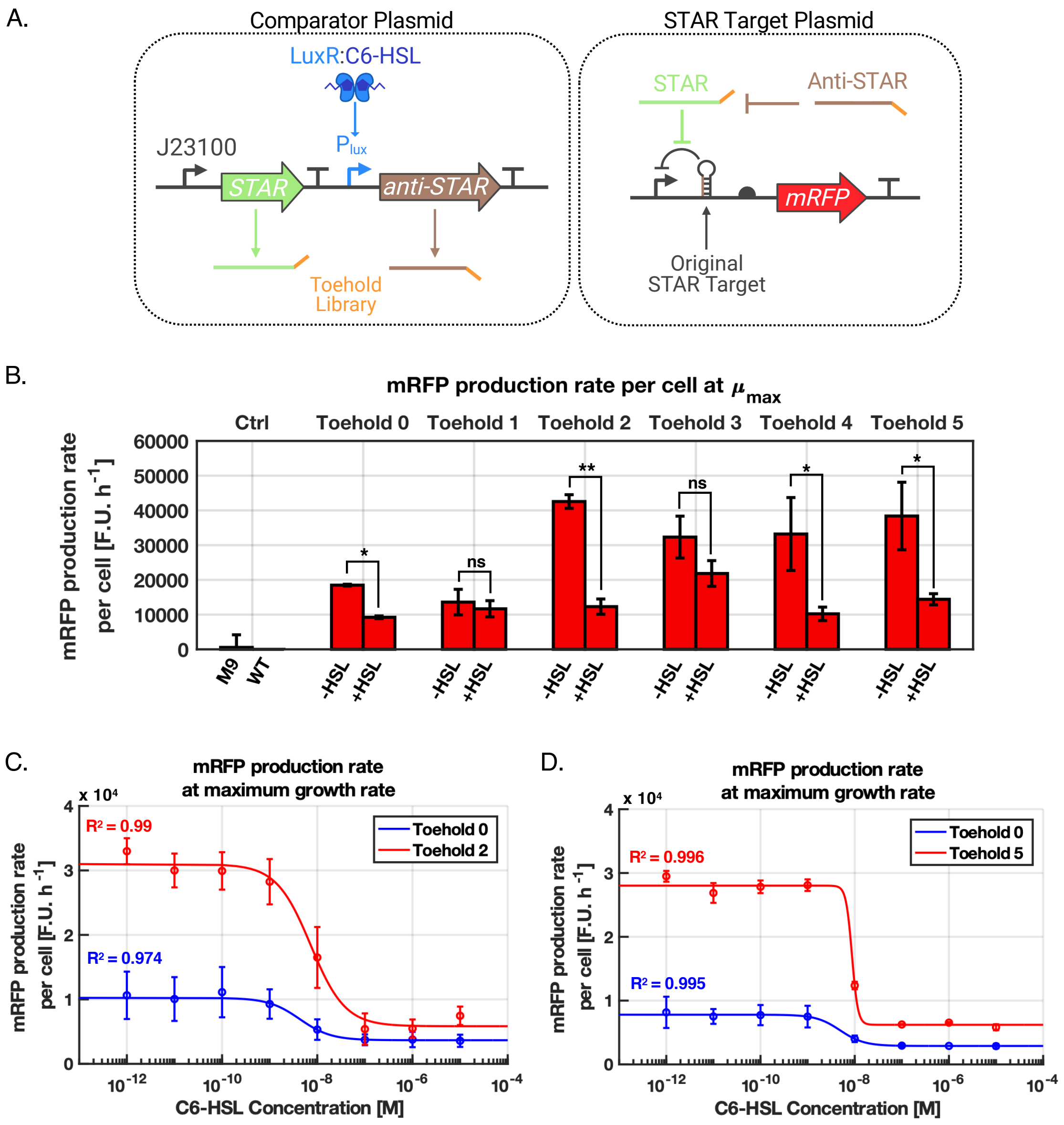

**Supplementary Figure 6:** **STAR-based comparator toehold library.** (A) Diagram of the J23100-pLux STAR-based comparator with the toehold library. The target plasmid pJBL5939 (Supplementary Table 3) carries a single STAR target driving expression of mRFP. (B) Bar graph representing the mRFP production rate of the strains carrying the comparators with the different toeholds. Plasmids pAB161, pAB232, pAB233, pAB234, pAB235, pAB236 are the plasmids carrying the comparator designs with toeholds 0 to 5 respectively (Supplementary Table 3). "-HSL" represents a concentration of 0M while "+HSL" represents a concentration of 10−7M of C6-HSL. (C) Toehold 0 and Toehold 2 comparators' normalised mRFP production rate per cell as a function of C6-HSL concentration. (D) Toehold 0 and Toehold 5 comparators' normalised mRFP production rate per cell as a function of C6-HSL concentration. Experiments were carried in BW25113. Data represent the mean values of n = 3 biological replicates ± s.d. Curves were fitted using MATLAB four-parameter nonlinear regression fit. Statistically significant differences were determined using two-tailed Student's t-test (**** represents p<0.0001, *** represents p<0.001, ** represents p<0.01, * represents p<0.1, ns represents not significant).

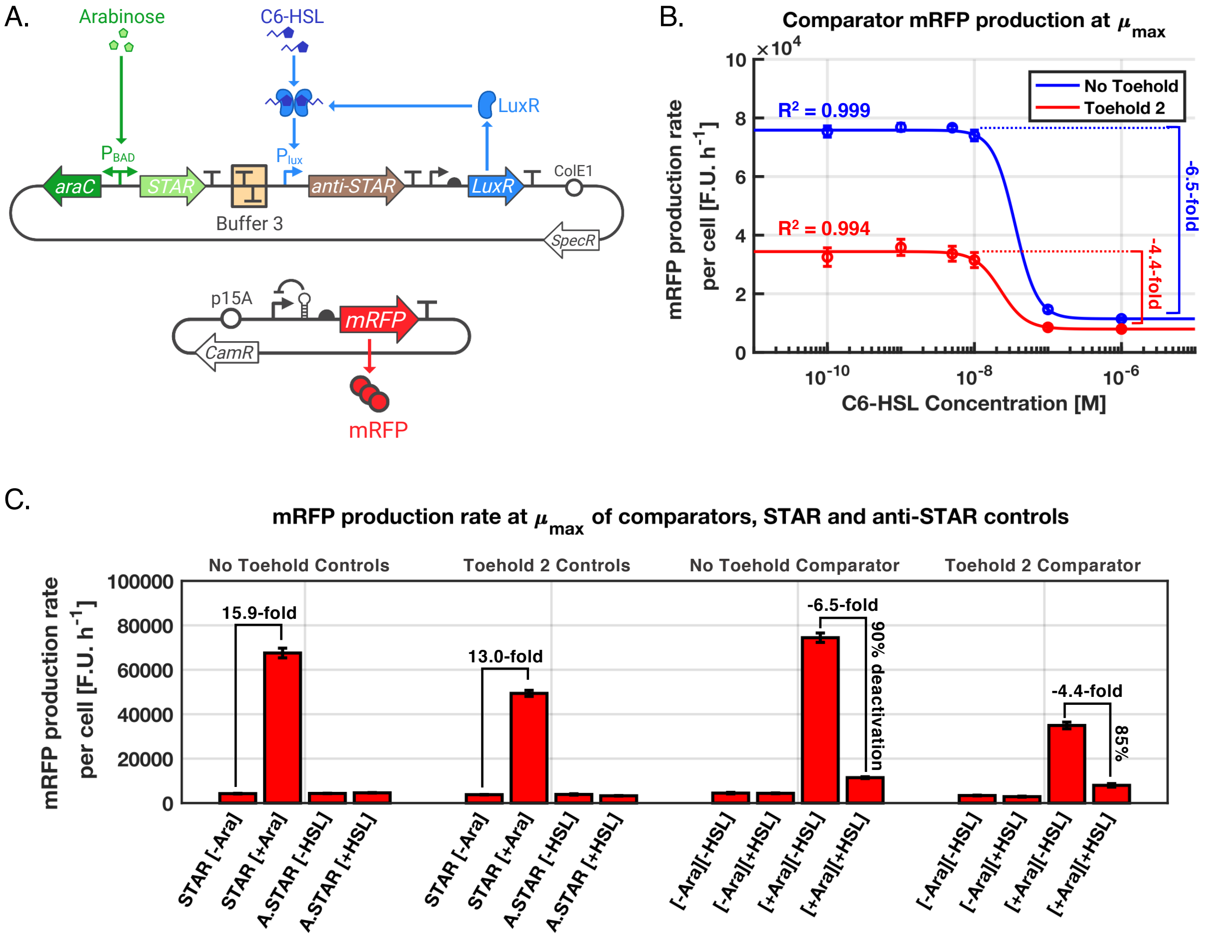

<u>**Supplementary Figure 7:**</u> **Characterisation of the STAR-based comparator with and without toehold.** (A) Diagram of the circuit used to compare the STAR-based comparator with and without toehold. DH10B carrying pJBL5939 and pAB300 was used to test the comparator design without a toehold, and DH10B carrying pJBL5939 and pAB317 was used to test the comparator design with Toehold 2 (Supplementary Table 3). (B) No Toehold and Toehold 2 comparators' VioB-mCherry production rate per cell at the time of maximum growth rate and as a function of C6-HSL concentration. (C) Bar graph of mRFP production rate per cell. The STAR controls only carried STAR with or without toehold on the comparator plasmid and were induced by L-arabinose. The "A.STAR" controls only carried anti-STAR with or without toehold on the comparator plasmid and were induced by C6-HSL. Two concentrations of L-arabinose were used for induction: "[-Ara]" for 0% and "[+Ara]" for 0.2%. 2 concentrations of C6-HSL were used for induction: "[-HSL]" for 0M and "[+HSL]" for $10^{-6}$M. The deactivation percentage of the comparators was calculated as $(mRFP_{[+Ara][-HSL]} - mRFP_{[+Ara][+HSL]})/(mRFP_{[+Ara][-HSL]} - mRFP_{[-Ara][-HSL]})$, where mRFP is the mRFP production rate per cell. Data represent the mean values of n = 3 biological replicates ± s.d. Curves were fitted using MATLAB four-parameter nonlinear regression fit.

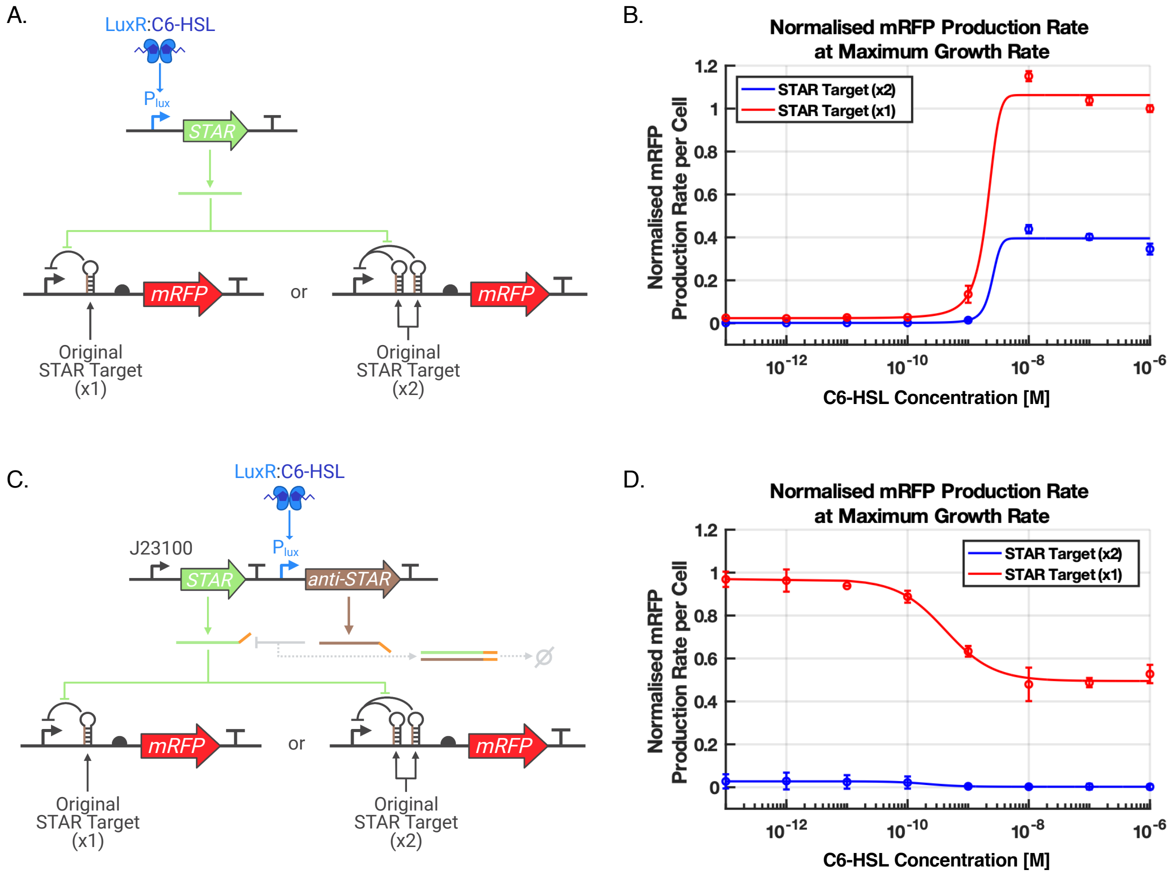

__Supplementary Figure 8:__ **Effect of STAR-double-target.** (A) Diagram of circuit used to characterise the effect of the STAR-double-target on mRFP expression. BW25113was transformed with the pAB127 plasmid expressing STAR and with either pJBL5939 or pAB262 carrying 1 or 2 STAR Target sites respectively (Supplementary Table 3). (B) Normalised mRFP production rate per cell as a function of C6-HSL concentration when the gene of interest (mRFP1) is controlled by either a single-STAR target or a double-STAR target. (C) Diagram of circuit used to characterise the effect of the STAR-double-target on the behaviour of the STAR-based comparator. BW25113 was transformed with the pAB161 plasmid expressing the Toehold 0 comparator and with either pJBL5939 or pAB262 carrying 1 or 2 STAR Target sites respectively (Supplementary Table 3). (D) Normalised mRFP production rate per cell as a function of C6-HSL concentration when the STAR-based comparator's gene of interest (mRFP1) is controlled by either a single-STAR target or a double-STAR target. Data represent the mean values of n = 3 biological replicates ± s.d. Curves were fitted using MATLAB four-parameter nonlinear regression fit.

A. B.

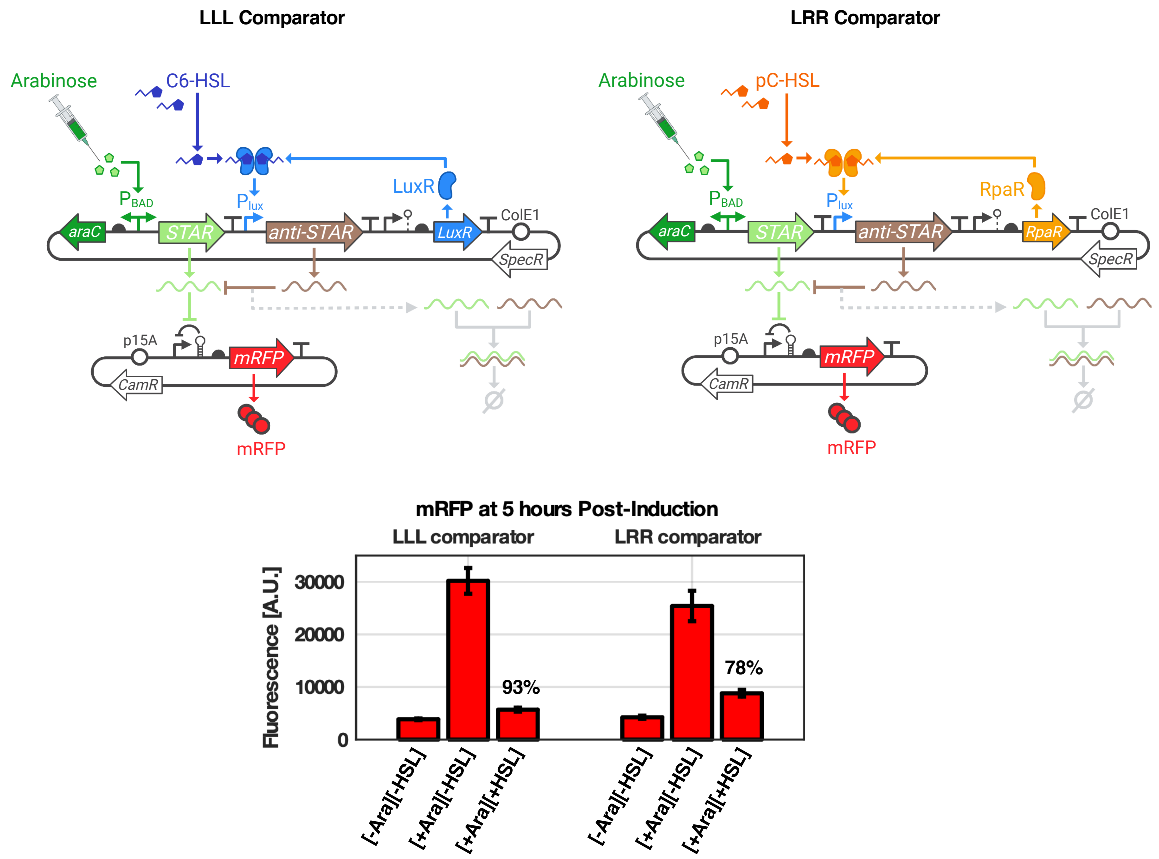

**Supplementary Figure 9:** **Deactivation of the LLL and LRR STAR-based comparator.** (A) Design of the LLL comparator (DH10B carrying the pJBL5939 and pAB300) and LRR comparator (DH10B carrying the pJBL5939 and pAB401, Supplementary Table 3). (B) The deactivation percentage is calculated for the LLL and LRR STAR-based comparators. The comparator is externally induced with three conditions: (1) "[-Ara][-HSL]" represents induction with 0% of L-arabinose and 0 M of C6-HSL or pC-HSL for the LLL or LRR comparator respectively; (2) "[+Ara][-HSL]" represents induction with 0.2% of L-arabinose and 0 M of C6-HSL or pC-HSL for the LLL or LRR comparator respectively; (3) "[+Ara][+HSL]" represents induction with 0.2% of L-arabinose and $10^{-7}$ M of C6-HSL or pC-HSL for the LLL or LRR comparator, respectively. The deactivation percentage is calculated as $((F_{STAR} - F_{STAR,anti-STAR})/(F_{STAR} - F_{neg.})) \times 100\%$, where $F_{STAR}$ corresponds to the fluorescence of the comparator when only STAR is expressed ("[+Ara][-HSL]"), $F_{STAR,anti-STAR}$ corresponds to the fluorescence of the comparator when both STAR and anti-STAR are expressed ("[+Ara][+HSL]"), and $F_{neg.}$ corresponds to the fluorescence of the comparator when neither STAR nor anti-STAR are expressed ("[-Ara][-HSL]"). Fluorescence was measured through flow cytometry. Data represent the mean and standard deviation of 3 biological replicates.

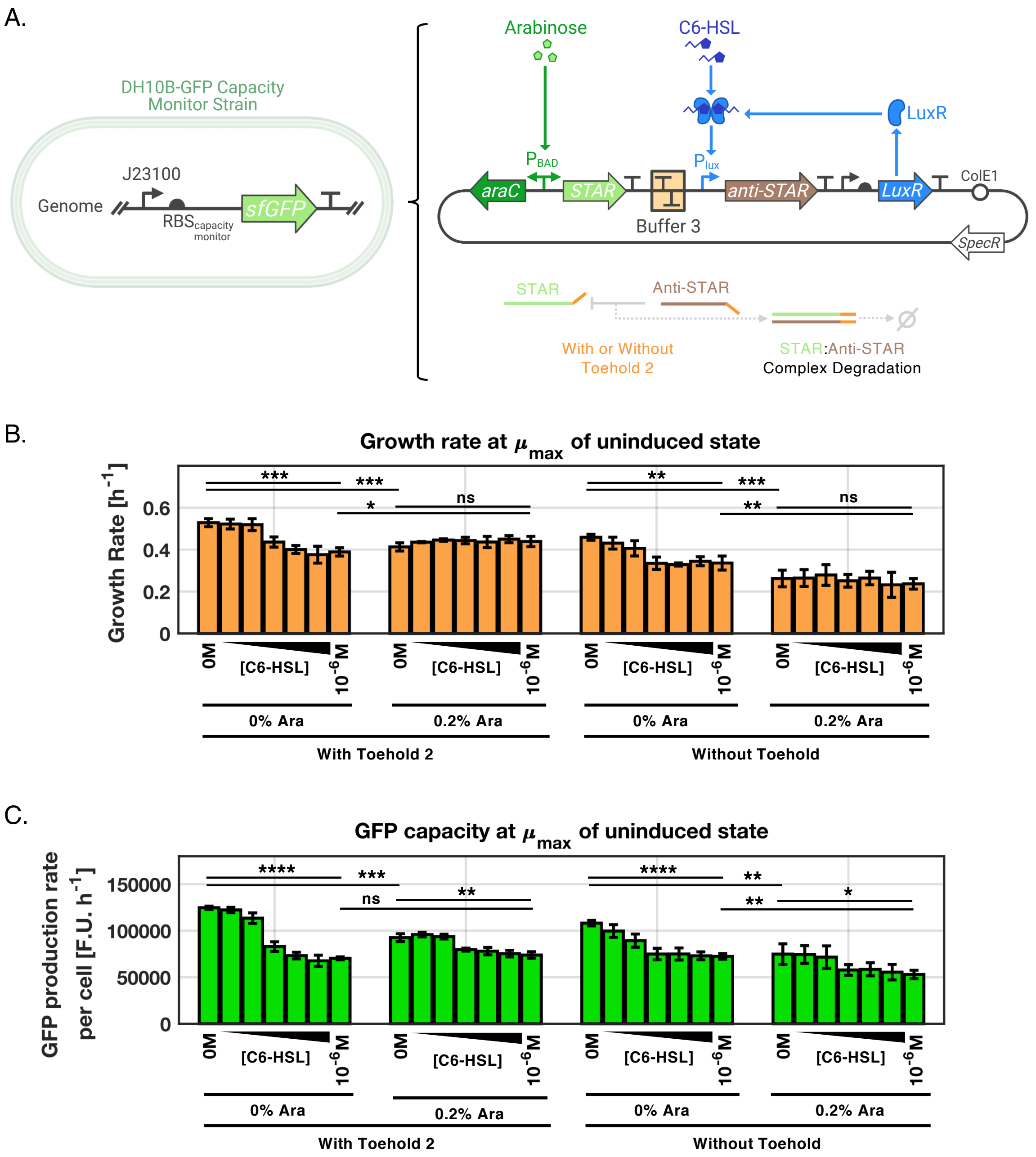

**Supplementary Figure 10:** **Burden characterisation of the STAR-based comparator with and without toehold.** (A) Diagram of the comparator plasmid tested in the DH10B-GFP strain carrying the GFP capacity monitor to assess the burden caused by expression of STAR and anti-STAR. STAR expression is enabled by L-arabinose, while anti-STAR expression is enabled by C6-HSL. DH10B-GFP carrying pAB300 was used to study the impact of expression the comparator with Toehold 2 and DH10B-GFP carrying pAB317 was used to study the impact of expression the comparator without toehold (Supplementary Table 3). (B) Growth rate of the comparator plasmids with and without toehold. (C) GFP capacity is the GFP production rate per cell of the DH10B-GFP capacity monitor strain transformed the comparator plasmids with and without toehold. Two concentrations of L-arabinose were used: 0% and 0.2%. The concentrations of C6-HSL used were: 0 M, $10^{-10}$ M, $10^{-9}$ M, $5 \times 10^{-9}$ M, $10^{-8}$ M, $10^{-7}$ M, $10^{-6}$ M. Data represent the mean values of n = 3 biological replicates ± s.d. Statistically significant differences were determined using two-tailed Student's t-test (**** represents p<0.0001, *** represents p<0.001, ** represents p<0.01, * represents p<0.1, ns represents not significant).

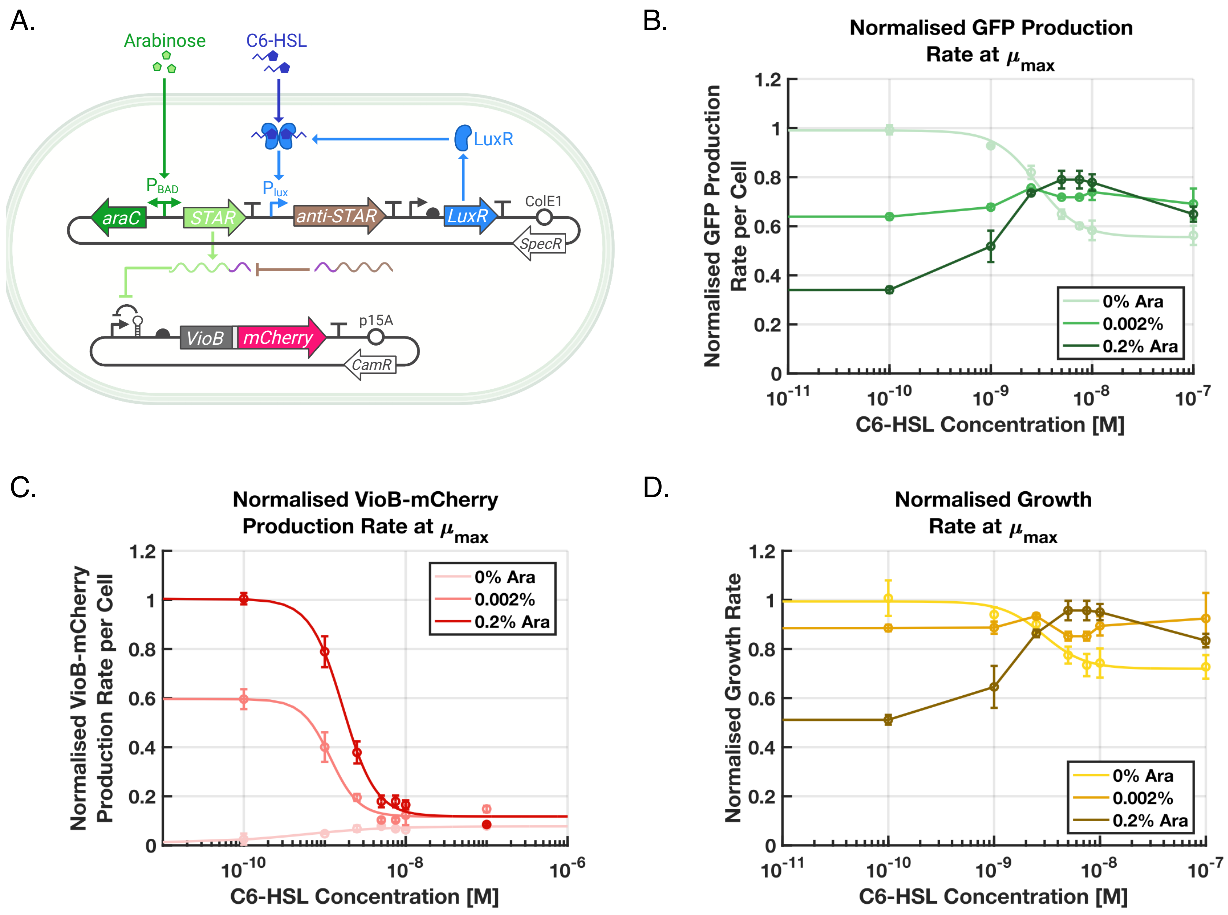

**<u>Supplementary Figure 11:</u> A burden-based control system using the STAR-based comparator controlling growth rate through VioB-mCherry expression**. (A) The LLL comparator is linked to VioB-mCherry production to control growth rate. DH10B-GFP is carrying the pAB300 and pAB517 plasmids (Supplementary Tables 1 and 3). (B) Effect of increasing C6-HSL concentration on GFP capacity (GFP production rate per cell) for three different concentrations of L-arabinose. (C) Effect of increasing C6-HSL concentration on VioB-mCherry production rate per cell for three different concentrations of L-arabinose. (D) Effect of increasing C6-HSL concentration on growth rate for three different concentrations of L-arabinose. The system was induced with a range of C6-HSL concentration from 0M to $10^{-7}$M. Growth rates and GFP capacities were normalised with the growth rate of the system induced with 0% L-arabinose and 0M C6-HSL. VioB-mCherry expressions were normalised with the VioB-mCherry production rate per cell of the system induced with 0.2% L-arabinose and 0M C6-HSL. Growth was monitored in a time-course plate-reader assay. Data represent the mean values of n = 3 biological replicates ± s.d.

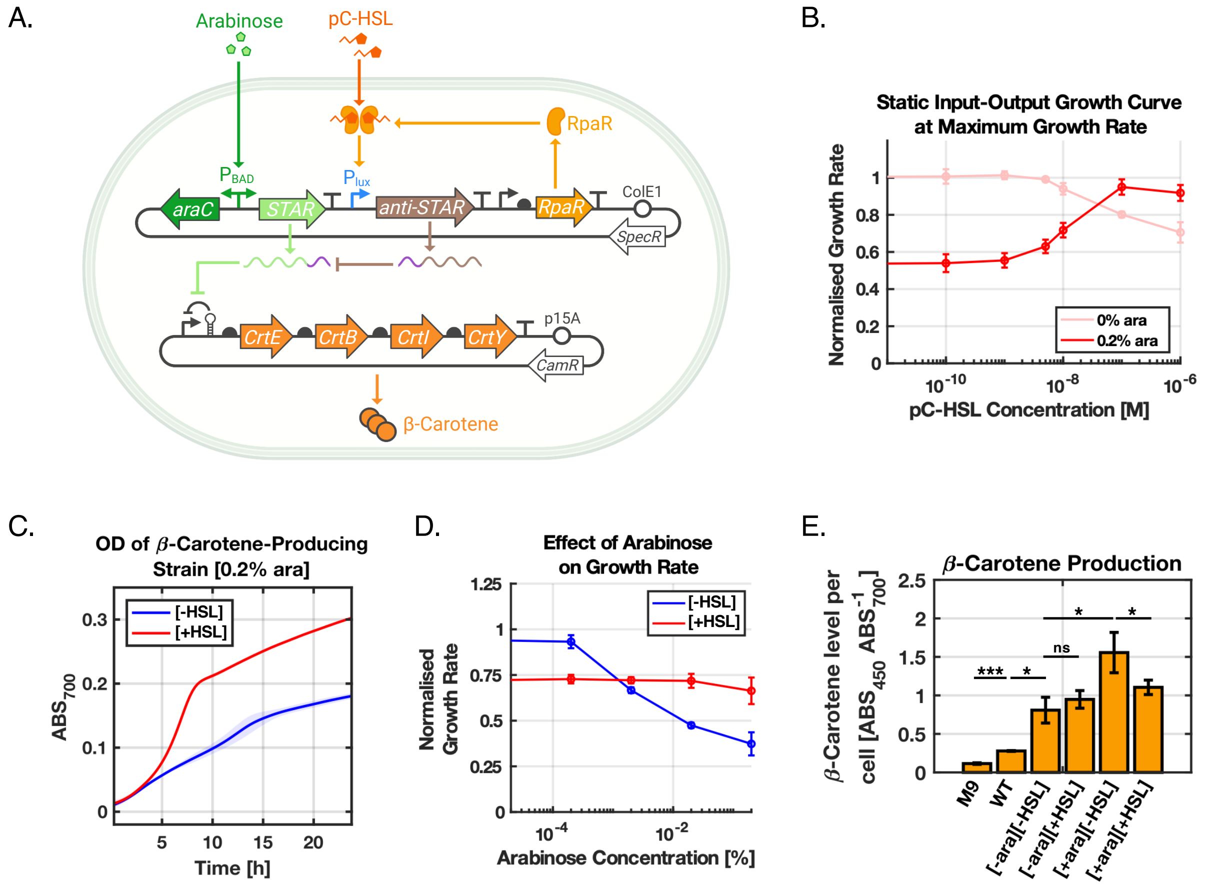

**Supplementary Figure 12: A burden-based control system using the STAR-based comparator controlling growth rate through β-carotene expression.** (A) The STAR-based comparator (with Toehold 2) based on the LRR quorum sensing system is linked to β-carotene production to control growth rate in DH10B carrying pAB401 and pAB550 (Supplementary Table 3). β-carotene is produced through the expression of four enzymes: CrtE, CrtB, CrtI and CrtY organised in an operon as described by Borkowski et al. [1].(B) Effect of increasing pC-HSL concentration on growth rate when the cells are induced with either 0% L-arabinose or 0.2% L-arabinose. Cells were induced with a range of pC-HSL from 0M to 10−6M. Growth rate was normalised with the growth rate of the system induced with 0% L-arabinose and 0M of pC-HSL. (C) Effect of adding pC-HSL on the growth profile of the cells expressing maximal levels of STAR. Cells were induced with 0.2% L- arabinose. In the "-HSL" conditions, 0M of pC-HSL was externally added to the media, while in the "+HSL" condition, 10−7M of pC-HSL was added. (D) Effect of increasing L-arabinose concentration on normalised growth rate with 0M or $10^{-7}$M of pC-HSL was added to the media ("-HSL" and "+HSL" respectively). Growth rate was normalised with the growth rate of the system induced with 0% L-arabinose and 0M of pC-HSL. (E) β-carotene production per cell measured as the absorbance read at 450 nm divided by the absorbance read at 700 nm. The conditions refer to the concentration of inducer externally added to the media. "-ara": 0% of L-arabinose, "+ara": 0.2% of L-arabinose, "-HSL": 0M of pC-HSL, "+HSL": $10^{-7}$M of pC-HSL. Experiments were carried out in DH10B. Growth was monitored in a time-course plate-reader assay. Data represent the mean values of n = 3 biological replicates ± s.d. Statistically significant differences were determined using two-tailed Student’s t-test (**** represents p<0.0001, *** represents p<0.001, ** represents p<0.01, * represents p<0.1, ns represents not significant).

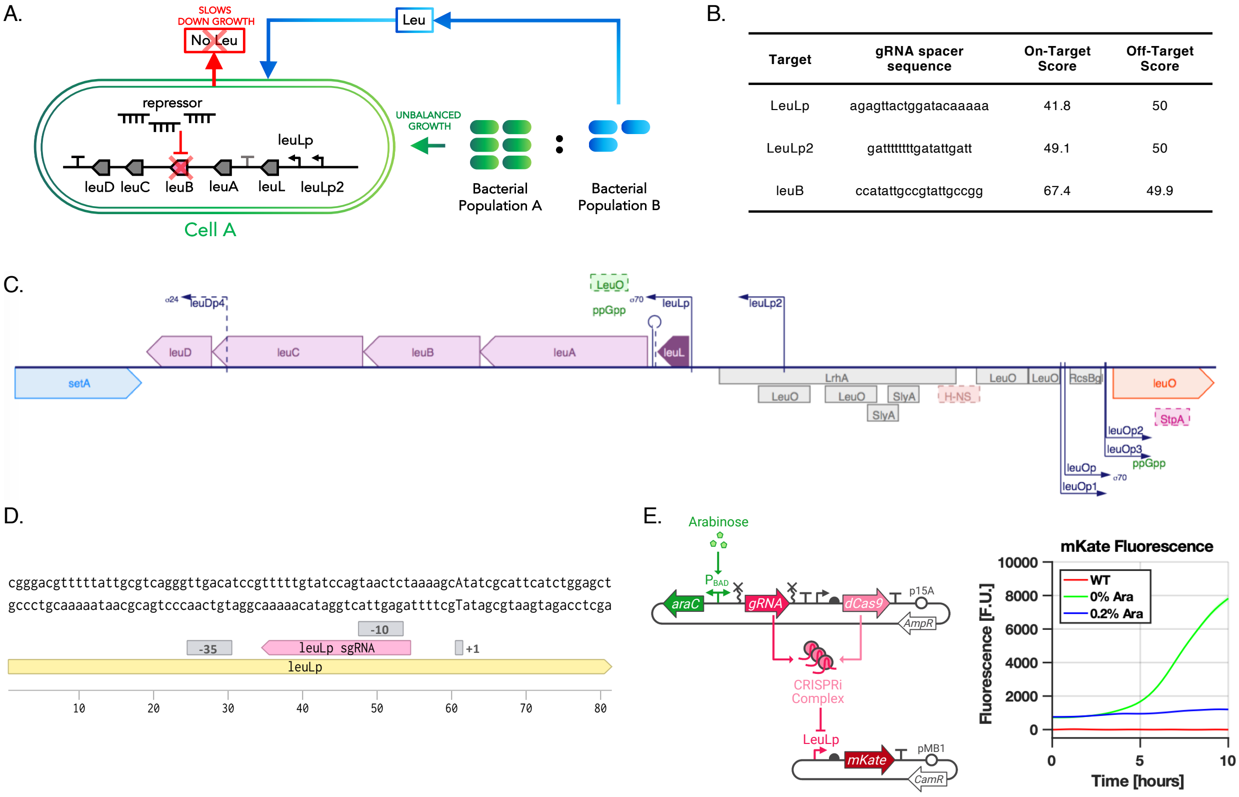

| Target | gRNA spacer sequence | On-Target Score | Off-Target Score |
|---|---|---|---|
| LeuLp | agagttactggatacaaaaa | 41.8 | 50 |
| LeuLp2 | gattttttttgatattgatt | 49.1 | 50 |
| leuB | ccatattgccgtattgccgg | 67.4 | 49.9 |

**Supplementary Figure 13: Growth rate modulation via reversible auxotrophy**. (A) When the coculture composition is out of balance, an RNA-based repressor is expressed to tune growth rate and as a result correct coculture composition. (B) Designing an sgRNA to build a leucine knockdown using CRISPRi. (C) Structure of the native leucine operon in BW25113 from EcoCyc [187]. (D) The LeuLp promoter annotated with the -35, -10 and +1 sites. The pink "LeuLp sgRNA" annotation identifies the promoter region that will be targeted by the single guide RNA (sgRNA). (E) gRNA expression is driven by the araBAD promoter. The gRNA sequence is flanked by the Hammerhead ribozyme (HH) on its 5' side and by the hepatitis delta virus ribozyme (HDV) on its 3' side. dCas9 is expressed using weak constitutive promoter and RBS sequences. gRNA and dCas9 bind to form a CRISPRi complex that targets the LeuLp promoter controlling mKate expression. In the absence of L-arabinose, the LeuLp promoter drives the expression of mKate. But in the presence of L-arabinose (0.2%), gRNA is produced and forms a complex with dCas9. The complex binds to the LeuLp promoter and inhibits its activity. Experiment was carried out in BW25113 carrying the pAB81 and B0034_mKate plasmids (Supplementary Table 3). Growth and fluorescence were monitored in a time-course plate-reader assay. Data represent the mean values of n = 3 biological replicates ± s.d.

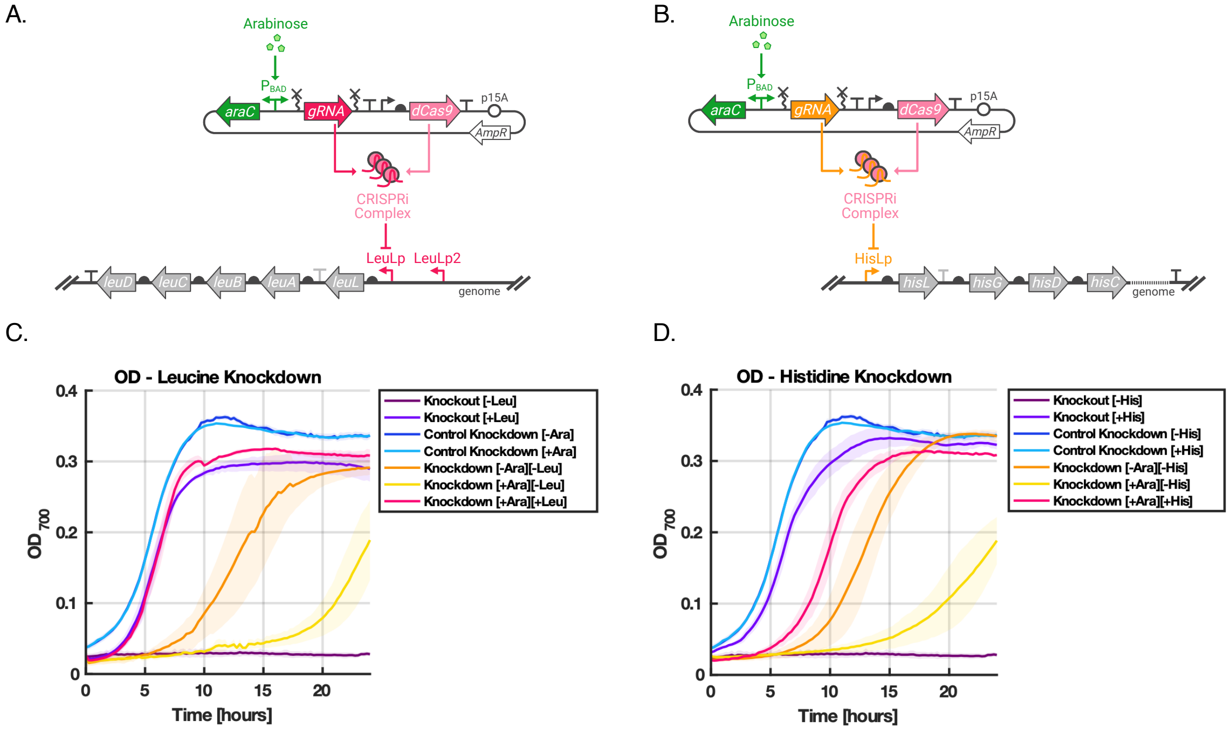

**<u>Supplementary Figure 14:</u>** **Targeting the LeuLp and HisLp promoters with CRISPRi**. (A) The circuit knocking down leucine production in BW25113 carrying pAB61 (Supplementary Table 3) consists in a constitutively expressed dCas9, and an L-arabinose inducible gRNA targeting the native LeuLp promoter in the genome. (B) The circuit knocking down histidine production in BW25113 carrying pAB60 (Supplementary Table 3) consists in a constitutively expressed dCas9, and an L-arabinose inducible gRNA targeting the native HisLp promoter in the genome. (C) OD of the leucine knockdown, the ΔLeuB knockout from the Keio collection and the control knockdown expressing the "No Target" gRNA from Ceroni et al. [2] (BW25113 carrying pAB58, Supplementary Table 3). Conditions are as follows: "[-Ara]": induction with 0% of L-arabinose, "[+Ara]": induction with 0.2% of L-arabinose, "[-Leu]": poor M9 medium is supplemented with 0 µg/mL of leucine, "[+Leu]": poor M9 medium is supplemented with 40 µg/mL of leucine. (D) OD of the histidine knockdown, the ΔHisD knockout from the Keio collection and the control knockdown expressing the "No Target" gRNA [2]. Conditions are as follows:"[-Ara]": induction with 0% of L-arabinose, "[+Ara]": induction with 0.2% of L-arabinose, "[-His]": poor M9 medium is supplemented with 0 µg/mL of L- histidine, "[+His]": poor M9 medium is supplemented with 40 µg/mL of L-histidine. OD measurements were collected in a time-course plate-reader assay. Data represent the mean values of n = 3 biological replicates ± s.d.

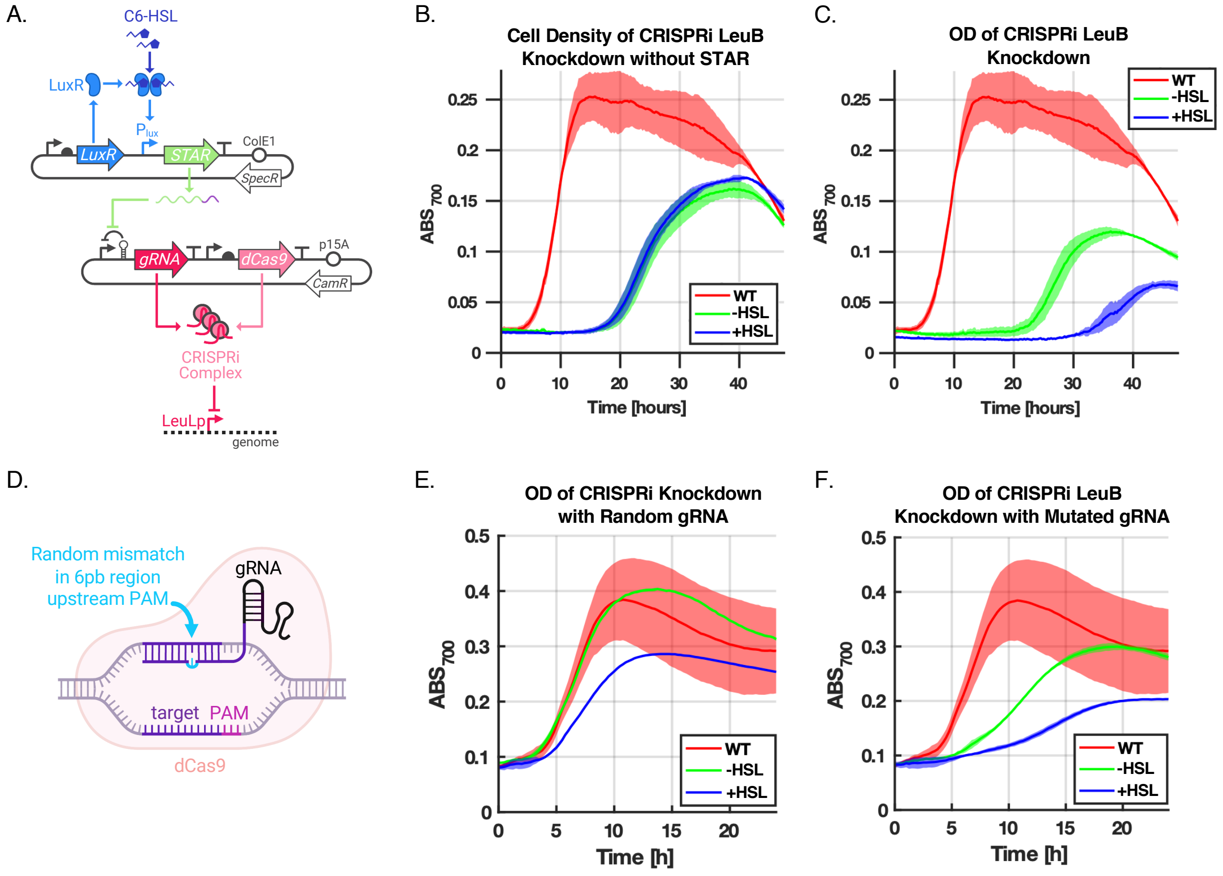

**<u>Supplementary Figure 15:</u>** **Using CRISPRi to knockdown the leucine operon**. (A) The LLL STAR system controls gRNA production targeting the LeuLp promoter in BW25113 carrying pAB127 and either pAB58, pAB96 or pAB205 (Supplementary Table 3). In the presence of STAR, gRNA is transcribed and forms an inhibiting complex with dCas9 that targets to LeuLp promoter to repress expression of the leucine operon. (B) In the absence the plasmid expressing STAR, OD of the cells is inhibited by leaky transcription of gRNA. (C) In the presence of both plasmids, $10^{-7}$M C6-HSL ("+HSL") activates STAR expression, in turn increasing gRNA production, which consequently inhibits cellular growth and extends lag-phase. (D) Building a random library of gRNA containing 1 bp mutation in the 6 bp region preceding the PAM sequence flanking the targeting region in the LeuLp promoter. (E) OD of the cells with the double plasmid system expressing a random gRNA designed with R2oDNA Designer [3]. (F) OD of the cells with the double plasmid system expressing a mutated gRNA (A→C in the bp preceding the PAM sequence). "WT" shows OD of the wild-type BW25113 strain. "-HSL" and "+HSL" show growth of the cell containing the CRISPRi plasmid when 0M and 10−7M of C6-HSL is added externally. Growth was monitored in a time-course plate-reader assay. Data represent the mean values of n = 3 biological replicates ± s.d.

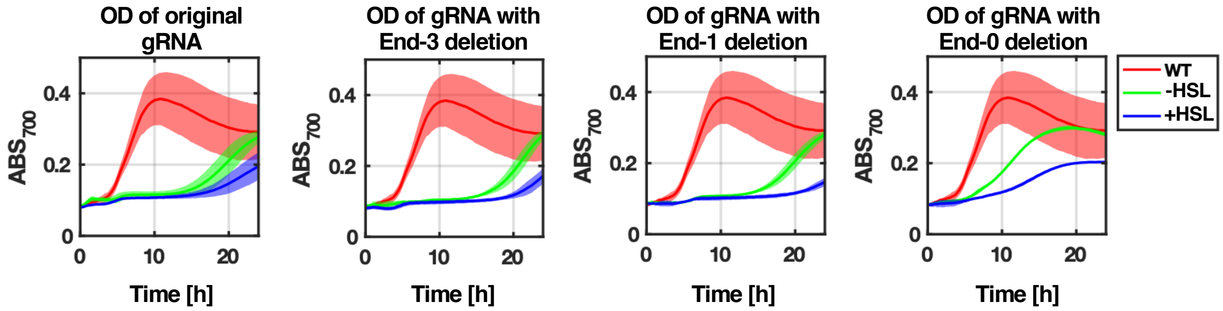

**<u>Supplementary Figure 16:</u>** **Mutations in gRNA targeting LeuLp promoter.** The mutated version of the gRNA targeting the LeuLp promoter is expressed under the control of STAR expressed in the presence of C6-HSL in BW25113 carrying pAB127 and either pAB96, pAB205, pAB206 or pAB208 (Supplementary Table 3). Here we show the effect of adding 0 M or $10^{-7}$ M of C6-HSL on cell growth for different gRNA containing either no mutation ("Original gRNA"), or mutations at 4 bp, 2 bp or directly before the PAM sequence (denoted "End-3", "End-2" and "End-0" respectively). "WT" shows OD of the wild-type BW25113 strain. "-HSL" and "+HSL" show growth of the cell containing the CRISPRi plasmid when 0M and 10−7M of C6-HSL is added externally. OD measurements were collected in a plate-reader. Data represent the mean values of n = 3 biological replicates ± s.d.

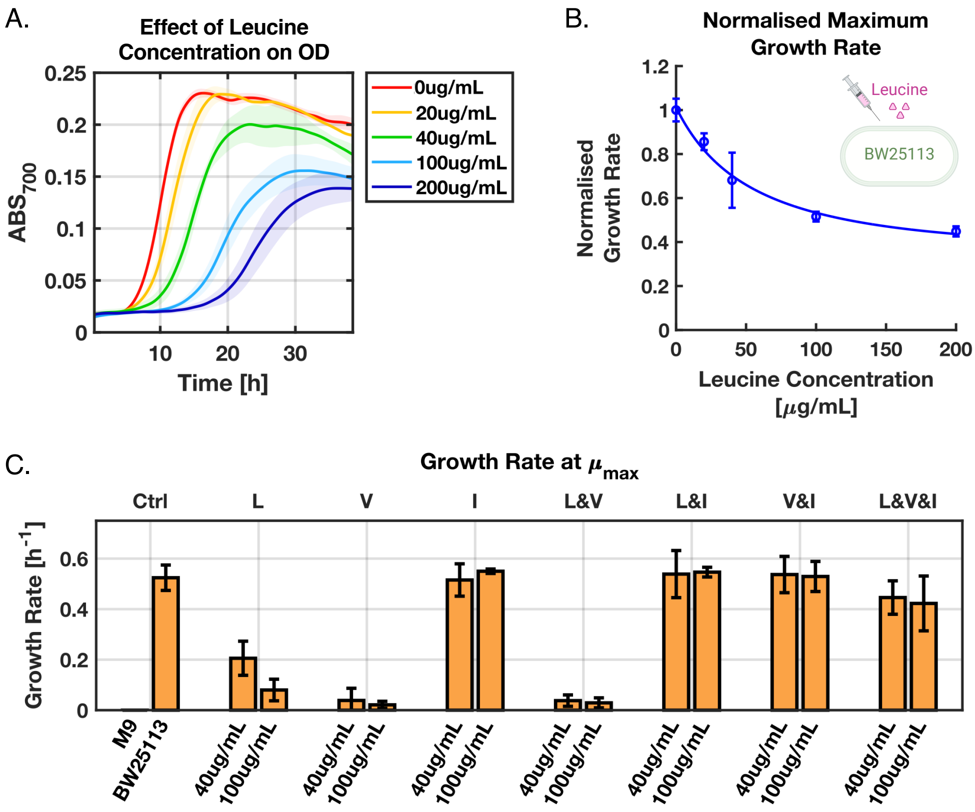

**<u>Supplementary Figure 17:</u>** **Growth rate modulation via external chemical control.** (A) BW25113 cells were grown in amino-acid-deficient M9 media with different concentrations of L-leucine. L-leucine causes toxicity by inhibiting the expression of the ilvGM and ilvBN operons unless isoleucine is present in the media. (B) Maximum growth rate of BW25113 when different concentrations of leucine are exogenously added to the media. (C) Maximum growth rate of BW25113 when either 40 µg/mL or 100 µg/mL of L-leucine, L-valine and L-isoleucine are added to the M9 minimal medium. Combinations of supplemented amino acids are represented at the top of the graph, where “L” denotes addition of L-leucine, “V” of L-valine and “I” of L-isoleucine. Data represent the mean values of n = 3 biological replicates ± s.d.

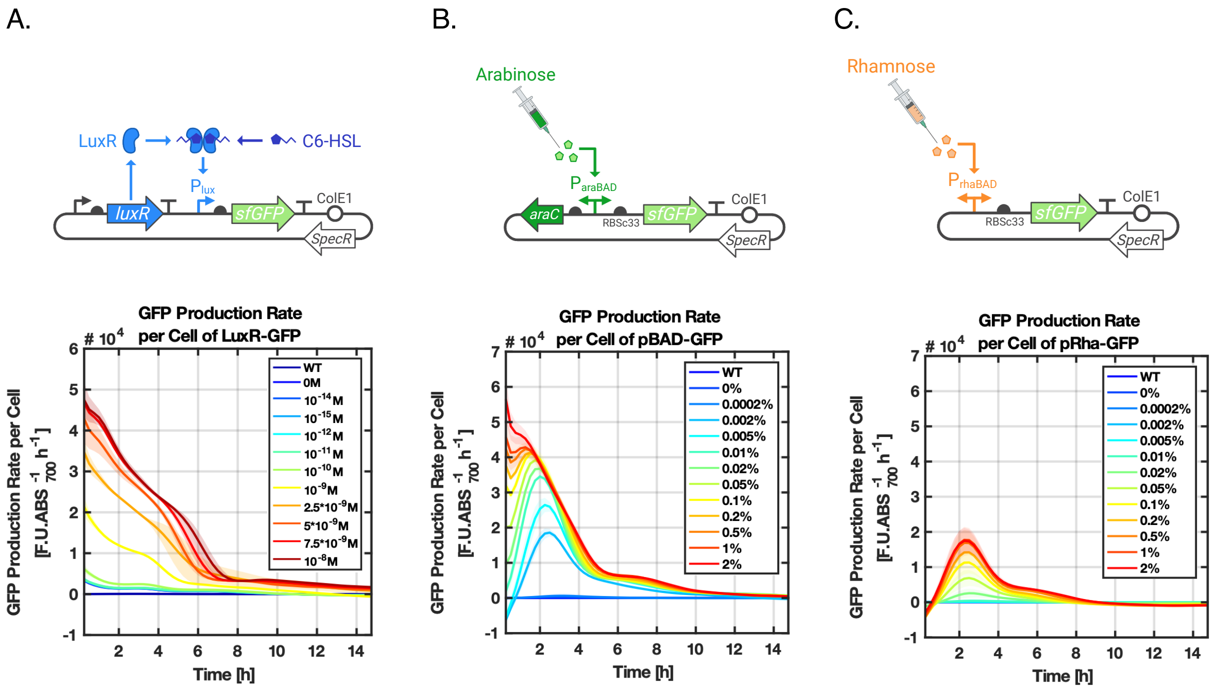

**<u>Supplementary Figure 18:</u>** **Comparison of the LLL, araBAD and rhaBAD inducible systems.** (A) sfGFP is regulated by the LLL quorum sensing system in BW25113 carrying pAB252 (Supplementary Table 3), such that sfGFP production rate per cell increases as C6-HSL concentration increases. (B) sfGFP production rate per cell as a function of increasing L-arabinose concentration. The araBAD promoter driving expression of sfGFP was externally induced with L-arabinose in BW25113 carrying pAB409. (C) sfGFP production rate per cell as a function of increasing L-arabinose concentration. The rhaBAD promoter driving expression of sfGFP was externally induced with L-rhamnose in BW25113 carrying pAB410. Fluorescence measurements were collected in a plate-reader. Data represent the mean values of n = 3 biological replicates ± s.d.

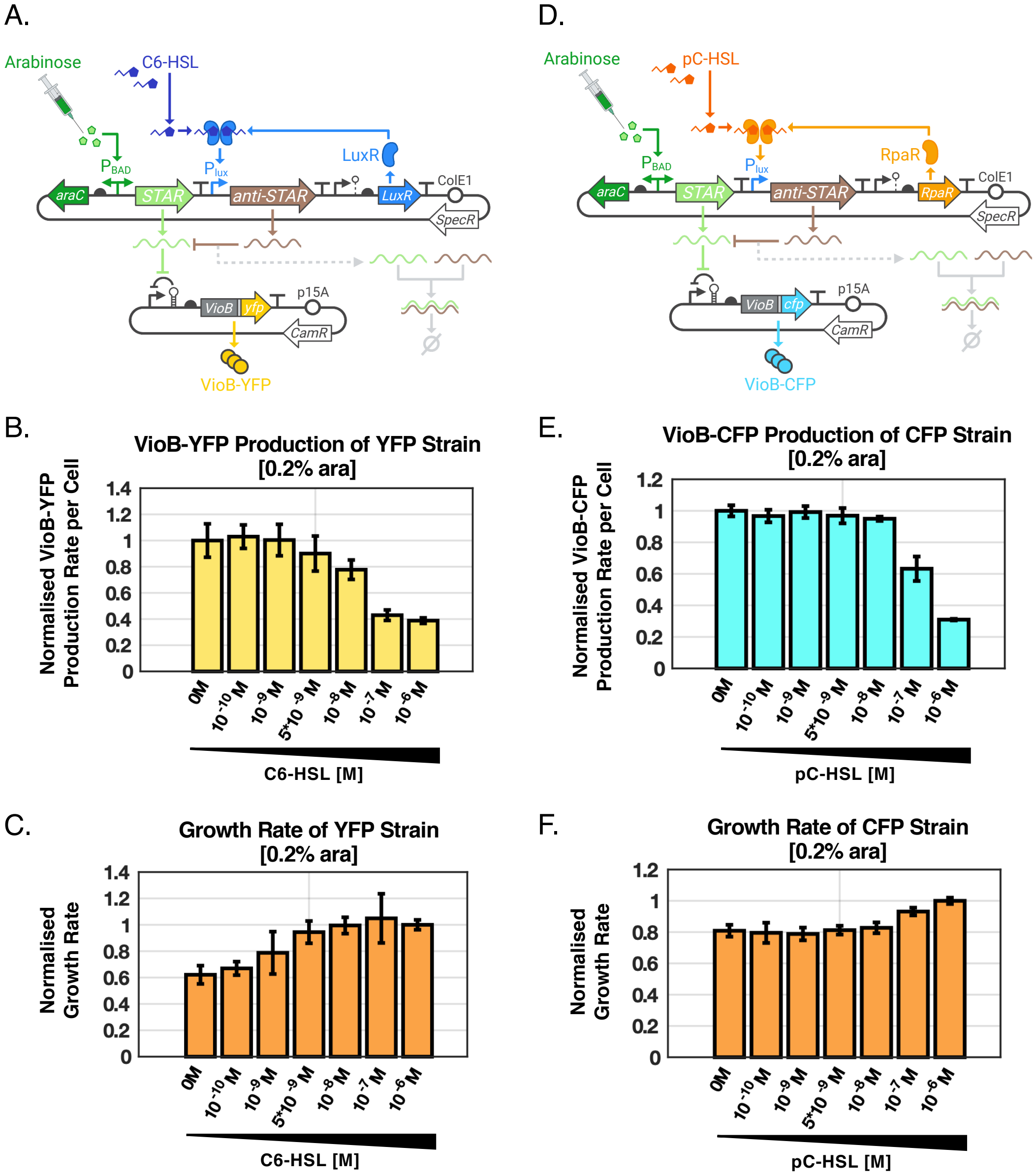

**Supplementary Figure 19: Building coculture strains with different burden levels.** (A) The YFP strain regulates VioB- sfYFP production under the control of the LLL STAR-based comparator with Toehold 2 in DH10B carrying pAB518 and pAB300 (Supplementary Table 3). VioB- sfYFP is activated by L-arabinose and inhibited with increasing concentration of C6-HSL. (B-C) The YFP strain is induced with 0.2% of L-arabinose to activate VioB-sfYFP expression. Increasing C6-HSL levels gradually turns down VioB-sfYFP and improves growth rate. (D) The CFP strain regulates VioB-sfCFP expression using the LRR STAR-based comparator with Toehold 2 in DH10B carrying pAB519 and pAB401. (E-F) The CFP strain was induced with 0.2% L-arabinose to activate VioB-sfCFP expression. Increasing pC-HSL concentration gradually turns down VioB-sfYFP expression and recovers growth rate. Fluorescence and OD measurements were collected in a plate-reader. Data represent the mean values of n = 3 biological replicates ± s.d.

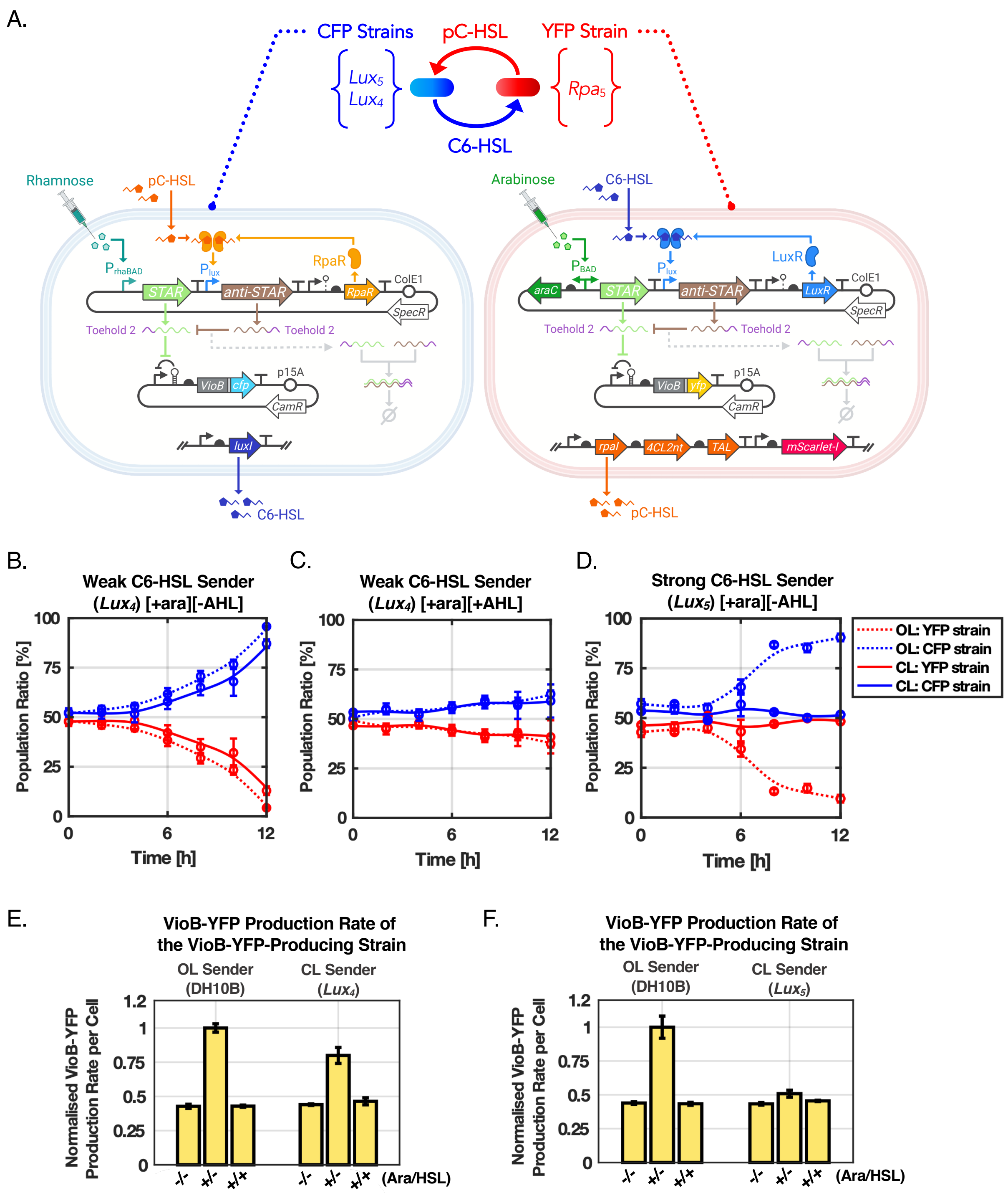
A.
CFP Strains
pC-HSL
YFP Strain
Lux5
Lux4
Rpa5
C6-HSL
Rhamnose
pC-HSL
RpaR
Arabinose
C6-HSL
LuxR
Toehold 2
p15A
ColE1
C6-HSL
pC-HSL
B.
Weak C6-HSL Sender (Lux4) [+ara][-AHL]
C.
Weak C6-HSL Sender (Lux4) [+ara][+AHL]
D.
Strong C6-HSL Sender (Lux5) [+ara][-AHL]
Population Ratio [%]
Time [h]
OL: YFP strain
OL: CFP strain
CL: YFP strain
CL: CFP strain
E.
VioB-YFP Production Rate of the VioB-YFP-Producing Strain
OL Sender (DH10B)
CL Sender (Lux4)
Normalised VioB-YFP Production Rate per Cell
(Ara/HSL)
F.
VioB-YFP Production Rate of the VioB-YFP-Producing Strain
OL Sender (DH10B)
CL Sender (Lux5)

**Supplementary Figure 20:** **Effect of C6-HSL production on coculture composition.** (A) *$Lux_4$* and *$Lux_5$* C6-HSL sender strains, or CFP strains, are tested to study the effect of C6-HSL signal strength on coculture composition of the closed- loop circuit. The YFP strain produces VioB-YFP in the presence of L-arabinose and VioB-YFP production is down-regulated by C6-HSL in *$Rpa_5$* carrying pAB518 and pAB300 (Supplementary Tables 1 and 3). The CFP strain produces VioB-CFP in the presence of L-rhamnose and VioB-CFP production is down-regulated by pC-HSL in *$Lux_4$* or *$Lux_5$* carrying pAB519 and pAB537 (Supplementary Tables 1 and 3). In the open-loop circuit, to break communication between the strains, neither strain produces quorum sensing molecule. (B) Change in population ratio when the coculture is made up of the weak C6-HSL *$Lux_4$* sender CFP strain and the *$Rpa_5$* YFP strain. Coculture is induced with 0.2% of L-arabinose, 0% of L-rhamnose, and 0M of both C6-HSL and pC-HSL. (C) Change in population ratio when the coculture is made up of the weak C6-HSL *$Lux_4$* sender CFP strain and the *$Rpa_5$* YFP strain, when the coculture is induced with 0.2% of L-arabinose, 0% of L-arabinose, 10−7M of both C6-HSL and pC-HSL. (D) Change in population ratio when the coculture is made up of the strong C6-HSL *$Lux_5$* sender CFP strain and the *$Rpa_5$* YFP strain. The coculture is induced with 0.2% of L-arabinose, 0% L- rhamnose and 0M of both C6-HSL and pC-HSL. (E-F) Normalised VioB-sfYFP production rate per cell for the open-loop and closed-loop cocultures with the *$Lux_4$* and *$Lux_5$* C6-HSL sender strains. Three inducer conditions were tested by externally adding different combinations of the four inducers. (1) "-/-": 0% of L-arabinose, 0% of L-rhamnose, 0M of pC-HSL, 0M of C6-HSL. (2) "+/-": 0.2% of L-arabinose, 0% of L-rhamnose, 0M of pC-HSL, 0M of C6-HSL. (3) "+/+": 0.2% of L-arabinose, 0% of L-rhamnose, 10−7M of pC-HSL, 0M of C6-HSL. The population ratio was measured through flow-cytometry. VioB-sfYFP fluorescence was measured in a timecourse plate-reader assay. Data represent the mean values of n = 3 biological replicates ± s.d.

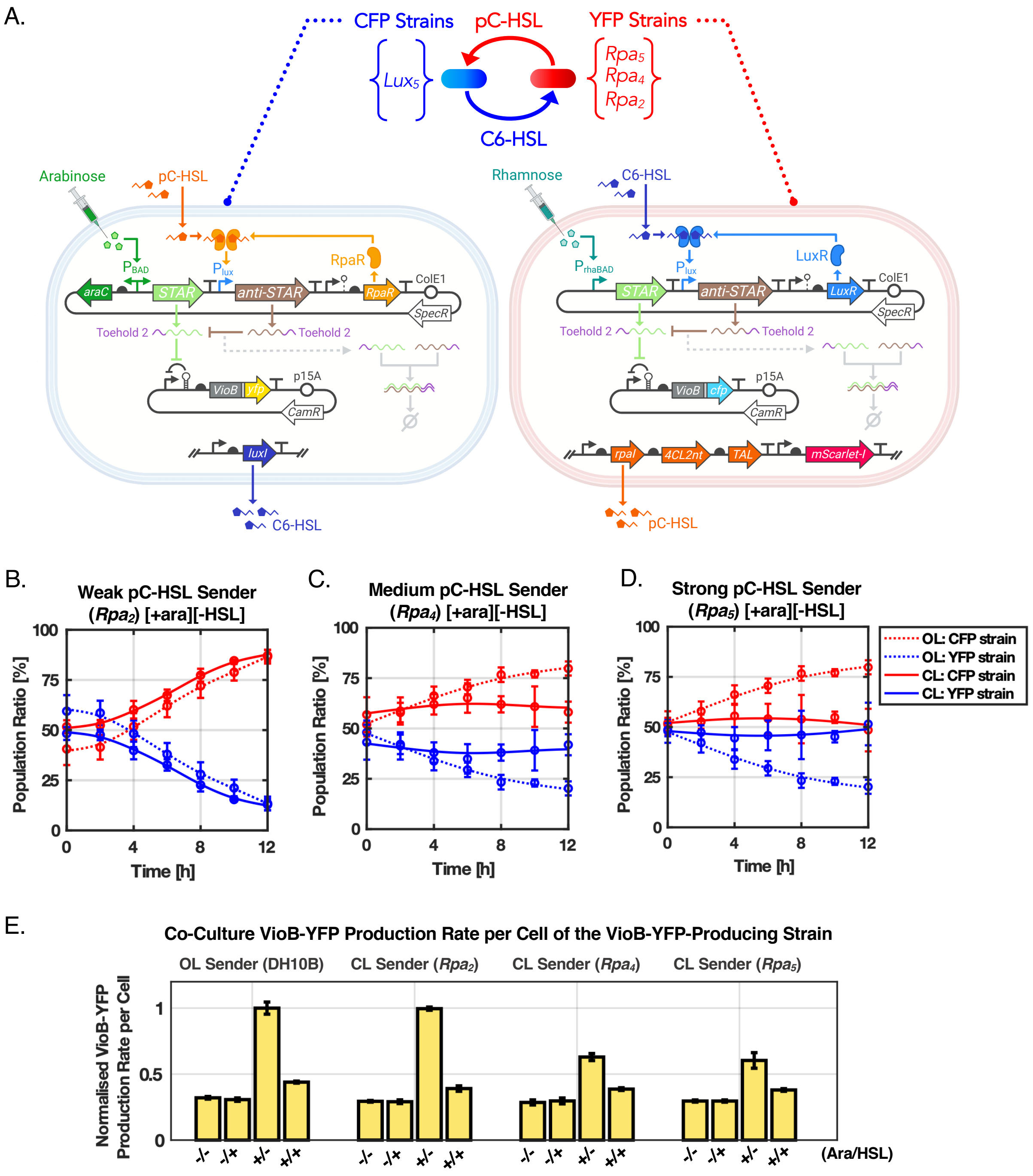
A.
CFP Strains
pC-HSL
YFP Strains
Lux5
Rpa5
Rpa4
Rpa2
C6-HSL
Arabinose
pC-HSL
RpaR
araC
STAR
anti-STAR
RpaR
ColE1
SpecR
Toehold 2
Toehold 2
VioB
yfp
p15A
CamR
luxI
C6-HSL
Rhamnose
C6-HSL
LuxR
STAR
anti-STAR
LuxR
ColE1
SpecR
Toehold 2
Toehold 2
VioB
cfp
p15A
CamR
rpaI
4CL2nt
TAL
mScarlet-I
pC-HSL
B.
Weak pC-HSL Sender
(Rpa2) [+ara][-HSL]
C.
Medium pC-HSL Sender
(Rpa4) [+ara][-HSL]
D.
Strong pC-HSL Sender
(Rpa5) [+ara][-HSL]
Population Ratio [%]
Time [h]
OL: CFP strain
OL: YFP strain
CL: CFP strain
CL: YFP strain
E.
Co-Culture VioB-YFP Production Rate per Cell of the VioB-YFP-Producing Strain
OL Sender (DH10B)
CL Sender (Rpa2)
CL Sender (Rpa4)
CL Sender (Rpa5)
Normalised VioB-YFP Production Rate per Cell
-/-
-/+
+/-
+/+
(Ara/HSL)

**<u>Supplementary Figure 21:</u>** **Effect of pC-HSL production on coculture composition.** (A) $Rpa_{2,4,5}$ pC-HSL sender strains, or CFP strains, are tested to study the effect of pC-HSL signal strength on coculture composition of the closed- loop circuit. The YFP strain produces VioB-YFP when L-arabinose is present and VioB-YFP production is down-regulated by pC-HSL in $Lux_5$ carrying pAB518 and pAB401 (Supplementary Tables 1 and 3). The CFP strain produces VioB-CFP in the presence of L-rhamnose and VioB-CFP production is down-regulated by C6-HSL in $Rpa_{2,4,5}$ carrying pAB519 and pAB399 (Supplementary Tables 1 and 3). In the open-loop circuit, to break communication between the strains, neither of the strains produce quorum sensing molecules. (B) Change in population ratio when the coculture is made up of the weak pC-HSL $Rpa_2$ sender CFP strain and the $Lux_5$ YFP strain. Coculture is induced with 0.2% of L-arabinose and 0% of L-rhamnose. (C) Change in population ratio when the coculture is made up of the medium-strength pC-HSL $Rpa_4$ sender CFP strain and the $Lux_5$ YFP strain. Coculture is induced with 0.2% of L-arabinose and 0% of L-rhamnose. (D) Change in population ratio when the coculture is made up of the strong pC-HSL $Rpa_5$ sender CFP strain and the $Lux_5$ YFP strain. Coculture is induced with 0.2% of L-arabinose and 0% of L-rhamnose. E) Normalised VioB-sfYFP production rate per cell of the open-loop coculture and closed-loop coculture with the $Rpa_{2,4,5}$ pC-HSL sender strains. Four inducer conditions were tested by externally adding different combinations of the four inducers. (1) "-/-": 0% of L-arabinose, 0% of L-rhamnose, 0M of pC-HSL, 0M of C6-HSL. (2) "-/+": 0% of L-arabinose, 0% of L-rhamnose, 10−7M of pC-HSL, 0M of C6-HSL. (3) "+/-": 0.2% of L-arabinose, 0% of L-rhamnose, 0M of pC-HSL, 0M of C6-HSL. (4) "+/+": 0.2% of L-arabinose, 0% of L-rhamnose, 10−7M of pC-HSL, 0M of C6-HSL. The population ratio was measured through flow-cytometry. VioB-sfYFP fluorescence was measured in a timecourse plate-reader assay. Data represent the mean values of n = 3 biological replicates ± s.d.

**Supplementary Table 1: Strains**

| Name | Genotype | Antibiotic | Source/Parent |
|---|---|---|---|
| DH10B | K-12 strain, F– mcrA Δ(mrr-hsdRMS-mcrBC) ϕ80lacZΔM15 ΔlacX74 recA1 endA1 araD139 Δ(ara,leu)7697 galU galK λ–rpsL(StrR) nupG | Streptomycin | Invitrogen |
| BW25113 | K-12 strain, acI+rrnBT14 ΔlacZWJ16 hsdR514 ΔaraBADAH33 ΔrhaBADLD78 rph-1 Δ(araB–D)567 Δ(rhaD–B)568 ΔlacZ4787(::rrnB-3) hsdR514 rph-1 | None | Keio Collection |
| BW25113(rpsL150 ) | rpsL150 (strR) | Streptomycin | BW25113 |
| PIR2 One Shot | F- Δlac169 rpoS(Am) robA1 creC510 hsdR514 endA recA1 uidA(ΔMluI)::pir | None | Invitrogen |
| TransforMax EC100D pir+ | F- mcrA Δ(mrr-hsdRMS-mcrBC) ϕ80dlacZΔM15 ΔlacX74 recA1 endA1 araD139 Δ(ara, leu)7697 galU galK λ- rpsL nupG pir+(DHRF) | None | Lucigen |
| JW5807 | ΔleuB | None | BW25113 |
| DH10B-GFP | J23100–sfGFP–BBa_1002 at attλ | Kanamycin | DH10B |
| DH10B-mScarlet | J23100–mScarlet-I–BBa_1002 at attλ | Kanamycin | DH10B |
| BW25113-GFP | J23100–sfGFP–BBa_1002 at attλ | Kanamycin | BW25113 |
| BW25113-mScarlet | J23100–mScarlet-I–BBa_1002 at attλ | Kanamycin | BW25113 |
| Lux1 | J23100–RBSc33–luxI–L3S2P55 at attλ (CRIM plasmid pAB386) | Kanamycin | DH10B |
| Lux2 | J23115–B0030–luxI–L3S2P55 at attλ (CRIM plasmid pAB566) | Kanamycin | DH10B |
| Lux3 | J23105–B0030–luxI–L3S2P55 at attλ (CRIM plasmid pAB564) | Kanamycin | DH10B |
| Lux4 | J23106–B0030–luxI–L3S2P55 at attλ (CRIM plasmid pAB554) | Kanamycin | DH10B |
| Lux5 | J23100–B0030–luxI–L3S2P55 at attλ plasmid (CRIM plasmid pAB553) | Kanamycin | DH10B |
| Rpa1 | J23109–B0030–rpaI–B0030–4CL2nt–B0032– TAL–L3S2P55 at attλ (CRIM plasmid pAB572) | Kanamycin | DH10B |
| Rpa2 | J23109–B0030–rpaI–B0030–4CL2nt–B0030– TAL–L3S2P55 at attλ (CRIM plasmid pAB496) | Kanamycin | DH10B |
| Rpa3 | J23106–B0030–rpaI–B0030–4CL2nt–B0030– TAL–L3S2P55 at attλ (CRIM plasmid pAB495) | Kanamycin | DH10B |
| Rpa4 | J23105–B0030–rpaI–B0030–4CL2nt–B0030– TAL–L3S2P55 at attλ (CRIM plasmid pAB494) | Kanamycin | DH10B |
| Rpa5 | J23114–B0030–rpaI–B0030–4CL2nt–B0030– TAL–L3S2P55 at attλ (CRIM plasmid pAB493) | Kanamycin | DH10B |
| Rpa6 | J23100–B0030–rpaI–B0030–4CL2nt–B0032– TAL–L3S2P55 at attλ (CRIM plasmid pAB482) | Kanamycin | DH10B |

**Supplementary Table 2:** **Genetic parts**

| Part | Type | DNA Sequence (5' to 3') | Source |
|---|---|---|---|
| J23119 | Promoter | TTTACGGCTAGCTCAGTCCTAGGTATTGTGCTAGC | iGEM J23119 |
| J23100 | Promoter | TTTACGGCTAGCTCAGTCCTAGGTATTGTGCTAGC | iGEM J23100 |
| J23106 | Promoter | TTTACGGCTAGCTCAGTCCTAGGTATTGTGCTAGC | iGEM J23106 |
| J23105 | Promoter | TTTACGGCTAGCTCAGTCCTAGGTATTGTGCTAGC | iGEM J23105 |
| J23115 | Promoter | TTTACGGCTAGCTCAGTCCTAGGTATTGTGCTAGC | iGEM J23115 |
| J23114 | Promoter | TTTACGGCTAGCTCAGTCCTAGGTATTGTGCTAGC | iGEM J23114 |
| J23109 | Promoter | TTTACGGCTAGCTCAGTCCTAGGTATTGTGCTAGC | iGEM J23109 |
| $P_{RAND14}$ | Promoter | TTTACGGCTAGCTCAGTCCTAGGTATTGTGCTAGC | 4 |
| $P_{lux}$ | Promoter | ACCTGTAGGATCGTACAGGTTTACGCAAGAAAATGGTTTGTTATAGTCGAATAAA | 5 |
| $P_{luxA}$ | Promoter | ACCTGTAGGATCGTACAGGTTTACGCAAGAAAATGGTTTGTTATTGTCGAATAAA | 5 |
| $P_{luxB}$ | Promoter | ACCTGTAGGATCGTACAGGTTTACGCAAGAAAATGGTTTGTTACAGTCGAATAAA | 5 |
| $P_{rpa}$ | Promoter | GCACCTGTCCGATCGGACAGTATTACGCAAGAAAATGGTTTGTTATAGTCGAATAT | 6 |
| $P_{rhaBAD}$ | Promoter | CCACAATTCAGCAAATTGTGAACATCATCACGTTCATCTTTCCCTGGTTGCCAATGGCCCATTTTCCTGTCAGT<br>AACGAGAAGGTCGCGTATTCAGGCGCTTTTTAGACTGGTCGTAATGAA | 4 |
| AraC-$P_{araBAD}$ | Promoter | TTATGACAACTTGACGGCTACATCATTCACTTTTTCTTCACAACCGGCACGGAACTCGCTCGGGCTGGCCCCG<br>GTGCATTTTTTAAATACCCGCGAGAAATAGAGTTGATCGTCAAAACCAACATTGCGACCGACGGTGGCGATA<br>GGCATCCGGGTGGTGCTCAAAAGCAGCTTCGCCTGGCTGATACGTTGGTCCTCGCGCCAGCTTAAGACGCTA<br>ATCCCTAACTGCTGGCGGAAAAGATGTGACAGACGCGACGGCGACAAGCAAACATGCTGTGCGACGCTGGC<br>GATATCAAAATTGCTGTCTGCCAGGTGATCGCTGATGTACTGACAAGCCTCGCGTACCCGATTATCCATCGGT<br>GGATGGAGCGACTCGTTAATCGCTTCCATGCGCCGCAGTAACAATTGCTCAAGCAGATTTATCGCCAGCAGC<br>TCCGAATAGCGCCCTTCCCCTTGCCCGGCGTTAATGATTTGCCCAAACAGGTCGCTGAAATGCGGCTGGTGC<br>GCTTCATCCGGGCGAAAGAACCCCGTATTGGCAAATATTGACGGCCAGTTAAGCCATTCATGCCAGTAGGCG<br>CGCGGACGAAAGTAAACCCACTGGTGATACCATTCGCGAGCCTCCGGATGACGACCGTAGTGATGAATCTCT<br>CCTGGCGGGAACAGCAAAATATCACCCGGTCGGCAAACAAATTCTCGTCCCTGATTTTTCACCACCCCCTGAC<br>CGCGAATGGTGAGATTGAGAATATAACCTTTCATTCCCAGCGGTCGGTCGATAAAAAAATCGAGATAACCGT<br>TGGCCTCAATCGGCGTTAAACCCGCCACCAGATGGGCATTAAACGAGTATCCCGGCAGCAGGGGATCATTTT<br>GCGCTTCAGCCATACTTTTCATACTCCCGCCATTCAGAGAAGAAACCAATTGTCCATATTGCATCAGACATTG<br>CCGTCACTGCGTCTTTTACTGGCTCTTCTCGCTAACCAAACCGGTAACCCCGCTTATTAAAAGCATTCTGTAAC<br>AAAGCGGGACCAAAGCCATGACAAAAACGCGTAACAAAAGTGTCTATAATCACGGCAGAAAAGTCCACATT<br>GATTATTTGCACGGCGTCACACTTTGCTATGCCATAGCATTTTTATCCATAAGATTAGCGGATCCTACCTGAC<br>GCTTTTTATCGCAACTCTCTACTGTTTCTCCATACCCGTTTTTTTGGGCTAGC | 4 |
| B0030 | RBS | TCTAGAGTCACACAGGAAACCTACTAG | iGEM B0030 |
| B0031 | RBS | TCTAGAGTCACACAGGAAACCTACTAG | iGEM B0031 |
| B0032 | RBS | TCTAGAGTCACACAGGAAAGTACTAG | iGEM B0032 |
| B0033 | RBS | TCTAGAGTCACACAGGACTACTAG | iGEM B0033 |
| B0034 | RBS | TACTAGAGAAAGAGGAGAAATACTAG | iGEM B0034 |
| RBS8B | RBS | AAGAATTCAAAAGCTCTACAGAGGAGAAAGGATAT | 4 |
| RBSc33 | RBS | CTACGTTTTTTAGAAAAAGGAGGTATGCGAG | 7 |
| RBSc44 | RBS | ATCGGATTGGATCCAAAGGAGGTTATACCG | 7 |
| $RBS_{Bujard}$ | RBS | GAATTCATTAAAGAGGAGAAAGGTACC | 8 |
| $RBS_{VioB}$ | RBS | AGGCATCTTTCAACTAAAGTAAGAGGTAATAATT | 4 |
| $RBS_{mCherry}$ | RBS | AGGCTGCAGCGAAAGCG | 4 |
| $RBS_{RFP}$ | RBS | ATGGCGAGTAGCG | 8 |
| $RBS_{ctrE}$ | RBS | TAAGGAGGCTCCTA | 1 |
| $RBS_{ctrB}$ | RBS | AGAGGATACATATA | 1 |
| $RBS_{ctrI}$ | RBS | TCAGGATTTTTGTA | 1 |
| $RBS_{ctrY}$ | RBS | CGAGGAGGTAATAA | 1 |
| HH | Ribozyme | ATCTGGCTGATGAGTCCGTGAGGACGAAACGAGTAAGCTCGTCCC | 9 |
| HDV | Ribozyme | GGCCGGCATGGTCCCAGCCTCCTCGCTGGCGCCGGCTGGGCAACACCTTCGGGTGGCGAATGG GACT | 9 |
| RiboJ | Ribozyme | AGCTGTCACCGGATGTGCTTTCCGGTCTGATGAGTCCGTGAGGACGAAACAGCCTCTACAAAT<br>AATTTTGTTTAA | 5 |
| L3S3P21 | Terminator | CCAATTATTGAAGGCCTCCCTAACGGGGGGCCTTTTTTTGTTTCTGGTCTCCC | 10 |
| ECK120010793 | Terminator | TACGTAAAAACCCGCTTCGGCGGGTTTTTACTTT | 10 |

| | | | |
|---|---|---|---|
| L3S2P55 | Terminator | CTCGGTACCAAAGACGAACAATAAGACGCTGAAAAGCGTCTTTTTT | 10 |
| L3S2P21 | Terminator | CTCGGTACCAAATTCCAGAAAAGAGGCCTCCCGAAAGGGGGGCCTTTTTTCGTTTTGGTCC | 10 |
| ECK120033737 | Terminator | GGAAACACAGAAAAAAGCCCGCACCTGACAGTGCGGGCTTTTTTTTTCGACCAAAGG | 10 |
| ECK120029600 | Terminator | TTCAGCCAAAAAACTTAAGACCGCCGGTCTTGTCCACTACCTTGCAGTAATGCGGTGGACAGGATCGGCGGT<br>TTTCTTTTCTCTTCTCAA | 10 |
| t500 | Terminator | CAAAGCCCGCCGAAAGGCGGGCTTTTTTTT | 11 |
| B1001 | Terminator | AAAAAAAAACCCCGCTTCGGCGGGGTTTTTTTTT | iGEM B1001 |
| B1002 | Terminator | CGCAAAAAACCCCGCTTCGGCGGGGTTTTTTCGC | iGEM B1002 |
| B0015 | Terminator | CCAGGCATCAAATAAAACGAAAGGCTCAGTCGAAAGACTGGGCCTTTCGTTTTATCTGTTGTTTGTCGGTGA<br>ACGCTCTCTACTAGAGTCACA CTGGCTCACCTTCGGGTGGGCCTTTCTGCGTTTATA | iGEM B0015 |
| tL3 | Terminator | AATGGCGATGACGCATCCTCACGATAATATCCGGGTAGGCGCAATCACTTTCGTCTCTACTCCGTTACAAAGC<br>GAGGCTGGGTATTTCCCGGCCTTTCTGTTATCCGAAATCCACTGAAAGCACAGCGGCTGGCTGAGGAGATAA<br>ATAATAAACGAGGGGCTGTATGCACAAAGCATCTTCTGTTGAGTTAAGAACGAGTATCGAGATGGCACATA<br>GCCTTGCTCAAATTGGAATCAGGTTTGTGCCAATACCAGTAGAAACAGACGAAGAA | 12 |
| rgnB | Terminator | GATGGTAGTGTGGGGTCTCCCCATGCGAGAGTAGGGAACTGCCAGGCATCAAATAAAACGAAAGGCTCAGT<br>CGAAAGACTGGGCCTTTCGTTTTATCTGTTGTTTGTCGGTGAACGCTCTCCTGAGTAGGACAAATCCGCCGG<br>GAGCGGATTTGAACGTTGCGAAGCAACGGCCCGGAGGGTGGCGGGCAGGACGCCCGCCATAAACTGCCAG<br>GCATCAAATTAAGCAGAAGGCCATCCTGACGGATGGCCTTTTTGCGT | 12 |
| rrnB T1 | Terminator | ATTTGTCCTACTCAGGAGAGCGTTCACCGACAAACAACAGATAAAACGAAAGGCCCAGTCTTTCGACTGAGC<br>CTTTCGTTTTATTTG | 7 |
| Lambda t0 | Terminator | GACTCCTGTTGATAGATCCAGTAATGACCTCAGAACTCCATCTGGATTTGTTCAGAACGCTCGGTTGCCGCCG<br>GGCGTTTTTTATTGGTGAGAAT | 7 |
| His operon terminator | Terminator | TCCGGCAAAAAAGGGCAAGGTGTCACCACCCTGCCCTTTTTCTTTAAAACCGAAAAGA | 13 |
| LuxR | Gene | ATGAAAAACATAAATGCCGACGACACATACAGAATAATTAATAAAATTAAAGCTTGTAGAAGCAATAATGAT<br>ATTAATCAATGCTTATCTGATATGACTAAAATGGTACATTGTGAATATTATTTACTCGCGATCATTTATCCTCA<br>TTCTATGGTTAAATCTGATATTTCAATCCTAGATAATTACCCTAAAAAATGGAGGCAATATTATGATGACGCT<br>AATTTAATAAAATATGATCCTATAGTAGATTATTCTAACTCCAATCATTCACCAATTAATTGGAATATATTTGA<br>AAACAATGCTGTAAATAAAAAATCTCCAAATGTAATTAAAGAAGCGAAAACATCAGGTCTTATCACTGGGTT<br>TAGTTTCCCTATTCATACGGCTAACAATGGCTTCGGAATGCTTAGTTTTGCACATTCAGAAAAAGACAACTAT<br>ATAGATAGTTTATTTTTACATGCGTGTATGAACATACCATTAATTGTTCCTTCTCTAGTTGATAATTATCGAAA<br>AATAAATATAGCAAATAATAAATCAAACAACGATTTAACCAAAAGAGAAAAAGAATGTTTAGCGTGGGCAT<br>GCGAAGGAAAAAGCTCTTGGGATATTTCAAAAATATTAGGTTGCAGTGAGCGTACTGTCACTTTCCATTTAA<br>CCAATGCGCAAATGAAACTCAATACAACAAACCGCTGCCAAAGTATTTCTAAAGCAATTTTAACAGGAGCAA<br>TTGATTGCCCATACTTTAAAAATTAA | 8 |
| RpaR | Gene | ATGATCGTCGGCGAAGATCAGCTTTGGGGACGGCGTGCGCTGGAATTCGTCGATTCCGTCGAACGGCTCGA<br>GGCGCCGGCGCTGATCAGCCGGTTCGAATCGCTGATCGCGAGCTGCGGATTTACCGCCTACATCATGGCCG<br>GCCTGCCGTCGCGCAATGCCGGACTACCGGAGCTGACGCTGGCCAATGGCTGGCCGCGAGACTGGTTCGAT<br>CTGTATGTCAGCGAAAACTTCAGCGCGGTCGATCCGGTGCCGCGCCACGGCGCTACCACGGTTCATCCTTTC<br>GTATGGTCCGATGCACCCTACGACCGCGACCGTGATCCGGCCGCCCACCGGGTCATGACCCGGGCGGCGGA<br>ATTCGGACTGGTCGAGGGTTACTGCATTCCGCTGCACTACGACGACGGTAGCGCCGCGATCAGCATGGCCG<br>GCAAGGATCCGGACCTCAGCCCGGCCGCGCGCGGCGCGATGCAGCTGGTCAGCATCTACGCGCATAGTCGC<br>CTGCGCGCACTCAGCCGGCCAAAGCCGATCCGGCGCAACCGGCTCACGCCGCGCGAGTGCGAGATCCTGCA<br>ATGGGCAGCGCAGGGCAAGACCGCCTGGGAAATCTCGGTAATCCTCTGCATCACCGAACGCACGGTGAAAT<br>TCCATCTGATCGAAGCCGCCCGCAAGCTCGACGCCGCCAACCGCACCGCGGCGGTTGCCAAGGCATTGACG<br>CTCGGATTGATCCGTTTGTAA | 6 |
| LuxI | Gene | ATGACTATAATGATAAAAAAATCGGATTTTTTGGCAATTCCATCGGAGGAGTATAAAGGTATTCTAAGTCTTC<br>GTTATCAAGTGTTTAAGCAAAGACTTGAGTGGGACTTAGTTGTAGAAAATAACCTTGAATCAGATGAGTATG<br>ATAACTCAAATGCAGAATATATTTATGCTTGTGATGATACTGAAAATGTAAGTGGATGCTGGCGTTTATTACC<br>TACAACAGGTGATTATATGCTGAAAAGTGTTTTTCCTGAATTGCTTGGTCAACAGAGTGCTCCCAAAGATCCT<br>AATATAGTCGAATTAAGTCGTTTTGCTGTAGGTAAAAATAGCTCAAAGATAAATAACTCTGCTAGTGAAATTA<br>CAATGAAACTATTTGAAGCTATATATAAACACGCTGTTAGTCAAGGTATTACAGAATATGTAACAGTAACATC<br>AACAGCAATAGAGCGATTTTTAAAGCGTATTAAAGTTCCTTGTCATCGTATTGGAGACAAAGAAATTCATGT<br>ATTAGGTGATACTAAATCGGTTGTATTGTCTATGCCTATTAATGAACAGTTTAAAAAAGCAGTCTTAA | iGEM C0061 |
| RpaI | Gene | ATGCAGGTTCATGTCATCCGTCGAGAGAACCGCGCGCTCTATGCCGGTCTGCTCGAAAAGTACTTCCGCATC<br>CGTCACCAGATCTACGTCGTCGAGCGCGGCTGGAAGGAGCTCGATCGGCCGGATGGCCGCGAGATCGATCA<br>GTTCGACACCGAAGACGCCGTGTATCTGCTCGGCGTCGACAATGACGACATCGTCGCCGGCATGCGGATGG<br>TGCCGACCACGTCACCGACGCTCCTCAGCGACGTCTTCCCGCAGCTTGCGCTGGCAGGCCCGGTGCGGCGG<br>CCGGATGCCTACGAGCTGTCGCGGATCTTCGTGGTACCGCGCAAGCGCGGCGAGCATGGCGGCCCGCGCGC<br>CGAAGCCGTGATCCAGGCGGCCGCGATGGAGTACGGCCTGTCGATCGGTCTGTCGGCCTTCACCATCGTGC<br>TGGAGACCTGGTGGCTGCCGCGACTGGTGGACCAGGGCTGGAAGGCAAAGCCGCTCGGCCTGCCTCAGGA<br>CATCAACGGATTCTCGACCACCGCAGTGATCGTCGACGTCGACGACGACGCCTGGGTCGGCATCTGCAATCG<br>CCGCTCGGTGCCCGGACCCACGCTGGAATGGCGCGGGCTCGAAGCCATCCGCCGTCATTCGCTTCCGGAATT<br>CCAGGTGATTTCATAA | CIDAR MoClo Extension[i] |
| TAL | Gene | ATGACCCAGGTGGTTGAACGCCAGGCCGATCGCCTGAGTAGTCGTGAATACTTAGCTCGCGTCGTTCGTAGT<br>GCCGGCTGGGATGCGGGCCTAACCTCTTGTACAGATGAAGAAATTGTTCGCATGGGCGCGTCAGCCCGCAC<br>CATCGAGGAATATTTAAAAAGTGATAAACCGATTTATGGTTTAACCCAAGGCTTCGGCCCGCTGGTACTGTTT<br>GATGCGGATAGCGAATTAGAACAGGGTGGTAGCCTGATTAGCCATCTGGGCACCGGTCAGGGCGCGCCGC<br>TGGCGCCGGAAGTGAGTCGTTTAATTCTGTGGCTGCGTATTCAAAACATGCGCAAAGGTTATAGCGCCGTTA<br>GCCCGGTTTTCTGGCAAAAACTGGCAGACCTATGGAATAAAGGCTTTACCCCGGCAATTCCGCGTCATGGTA<br>CCGTTTCCGCCTCGGGTGATCTGCAACCGCTGGCGCATGCCGCGCTGGCATTCACCGGCGTGGGTGAAGCG<br>TGGACCCGCGATGCAGATGGCCGCTGGAGCACCGTTCCGGCTGTTGATGCCCTGGCAGCGCTGGGTGCCGA<br>ACCGTTTGATTGGCCTGTCCGCGAAGCACTGGCGTTTGTTAATGGCACCGGAGCCAGCCTGGCGGTTGCTGT<br>TTTAAATCATCGTTCTGCCCTGCGCCTGGTTCGCGCGTGTGCGGTACTGAGCGCACGCCTGGCGACCCTGCT<br>GGGCGCAAATCCGGAACATTATGACGTTGGTCATGGCGTTGCCCGCGGTCAGGTTGGCCAGCTGACCGCGG<br>CGGAATGGATTCGTCAGGGCCTGCCACGTGGTATGGTGCGCGATGGAAGCCGTCCGTTGCAGGAACCTTAT<br>AGCCTTCGCTGCGCTCCGCAGGTTCTAGGCGCTGTTCTGGATCAGCTGGACGGTGCGGGTGACGTGCTGGC | 14 |

[i] CIDAR MoClo Extension, Volume I was a gift from Richard Murray (Addgene kit #1000000161).

| | | | |
|---|---|---|---|
| | | CCGCGAAGTTGATGGTTGCCAGGATAACCCTATTACCTACGAAGGTGAATTGCTGCATGGCGGTAACTTCCA<br>TGCCATGCCGGTTGGTTTTGCAAGTGATCAGATTGGTCTGGCGATGCACATGGCGGCCTACCTGGCTGAACG<br>CCAGCTGGGCCTGCTGGTTAGCCCGGTAACCAATGGTGATTTACCACCGATGCTGACCCCGCGTGCCGGCCG<br>TGGTGCGGGTCTTGCTGGCGTCCAGATTTCTGCCACCAGCTTCGTTTCTCGTATTCGCCAACTGGTTTTCCCG<br>GCGTCTCTGACCACCCTGCCGACCAACGGTTGGAATCAAGACCATGTACCGATGGCACTGAATGGCGCTAAT<br>AGCGTTTTCGAAGCACTGGAACTGGGTTGGTTAACCGTTGGAAGCCTGGCGGTGGGCGTTGCACAGCTGGC<br>GGCGATGACCGGTCATGCGGCTGAAGGGGTTTGGGCAGAACTGGCAGGCATTTGCCCGCCGTTAGATGCC<br>GACCGTCCGCTGGGTGCGGAAGTTCGCGCAGCCCGTGATCTGCTGAGCGCGCACGCTGATCAGCTGTTGGT<br>GGACGAAGCCGATGGTAAAGACTTTGGCTAA | |
| 4CL2nt | Gene | ATGGAGAAAGACACGAAGCAAGTTGACATCATTTTTCGCTCGAAACTGCCGGACATTTACATTCCGAATCAT<br>CTGCCGCTGCATAGCTACTGCTTCGAGAACATTTCTGAATTTTCTAGCCGTCCGTGTCTGATTAACGGTGCCA<br>ATAAACAGATCTATACGTACGCGGACGTCGAGTTGAACAGCCGTAAGGTCGCAGCGGGTCTGCACAAGCAA<br>GGCATCCAGCCTAAAGATACCATCATGATTCTGTTGCCAAATTCTCCGGAGTTTGTGTTTGCGTTTATCGGCG<br>CAAGCTACCTGGGTGCGATTAGCACGATGGCAAATCCGCTGTTTACCCCGGCTGAGGTTGTTAAACAAGCAA<br>AAGCCAGCAGCGCGAAGATCATCGTGACCCAAGCATGCCACGTCAACAAAGTTAAGGACTATGCCTTCGAA<br>AATGACGTCAAGATCATTTGCATCGATAGCGCGCCTGAAGGTTGTCTGCATTTCAGCGTTCTGACGCAGGCT<br>AACGAACACGATATTCCGGAAGTTGAGATTCAGCCGGACGATGTGGTGGCCCTGCCGTACTCCAGCGGTAC<br>CACCGGCCTGCCGAAAGGCGTTATGCTGACCCACAAGGGCCTGGTGACGAGCGTCGCCCAGCAGGTCGATG<br>GTGAAAACCCGAACCTGTACATCCACAGCGAAGATGTTATGCTGTGTGTTCTGCCACTGTTCCACATCTATTC<br>CCTGAACAGCGTCCTGCTGTGCGGCCTGCGTGTGGGCGCTGCCATTTTGATTATGCAGAAGTTTGACATTGT<br>CAGCTTCTTGGAACTGATCCAACGCTACAAGGTGACGATCGGTCCGTTCGTCCCGCCGATTGTTTTGGCCATT<br>GCAAAAAGCCCAATGGTGGATGACTATGACCTGTCGAGCGTGCGTACCGTGATGTCCGGTGCAGCGCCGCT<br>GGGCAAAGAGCTGGAGGATACCGTTCGTGCGAAGTTTCCGAATGCGAAACTGGGTCAAGGCTACGGTATG<br>ACTGAAGCAGGTCCGGTGCTGGCGATGTGCTTGGCGTTCGCGAAAGAGCCGTTCGAAATCAAAAGCGGTGC<br>GTGCGGTACCGTGGTGCGTAATGCTGAAATGAAAATTGTGGATCCGAAAACCGGCAACAGCCTGCCGCGCA<br>ACCAGAGCGGTGAGATTTGTATTCGCGGTGACCAGATTATGAAGGGCTACCTGAATGACCCGGAGGCCACT<br>GCGCGTACGATCGACAAAGAGGGTTGGCTGTATACCGGCGACATCGGTTATATCGATGACGACGACGAGCT<br>GTTCATCGTTGATCGCCTGAAAGAGTTGATTAAGTACAAGGGTTTCCAAGTTGCGCCTGCGGAACTGGAGGC<br>TCTGCTGTTGAATCATCCGAACATTAGCGATGCAGCAGTCGTTCCGATGAAGGATGAGCAGGCGGGTGAAG<br>TTCCGGTCGCGTTTGTTGTGCGTAGCAACGGCAGCACGATCACCGAGGATGAGGTAAAGGATTTCATTTCCA<br>AACAAGTCATCTTCTATAAGCGTATCAAGCGTGTGTTTTTCGTCGATGCAATCCCGAAAAGCCCGTCCGGTAA<br>GATCCTGCGCAAAGACTTGCGTGGAAGCTGGCGGCAGGTCTGCCGAATTAGTAA | 14 |
| dCas9 | Gene | ATGGATAAGAAATACTCAATAGGCTTAGCTATCGGCACAAATAGCGTCGGATGGGCGGTGATCACTGATGA<br>ATATAAGGTTCCGTCTAAAAAGTTCAAGGTTCTGGGAAATACAGACCGCCACAGTATCAAAAAAAATCTTAT<br>AGGGGCTCTTTTATTTGACAGTGGAGAGACAGCGGAAGCGACTCGTCTCAAACGGACAGCTCGTAGAAGGT<br>ATACACGTCGGAAGAATCGTATTTGTTATCTACAGGAGATTTTTTCAAATGAGATGGCGAAAGTAGATGATA<br>GTTTCTTTCATCGACTTGAAGAGTCTTTTTTGGTGGAAGAAGACAAGAAGCATGAACGTCATCCTATTTTTGG<br>AAATATAGTAGATGAAGTTGCTTATCATGAGAAATATCCAACTATCTATCATCTGCGAAAAAAATTGGTAGAT<br>TCTACTGATAAAGCGGATTTGCGCTTAATCTATTTGGCCTTAGCGCATATGATTAAGTTTCGTGGTCATTTTTT<br>GATTGAGGGAGATTTAAATCCTGATAATAGTGATGTGGACAAACTATTTATCCAGTTGGTACAAACCTACAA<br>TCAATTATTTGAAGAAAACCCTATTAACGCAAGTGGAGTAGATGCTAAAGCGATTCTTTCTGCACGATTGAGT<br>AAATCAAGACGATTAGAAAATCTCATTGCTCAGCTCCCCGGTGAGAAGAAAAATGGCTTATTTGGGAATCTC<br>ATTGCTTTGTCATTGGGTTTGACCCCTAATTTTAAATCAAATTTTGATTTGGCAGAAGATGCTAAATTACAGCT<br>TTCAAAAGATACTTACGATGATGATTTAGATAATTTATTGGCGCAAATTGGAGATCAATATGCTGATTTGTTT<br>TTGGCAGCTAAGAATTTATCAGATGCTATTTTACTTTCAGATATCCTAAGAGTAAATACTGAAATAACTAAGG<br>CTCCCCTATCAGCTTCAATGATTAAACGCTACGATGAACATCATCAAGACTTGACTCTTTTAAAAGCTTTAGTT<br>CGACAACAACTTCCAGAAAAGTATAAAGAAATCTTTTTTGATCAATCAAAAAACGGATATGCAGGTTATATT<br>GATGGGGGAGCTAGCCAAGAAGAATTTTATAAATTTATCAAACCAATTTTAGAAAAAATGGATGGTACTGA<br>GGAATTATTGGTGAAACTAAATCGTGAAGATTTGCTGCGCAAGCAACGGACCTTTGACAACGGCTCTATTCC<br>CCATCAAATTCACTTGGGTGAGCTGCATGCTATTTTGAGAAGACAAGAAGACTTTTATCCATTTTTAAAAGAC<br>AATCGTGAGAAGATTGAAAAAATCTTGACTTTTCGAATTCCTTATTATGTTGGTCCATTGGCGCGTGGCAATA<br>GTCGTTTTGCATGGATGACTCGGAAGTCTGAAGAAACAATTACCCCATGGAATTTTGAAGAAGTTGTCGATA<br>AAGGTGCTTCAGCTCAATCATTTATTGAACGCATGACAAACTTTGATAAAAATCTTCCAAATGAAAAAGTACT<br>ACCAAAACATAGTTTGCTTTATGAGTATTTTACGGTTTATAACGAATTGACAAAGGTCAAATATGTTACTGAA<br>GGAATGCGAAAACCAGCATTTCTTTCAGGTGAACAGAAGAAAGCCATTGTTGATTTACTCTTCAAAACAAAT<br>CGAAAAGTAACCGTTAAGCAATTAAAAGAAGATTATTTCAAAAAAATAGAATGTTTTGATAGTGTTGAAATT<br>TCAGGAGTTGAAGATAGATTTAATGCTTCATTAGGTACCTACCATGATTTGCTAAAAATTATTAAAGATAAAG<br>ATTTTTTGGATAATGAAGAAAATGAAGATATCTTAGAGGATATTGTTTTAACATTGACCTTATTTGAAGATAG<br>GGAGATGATTGAGGAAAGACTTAAAACATATGCTCACCTCTTTGATGATAAGGTGATGAAACAGCTTAAACG<br>TCGCCGTTATACTGGTTGGGGACGTTTGTCTCGAAAATTGATTAATGGTATTAGGGATAAGCAATCTGGCAA<br>AACAATATTAGATTTTTTGAAATCAGATGGTTTTGCCAATCGCAATTTTATGCAGCTGATCCATGATGATAGT<br>TTGACATTTAAAGAAGACATTCAAAAAGCACAAGTGTCTGGACAAGGCGATAGTTTACATGAACATATTGCA<br>AATTTAGCTGGTAGCCCTGCTATTAAAAAAGGTATTTTACAGACTGTAAAAGTTGTTGATGAATTGGTCAAA<br>GTAATGGGGCGGCATAAGCCAGAAAATATCGTTATTGAAATGGCACGTGAAAATCAGACAACTCAAAAGGG<br>CCAGAAAAATTCGCGAGAGCGTATGAAACGAATCGAAGAAGGTATCAAAGAATTAGGAAGTCAGATTCTTA<br>AAGAGCATCCTGTTGAAAATACTCAATTGCAAAATGAAAAGCTCTATCTCTATTATCTCCAAAATGGAAGAG<br>ACATGTATGTGGACCAAGAATTAGATATTAATCGTTTAAGTGATTATGATGTCGATGCCATTGTTCCACAAAG<br>TTTCCTTAAAGACGATTCAATAGACAATAAGGTCTTAACGCGTTCTGATAAAAATCGTGGTAAATCGGATAAC<br>GTTCCAAGTGAAGAAGTAGTCAAAAAGATGAAAAACTATTGGAGACAACTTCTAAACGCCAAGTTAATCACT<br>CAACGTAAGTTTGATAATTTAACGAAAGCTGAACGTGGAGGTTTGAGTGAACTTGATAAAGCTGGTTTTATC<br>AAACGCCAATTGGTTGAAACTCGCCAAATCACTAAGCATGTGGCACAAATTTTGGATAGTCGCATGAATACT<br>AAATACGATGAAAATGATAAACTTATTCGAGAGGTTAAAGTGATTACCTTAAAATCTAAATTAGTTTCTGACT<br>TCCGAAAAGATTTCCAATTCTATAAAGTACGTGAGATTAACAATTACCATCATGCCCATGATGCGTATCTAAA<br>TGCCGTCGTTGGAACTGCTTTGATTAAGAAATATCCAAAACTTGAATCGGAGTTTGTCTATGGTGATTATAAA<br>GTTTATGATGTTCGTAAAATGATTGCTAAGTCTGAGCAAGAAATAGGCAAAGCAACCGCAAAATATTTCTTTT<br>ACTCTAATATCATGAACTTCTTCAAAACAGAAATTACACTTGCAAATGGAGAGATTCGCAAACGCCCTCTAAT<br>CGAAACTAATGGGGAAACTGGAGAAATTGTCTGGGATAAAGGGCGAGATTTTGCCACAGTGCGCAAAGTAT<br>TGTCCATGCCCCAAGTCAATATTGTCAAGAAAACAGAAGTACAGACAGGCGGATTCTCCAAGGAGTCAATTT<br>TACCAAAAAGAAATTCGGACAAGCTTATTGCTCGTAAAAAAGACTGGGATCCAAAAAAATATGGTGGTTTTG<br>ATAGTCCAACGGTAGCTTATTCAGTCCTAGTGGTTGCTAAGGTGGAAAAAGGGAAATCGAAGAAGTTAAAA<br>TCCGTTAAAGAGTTACTAGGGATCACAATTATGGAAAGAAGTTCCTTTGAAAAAAATCCGATTGACTTTTTAG<br>AAGCTAAAGGATATAAGGAAGTTAAAAAAGACTTAATCATTAAACTACCTAAATATAGTCTTTTTGAGTTAG<br>AAAACGGTCGTAAACGGATGCTGGCTAGTGCCGGAGAATTACAAAAAGGAAATGAGCTGGCTCTGCCAAGC<br>AAATATGTGAATTTTTTATATTTAGCTAGTCATTATGAAAAGTTGAAGGGTAGTCCAGAAGATAACGAACAA<br>AAACAATTGTTTGTGGAGCAGCATAAGCATTATTTAGATGAGATTATTGAGCAAATCAGTGAATTTTCTAAGC<br>GTGTTATTTTAGCAGATGCCAATTTAGATAAAGTTCTTAGTGCATATAACAAACATAGAGACAAACCAATACG<br>TGAACAAGCAGAAAATATTATTCATTTATTTACGTTGACGAATCTTGGAGCTCCCGCTGCTTTTAAATATTTTG<br>ATACAACAATTGATCGTAAACGATATACGTCTACAAAAGAAGTTTTAGATGCCACTCTTATCCATCAATCCAT<br>CACTGGTCTTTATGAAACACGCATTGATTTGAGTCAGCTAGGAGGTGACTAA | 4 |
| VioB | Gene | ATGAGCATTCTGGATTTCCCGCGTATCCACTTCCGTGGCTGGGCCCGTGTCAATGCGCCGACCGCGAACCGC<br>GATCCGCACGGCCACATCGATATGGCCAGCAATACCGTGGCGATGGCGGGTGAGCCGTTCGACCTGGCACG<br>CCATCCTACGGAGTTCCACCGTCACCTGCGCTCCCTGGGTCCGCGCTTCGGCTTGGATGGTCGTGCTGACCC | 13 |

| | | | |
|---|---|---|---|
| | | GGAAGGCCCGTTCAGCCTGGCCGAGGGCTACAACGCTGCCGGTAACAACCACTTTTCGTGGGAGAGCGCAA<br>CCGTTAGCCACGTGCAATGGGATGGCGGTGAGGCGGATCGTGGTGACGGTCTGGTCGGTGCTCGTTTGGCA<br>CTGTGGGGTCACTACAATGATTATCTGCGTACCACCTTCAATCGTGCTCGTTGGGTCGACAGCGACCCGACG<br>CGCCGTGACGCTGCACAAATCTATGCGGGCCAATTCACCATTAGCCCGGCTGGTGCCGGTCCGGGTACGCC<br>GTGGCTGTTTACGGCAGACATTGATGATAGCCATGGTGCACGTTGGACGCGTGGCGGCCACATTGCAGAGC<br>GTGGCGGCCACTTCTTGGATGAAGAGTTTGGTCTGGCACGCCTGTTTCAGTTCTCTGTGCCGAAAGATCACC<br>CACATTTTCTGTTTCACCCGGGTCCGTTTGATTCCGAGGCCTGGCGTCGTCTGCAATTGGCTCTGGAGGATGA<br>CGACGTTCTGGGTCTGACCGTGCAATATGCGTTGTTCAATATGAGCACCCCGCCTCAGCCGAACAGCCCGGT<br>TTTTCACGATATGGTCGGTGTTGTCGGTCTGTGGCGTCGTGGTGAACTGGCGAGCTACCCGGCTGGTCGTCT<br>GCTGCGTCCGCGTCAACCGGGTCTGGGTGACCTGACCCTGCGCGTCAACGGTGGTCGCGTTGCGCTGAATTT<br>GGCGTGTGCCATTCCGTTCAGCACTCGTGCCGCGCAGCCAAGCGCACCGGACCGCCTGACCCCGGACCTGG<br>GTGCCAAACTGCCGCTGGGCGATCTGCTGCTGCGTGATGAGGACGGCGCACTGTTGGCACGTGTGCCGCAG<br>GCTCTGTACCAAGACTATTGGACGAATCACGGTATTGTGGACCTGCCGCTGCTGCGCGAACCGCGTGGTAGC<br>TTGACCCTGAGCAGCGAACTGGCGGAGTGGCGTGAGCAAGACTGGGTCACCCAAAGCGACGCGTCTAACCT<br>GTACCTGGAGGCACCGGATCGCCGTCACGGTCGCTTTTTCCCTGAGAGCATCGCGCTGCGCAGCTACTTTCG<br>CGGTGAAGCGCGTGCGCGTCCGGATATCCCGCATCGTATCGAGGGCATGGGCCTGGTCGGCGTCGAATCTC<br>GTCAGGATGGCGACGCTGCGGAATGGCGTCTGACGGGTCTGCGTCCGGGTCCGGCACGCATTGTTCTGGAC<br>GATGGTGCCGAGGCGATCCCTCTGCGTGTTCTGCCTGACGATTGGGCGCTGGATGACGCGACCGTCGAAGA<br>AGTGGATTACGCCTTTTTGTACCGCCACGTTATGGCGTATTACGAGCTGGTGTATCCATTCATGAGCGACAA<br>GGTGTTTTCCCTGGCTGATCGTTGCAAATGTGAAACGTACGCACGTCTGATGTGGCAGATGTGTGATCCGCA<br>GAACCGCAACAAGTCCTATTACATGCCGAGCACCCGCGAACTGTCGGCACCGAAAGCTCGTTTGTTCTTGAA<br>GTATCTGGCCCACGTGGAAGGCCAGGCACGCCTGCAAGCACCTCCGCCAGCGGGTCCGGCACGCATTGAAT<br>CTAAAGCCCAGTTGGCGGCAGAGCTGCGTAAAGCCGTCGACCTGGAGCTGTCTGTGATGCTGCAATACCTG<br>TACGCGGCGTATAGCATTCCGAACTATGCACAGGGCCAACAACGTGTTCGTGACGGTGCGTGGACCGCCGA<br>GCAGCTGCAACTGGCGTGCGGTAGCGGTGACCGTCGCCGTGATGGCGGTATTCGTGCAGCACTGCTGGAAA<br>TTGCTCATGAAGAAATGATTCATTACCTGGTCGTTAACAACCTGCTGATGGCCCTGGGCGAGCCGTTCTACG<br>CGGGTGTCCCGCTGATGGGCGAAGCGGCACGTCAGGCGTTTGGCCTGGACACCGAGTTCGCTCTGGAACCG<br>TTTAGCGAAAGCACGCTGGCACGTTTTGTTCGTCTGGAATGGCCGCACTTTATCCCAGCACCGGGCAAATCC<br>ATCGCGGACTGCTATGCCGCCATTCGTCAGGCGTTTTTGGATCTGCCGGACTTGTTTGGTGGCGAGGCAGGT<br>AAGCGTGGCGGTGAACACCACCTGTTCCTGAATGAGCTGACCAACCGTGCGCATCCGGGTTATCAACTGGA<br>AGTTTTCGATCGCGACTCGGCGCTGTTTGGTATTGCATTTGTGACCGATCAGGGCGAAGGTGGCGCTCTGGA<br>CAGCCCGCACTACGAACATAGCCATTTTCAACGTCTGCGTGAAATGAGCGCGCGTATCATGGCTCAAAGCGC<br>ACCGTTCGAACCGGCGCTGCCGGCGTTGCGTAATCCGGTTCTGGATGAGAGCCCGGGTTGCCAACGTGTCG<br>CAGACGGTCGTGCGCGTGCGCTGATGGCATTGTACCAAGGCGTTTATGAGCTGATGTTTGCGATGATGGCG<br>CAGCACTTCGCCGTGAAACCGCTGGGTAGCTTGCGTCGCAGCCGCCTGATGAACGCAGCAATCGATCTGAT<br>GACCGGTCTGTTGCGTCCGCTGAGCTGCGCGCTGATGAACCTGCCAAGCGGCATCGCCGGTCGCACGGCCG<br>GTCCGCCGCTGCCGGGTCCGGTTGACACCCGTAGCTATGACGACTACGCGCTGGGCTGTCGCATGCTGGCA<br>CGCCGTTGCGAGCGTCTGCTGGAGCAGGCGAGCATGCTGGAACCGGGTTGGCTGCCGGATGCGCAGATGG<br>AGCTGCTGGATTTCTATCGTCGCCAAATGCTGGACTTGGCGTGCGGCAAACTGA | |
| mScarlet-I | Gene | ATGGTTAGCAAAGGCGAGGCGGTTATCAAGGAGTTTATGCGTTTTAAGGTTCACATGGAGGGTAGCATGAA<br>TGGTCACGAGTTCGAGATCGAGGGTGAAGGCGAGGGTCGTCCGTACGAAGGCACCCAGACCGCGAAGCTG<br>AAAGTGACCAAGGGTGGCCCGCTGCCGTTCAGCTGGGACATCCTGAGCCCGCAGTTCATGTATGGCAGCCG<br>TGCGTTTATCAAACACCCGGCGGACATTCCGGATTACTATAAGCAAAGCTTCCCGGAAGGTTTTAAATGGGA<br>GCGTGTTATGAACTTCGAAGATGGTGGCGCGGTGACCGTTACCCAGGACACCAGCCTGGAGGATGGCACCC<br>TGATTTACAAGGTGAAACTGCGTGGCACCAACTTTCCGCCGGATGGTCCGGTTATGCAGAAGAAAACGATG<br>GGTTGGGAAGCGAGCACCGAGCGTCTGTATCCGGAAGATGGCGTGCTGAAGGGTGATATCAAAATGGCGC<br>TGCGTCTGAAGGACGGTGGCCGTTACCTGGCGGATTTTAAGACCACCTATAAAGCGAAGAAACCGGTGCAA<br>ATGCCGGGTGCGTACAACGTTGACCGTAAACTGGATATTACCAGCCACAACGAGGATTATACCGTGGTTGA<br>GCAATATGAGCGTAGCGAGGGTCGCCACAGCACCGGCGGCATGGACGAACTGTATAAGTGA | CIDAR MoClo Extension |
| mCherry | Gene | ATGGTGAGCAAGGGCGAGGAGGATAACATGGCCATCATCAAGGAGTTCATGCGCTTCAAGGTTCACATGGA<br>GGGCTCCGTGAACGGCCACGAGTTCGAGATCGAGGGCGAGGGCGAGGGCCGCCCCTACGAGGGCACCCAG<br>ACCGCCAAGCTGAAGGTGACCAAGGGTGGCCCCCTGCCCTTCGCCTGGGACATCCTGTCCCCTCAGTTCATG<br>TACGGCTCCAAGGCCTACGTGAAGCACCCCGCCGACATCCCCGACTACTTGAAGCTGTCCTTCCCCGAGGGC<br>TTCAAGTGGGAGCGCGTGATGAACTTCGAGGACGGCGGCGTGGTGACCGTGACCCAGGACTCCTCCCTGCA<br>AGACGGCGAGTTCATCTACAAGGTGAAGCTGCGCGGCACCAACTTCCCCTCCGACGGCCCCGTAATGCAGA<br>AGAAGACTATGGGCTGGGAGGCCTCCTCCGAGCGGATGTACCCCGAGGACGGCGCGCTGAAGGGCGAGAT<br>CAAGCAGAGGCTGAAGCTGAAGGACGGCGGCCACTACGACGCTGAGGTCAAGACCACCTACAAGGCCAAG<br>AAGCCCGTGCAACTGCCCGGCGCGTACAACGTCAACATCAAGTTGGACATCACCTCCCACAACGAGGACTAC<br>ACCATCGTGGAACAGTACGAACGCGCCGAGGGCCGCCACTCCACCGGCGGCATGGACGAGCTGTATAAGTA<br>A | 4 |
| mRFP1 | Gene | ATGGTGAGCAAGGGCGAGGAGGATAACATGGCCATCATCAAGGAGTTCATGCGCTTCAAGGTTCACATGGA<br>GGGCTCCGTGAACGGCCACGAGTTCGAGATCGAGGGCGAGGGCGAGGGCCGCCCCTACGAGGGCACCCAG<br>ACCGCCAAGCTGAAGGTGACCAAGGGTGGCCCCCTGCCCTTCGCCTGGGACATCCTGTCCCCTCAGTTCATG<br>TACGGCTCCAAGGCCTACGTGAAGCACCCCGCCGACATCCCCGACTACTTGAAGCTGTCCTTCCCCGAGGGC<br>TTCAAGTGGGAGCGCGTGATGAACTTCGAGGACGGCGGCGTGGTGACCGTGACCCAGGACTCCTCCCTGCA<br>AGACGGCGAGTTCATCTACAAGGTGAAGCTGCGCGGCACCAACTTCCCCTCCGACGGCCCCGTAATGCAGA<br>AGAAGACTATGGGCTGGGAGGCCTCCTCCGAGCGGATGTACCCCGAGGACGGCGCGCTGAAGGGCGAGAT<br>CAAGCAGAGGCTGAAGCTGAAGGACGGCGGCCACTACGACGCTGAGGTCAAGACCACCTACAAGGCCAAG<br>AAGCCCGTGCAACTGCCCGGCGCGTACAACGTCAACATCAAGTTGGACATCACCTCCCACAACGAGGACTAC<br>ACCATCGTGGAACAGTACGAACGCGCCGAGGGCCGCCACTCCACCGGCGGCATGGACGAGCTGTATAAGTA<br>A | 8 |
| mKate | Gene | ATGGAACTGATTAAAGAAAACATGCATATGAAACTGTATATGGAAGGCACCGTGAACAACCATCACTTTAAA<br>TGTACCAGCGAAGGTGAAGGTAAACCGTATGAAGGCACCCAGACCATGCGTATTAAAGCAGTTGAAGGTGG<br>TCCGCTGCCGTTTGCATTTGATATTCTGGCAACCAGCTTTATGTATGGCAGCAAAACCTTTATTAACCATACCC<br>AGGGTATCCCGGATTTTTTCAAACAGAGCTTTCCGGAAGGTTTTACCTGGGAACGTGTTACCACCTATGAAG<br>ATGGTGGTGTTCTGACCGCAACCCAGGATACCAGTCTGCAGGATGGTTGTCTGATTTATAATGTGAAAATTC<br>GCGGTGTGAACTTTCCGAGCAATGGTCCGGTTATGCAGAAAAAAACCCTGGGTTGGGAAGCAAGCACCGAA<br>ACCCTGTATCCGGCAGATGGTGGCCTGGAAGGTCGTGCAGATATGGCACTGAAACTGGTTGGTGGTGGTCA<br>TCTGATTTGCAATCTGAAAACCACCTATCGTAGCAAAAAACCTGCCAAAAACCTGAAAATGCCTGGCGTTTAT<br>TATGTTGATCGTCGTCTGGAACGTATCAAAGAGGCAGATAAAGAAACCTATGTGGAACAGCATGAAGTTGC<br>AGTTGCACGTTATTGTGATCTGCCGAGCAAACTGGGTCATCGTTAATAA | 1 |
| sfGFP | Gene | ATGCGTAAAGGCGAGGAGCTGTTCACTGGTGTCGTCCCTATTCTGGTGGAACTGGATGGTGATGTCAACGG<br>TCATAAGTTTTCCGTGCGTGGCGAGGGTGAAGGTGACGCAACTAATGGTAAACTGACGCTGAAGTTCATCT<br>GTACTACTGGTAAACTGCCGGTACCTTGGCCGACTCTGGTAACGACGCTGACTTATGGTGTTCAGTGCTTTGC<br>TCGTTATCCGGACCATATGAAGCAGCATGACTTCTTCAAGTCCGCCATGCCGGAAGGCTATGTGCAGGAACG<br>CACGATTTCCTTTAAGGATGACGGCACGTACAAAACGCGTGCGGAAGTGAAATTTGAAGGCGATACCCTGG<br>TAAACCGCATTGAGCTGAAAGGCATTGACTTTAAAGAAGACGGCAATATCCTGGGCCATAAGCTGGAATAC<br>AATTTTAACAGCCACAATGTTTACATCACCGCCGATAAACAAAAAAATGGCATTAAAGCGAATTTTAAAATTC<br>GCCACAACGTGGAGGATGGCAGCGTGCAGCTGGCTGATCACTACCAGCAAAACACTCCAATCGGTGATGGT | 13 |

| | | | |
|---|---|---|---|
| | | CCTGTTCTGCTGCCAGACAATCACTATCTGAGCACGCAAAGCGTTCTGTCTAAAGATCCGAACGAGAAACGC<br>GATCATATGGTTCTGCTGGAGTTCGTAACCGCAGCGGGCATCACGCATGGTATGGATGAACTGTACAAATAA | |
| sfYFP | Gene | ATGCGTAAAGGCGAAGAGCTGTTCACTGGTGTCGTCCCTATTCTGGTGGAACTGGATGGTGATGTCAACGG<br>TCATAAGTTTTCCGTGCGTGGCGAGGGTGAAGGTGACGCAACTAATGGTAAACTGACGCTGAAGTTCATCT<br>GTACTACTGGTAAACTGCCGGTACCTTGGCCGACTCTGGTAACGACGCTGACTTATGGTGTTCAGTGCTTTGC<br>TCGTTATCCGGACCATATGAAGCAGCATGACTTCTTCAAGTCCGCCATGCCGGAAGGCTATGTGCAGGAACG<br>CACGATTTCCTTTAAGGATGACGGCACGTACAAAACGCGTGCGGAAGTGAAATTTGAAGGCGATACCCTGG<br>TAAACCGCATTGAGCTGAAAGGCATTGACTTTAAAGAAGACGGCAATATCCTGGGCCATAAGCTGGAATAC<br>AATTTTAACAGCCACAATGTTTACATCACCGCCGATAAACAAAAAAATGGCATTAAAGCGAATTTTAAAATTC<br>GCCACAACGTGGAGGATGGCAGCGTGCAGCTGGCTGATCACTACCAGCAAAACACTCCAATCGGTGATGGT<br>CCTGTTCTGCTGCCAGACAATCACTATCTGAGCTACCAAAGCGTTCTGTCTAAAGATCCGAACGAGAAACGC<br>GATCATATGGTTCTGCTGGAGTTCGTAACCGCAGCGGGCATCACGCATGGTATGGATGAACTGTACAAATAA | CIDAR MoClo Extension |
| sfCFP | Gene | ATGCGTAAAGGCGAAGAGCTGTTCACTGGTGTCGTCCCTATTCTGGTGGAACTGGATGGTGATGTCAACGG<br>TCATAAGTTTTCCGTGCGTGGCGAGGGTGAAGGTGACGCAACTAATGGTAAACTGACGCTGAAGTTCATCT<br>GTACTACTGGTAAACTGCCGGTACCTTGGCCGACTCTGGTAACGACGCTGACTTGGGGTGTTCAGTGCTTTG<br>CTCGTTATCCGGACCATATGAAGCAGCATGACTTCTTCAAGTCCGCCATGCCGGAAGGCTATGTGCAGGAAC<br>GCACGATTTCCTTTAAGGATGACGGCACGTACAAAACGCGTGCGGAAGTGAAATTTGAAGGCGATACCCTG<br>GTAAACCGCATTGAGCTGAAAGGCATTGACTTTAAAGAAGACGGCAATATCCTGGGCCATAAGCTGGAATA<br>CAATTACATCAGCGACAATGTTTACATCACCGCCGATAAACAAAAAAATGGCATTAAAGCGAATTTTAAAATT<br>CGCCACAACGTGGAGGATGGCAGCGTGCAGCTGGCTGATCACTACCAGCAAAACACTCCAATCGGTGATGG<br>TCCTGTTCTGCTGCCAGACAATCACTATCTGAGCACGCAAAGCGTTCTGTCTAAAGATCCGAACGAGAAACG<br>CGATCATATGGTTCTGCTGGAGTTCGTAACCGCAGCGGGCATCACGCATGGTATGGATGAACTGTACAAATA<br>A | CIDAR MoClo Extension |
| CtrE | Gene | ATGACGGTCTGCGCAAAAAAACACGTTCATCTCACTCGCGATGCTGCGGAGCAGTTACTGGCTGATATTGAT<br>CGACGCCTTGATCAGTTATTGCCCGTGGAGGGAGAACGGGATGTTGTGGGTGCCGCGATGCGTGAAGGTG<br>CGCTGGCACCGGGAAAACGTATTCGCCCCATGTTGCTGTTGCTGACCGCCCGCGATCTGGGTTGCGCTGTCA<br>GCCATGACGGATTACTGGATTTGGCCTGTGCGGTGGAAATGGTCCACGCGGCTTCGCTGATCCTTGACGATA<br>TGCCCTGCATGGACGATGCGAAGCTGCGGCGCGGACGCCCTACCATTCATTCTCATTACGGAGAGCATGTG<br>GCAATACTGGCGGCGGTTGCCTTGCTGAGTAAAGCCTTTGGCGTAATTGCCGATGCAGATGGCCTCACGCCG<br>CTGGCAAAAAATCGGGCGGTTTCTGAACTGTCAAACGCCATCGGCATGCAAGGATTGGTTCAGGGTCAGTT<br>CAAGGATCTGTCTGAAGGGGATAAGCCGCGCAGCGCTGAAGCTATTTTGATGACGAATCACTTTAAAACCA<br>GCACGCTGTTTTGTGCCTCCATGCAGATGGCCTCGATTGTTGCGAATGCCTCCAGCGAAGCGCGTGATTGCC<br>TGCATCGTTTTTCACTTGATCTTGGTCAGGCATTTCAACTGCTGGACGATTTGACCGATGGCATGACCGACAC<br>CGGTAAGGATAGCAATCAGGACGCCGGTAAATCGACGCTGGTCAATCTGTTAGGCCCGAGGGCGGTTGAA<br>GAACGTCTGAGACAACATCTTCAGCTTGCCAGTGAGCATCTCTCTGCGGCCTGCCAACACGGGCACGCCACT<br>CAACATTTTATTCAGGCCTGGTTTGACAAAAAACTCGCTGCCGTCAGTTAA | 1 |
| CtrB | Gene | ATGAATAATCCGTCGTTACTCAATCATGCGGTCGAAACGATGGCAGTTGGCTCGAAAAGTTTTGCGACAGCC<br>TCAAAGTTATTTGATGCAAAAACCCGGCGCAGCGTACTGATGCTCTACGCCTGGTGCCGCCATTGTGACGAT<br>GTTATTGACGATCAGACGCTGGGCTTTCAGGCCCGGCAGCCTGCCTTACAAACGCCCGAACAACGTCTGATG<br>CAACTTGAGATGAAAACGCGCCAGGCCTATGCAGGATCGCAGATGCACGAACCGGCGTTTGCGGCTTTTCA<br>GGAAGTGGCTATGGCTCATGATATCGCCCCGGCTTACGCGTTTGATCATCTGGAAGGCTTCGCCATGGATGT<br>ACGCGAAGCGCAATACAGCCAACTGGATGATACGCTGCGCTATTGCTATCACGTTGCAGGCGTTGTCGGCTT<br>GATGATGGCGCAAATCATGGGCGTGCGGGATAACGCCACGCTGGACCGCGCCTGTGACCTTGGGCTGGCAT<br>TTCAGTTGACCAATATTGCTCGCGATATTGTGGACGATGCGCATGCGGGCCGCTGTTATCTGCCGGCAAGCT<br>GGCTGGAGCATGAAGGTCTGAACAAAGAGAATTATGCGGCACCTGAAAACCGTCAGGCGCTGAGCCGTATC<br>GCCCGTCGTTTGGTGCAGGAAGCAGAACCTTACTATTTGTCTGCCACAGCCGGCCTGGCAGGGTTGCCCCTG<br>CGTTCCGCCTGGGCAATCGCTACGGCGAAGCAGGTTTACCGGAAAATAGGTGTCAAAGTTGAACAGGCCGG<br>TCAGCAAGCCTGGGATCAGCGGCAGTCAACGACCACGCCCGAAAAATTAACGCTGCTGCTGGCCGCCTCTG<br>GTCAGGCCCTTACTTCCCGGATGCGGGCTCATCCTCCCCGCCCTGCGCATCTCTGGCAGCGCCCGCTCTAG | 1 |
| CtrI | Gene | ATGAAACCAACTACGGTAATTGGTGCAGGCTTCGGTGGCCTGGCACTGGCAATTCGTCTACAAGCTGCGGG<br>GATCCCCGTCTTACTGCTTGAACAACGTGATAAACCCGGCGGTCGGGCTTATGTCTACGAGGATCAGGGGTT<br>TACCTTTGATGCAGGCCCGACGGTTATCACCGATCCCAGTGCCATTGAAGAACTGTTTGCACTGGCAGGAAA<br>ACAGTTAAAAGAGTATGTCGAACTGCTGCCGGTTACGCCGTTTTACCGCCTGTGTTGGGAGTCAGGGAAGG<br>TCTTTAATTACGATAACGATCAAACCCGGCTCGAAGCGCAGATTCAGCAGTTTAATCCCCGCGATGTCGAAG<br>GTTATCGTCAGTTTCTGGACTATTCACGCGCGGTGTTTAAAGAAGGCTATCTAAAGCTCGGTACTGTCCCTTT<br>TTTATCGTTCAGAGACATGCTTCGCGCCGCACCTCAACTGGCGAAACTGCAGGCATGGAGAAGCGTTTACAG<br>TAAGGTTGCCAGTTACATCGAAGATGAACATCTGCGCCAGGCGTTTTCTTTCCACTCGCTGTTGGTGGGCGG<br>CAATCCCTTCGCCACCTCATCCATTTATACGTTGATACACGCGCTGGAGCGTGAGTGGGGCGTCTGGTTTCCG<br>CGTGGCGGCACCGGCGCATTAGTTCAGGGGATGATAAAGCTGTTTCAGGATCTGGGTGGCGAAGTCGTGTT<br>AAACGCCAGAGTCAGCCATATGGAAACGACAGGAAACAAGATTGAAGCCGTGCATTTAGAGGACGGTCGC<br>AGGTTCCTGACGCAAGCCGTCGCGTCAAATGCAGATGTGGTTCATACCTATCGCGACCTGTTAAGCCAGCAC<br>CCTGCCGCGGTTAAGCAGTCCAACAAACTGCAGACTAAGCGCATGAGTAACTCTCTGTTTGTGCTCTATTTTG<br>GTTTGAATCACCATCATGATCAGCTCGCGCATCACACGGTTTGTTTCGGCCCGCGTTACCGCGAGCTGATTGA<br>CGAAATTTTTAATCATGATGGCCTCGCAGAGGACTTCTCACTTTATCTGCACGCGCCCTGTGTCACGGATTCG<br>TCACTGGCGCCTGAAGGTTGCGGCAGTTACTATGTGTTGGCGCCGGTGCCGCATTTAGGCACCGCGAACCTC<br>GACTGGACGGTTGAGGGGCCAAAACTACGCGACCGTATTTTTGCGTACCTTGAGCAGCATTACATGCCTGGC<br>TTACGGAGTCAGCTGGTCACGCACCGGATGTTTACGCCGTTTGATTTTCGCGACCAGCTTAATGCCTATCATG<br>GCTCAGCCTTTTCTGTGGAGCCCGTTCTTACCCAGAGCGCCTGGTTTCGGCCGCATAACCGCGATAAAACCAT<br>TACTAATCTCTACCTGGTCGGCGCAGGCACGCATCCCGGCGCAGGCATTCCTGGCGTCATCGGCTCGGCAAA<br>AGCGACAGCAGGTTTGATGCTGGAGGATCTGATATGA | 1 |
| CtrY | Gene | ATGCAACCGCATTATGATCTGATTCTCGTGGGGGCTGGACTCGCGAATGGCCTTATCGCCCTGCGACTTCAG<br>CAGCAGCAACCTGATATGCGTATTTTGCTTATCGACGCCGCACCCCAGGCGGGCGGGAATCATACGTGGTCA<br>TTTCACCACGATGATTTGACTGAGAGCCAACATCGTTGGATAGCTCCGCTGGTGGTTCATCACTGGCCCGACT<br>ATCAGGTACGCTTTCCCACACGCCGTCGTAAGCTGAACAGCGGCTACTTTTGTATTACTTCTCAGCGTTTCGC<br>TGAGGTTTTACAGCGACAGTTTGGCCCGCACTTGTGGATGGATACCGCGGTCGCAGAGGTTAATGCGGAAT<br>CTGTTCGGTTGAAAAAGGGTCAGGTTATCGGTGCCCGCGCGGTGATTGACGGGCGGGGTTATGCGGCAAAT<br>TCAGCACTGAGCGTGGGCTTCCAGGCGTTTATTGGCCAGGAATGGCGATTGAGCCACCCGCATGGTTTATCA<br>TCTCCCATTATCATGGATGCCACGGTCGATCAGCAAAATGGTTATCGCTTCGTGTACAGCCTGCCGCTCTCGC<br>CGACCAGATTGTTAATTGAAGATACGCACTATATTGATAATGCGACATTAGATCCTGAATGCGCGCGGCAAA<br>ATATTTGCGACTATGCCGCGCAACAGGGTTGGCAGCTTCAGACACTGCTGCGAGAAGAACAGGGCGCCTTA<br>CCCATTACTCTGTCGGGCAATGCCGACGCATTCTGGCAGCAGCGCCCCCTGGCCTGTAGTGGATTACGTGCC<br>GGTCTGTTCCATCCTACCACCGGCTATTCACTGCCGCTGGCGGTTGCCGTGGCCGACCGCCTGAGTGCACTT<br>GATGTCTTTACGTCGGCCTCAATTCACCATGCCATTACGCATTTTGCCCGCGAGCGCTGGCAGCAGCAGGGC<br>TTTTTCCGCATGCTGAATCGCATGCTGTTTTTAGCCGGACCCGCCGATTCACGCTGGCGGGTTATGCAGCGTT<br>TTTATGGTTTACCTGAAGATTTAATTGCCCGTTTTTATGCGGGAAAACTCACGCTGACCGATCGGCTACGTAT<br>TCTGAGCGGCAAGCCGCCTGTTCCGGTATTAGCAGCATTGCAAGCCATTATGACGACTCATCGTTAA | 1 |

| | | | |
|---|---|---|---|
| dBroccoli | RNA Aptamer | TTGCCATGTGTATGTGGGAGACGGTCGGGTCCATCTGAGACGGTCGGGTCCAGATATTCGTATCTGTCGAGTAGAGTGTGGGCTCAGATGTCGAGTAGAGTGTGGGCTCCCACATACTCTGATGATCCAGACGGTCGGGTCCATCTGAGACGGTCGGGTCCAGATATTCGTATCTGTCGAGTAGAGTGTGGGCTCAGATGTCGAGTAGAGTGTGGGCTGGATCATTCATGGCAA | 15 |
| STAR | Regulatory RNA | TGAACTGTATACATTCCCCGCTGAACGACGGAAACTTTGACTGGACTGACTTGATGACTGG | 8 |
| STAR toehold 0 | Regulatory RNA | TGAACTGTATACATTCCCCGCTGAACGACGGAAACTTTGACTGGACTGACTTGATGACTGGTAACTCCATTCCATC | This study |
| STAR toehold 1 | Regulatory RNA | TGAACTGTATACATTCCCCGCTGAACGACGGAAACTTTGACTGGACTGACTTGATGACTGGTCTTATCTTATCTA | This study |
| STAR toehold 2 | Regulatory RNA | TGAACTGTATACATTCCCCGCTGAACGACGGAAACTTTGACTGGACTGACTTGATGACTGGAGTTTGATTACATT | This study |
| STAR toehold 3 | Regulatory RNA | TGAACTGTATACATTCCCCGCTGAACGACGGAAACTTTGACTGGACTGACTTGATGACTGGATCTATTACTACTT | This study |
| STAR toehold 4 | Regulatory RNA | TGAACTGTATACATTCCCCGCTGAACGACGGAAACTTTGACTGGACTGACTTGATGACTGGCGATTATGGATTAG | This study |
| STAR toehold 5 | Regulatory RNA | TGAACTGTATACATTCCCCGCTGAACGACGGAAACTTTGACTGGACTGACTTGATGACTGGTATGTAATTGATTT | This study |
| Anti-STAR | Regulatory RNA | CCAGTCATCAAGTCAGTCCAGTCAAAGTTTCCGTCGTTCAGCGGGGAATGTATACAGTTCA | This study |
| Anti-STAR toehold 0 | Regulatory RNA | GATGGAATGGAGTTACCAGTCATCAAGTCAGTCCAGTCAAAGTTTCCGTCGTTCAGCGGGGAATGTATACAGTTCA | This study |
| Anti-STAR toehold 1 | Regulatory RNA | TAGATAAGATAAGACCAGTCATCAAGTCAGTCCAGTCAAAGTTTCCGTCGTTCAGCGGGGAATGTATACAGTTCA | This study |
| Anti-STAR toehold 2 | Regulatory RNA | AATGTAATCAAACTCCAGTCATCAAGTCAGTCCAGTCAAAGTTTCCGTCGTTCAGCGGGGAATGTATACAGTTCA | This study |
| Anti-STAR toehold 3 | Regulatory RNA | AAGTAGTAATAGATCCAGTCATCAAGTCAGTCCAGTCAAAGTTTCCGTCGTTCAGCGGGGAATGTATACAGTTCA | This study |
| Anti-STAR toehold 4 | Regulatory RNA | CTAATCCATAATCGCCAGTCATCAAGTCAGTCCAGTCAAAGTTTCCGTCGTTCAGCGGGGAATGTATACAGTTCA | This study |
| Anti-STAR toehold 5 | Regulatory RNA | AAATCAATTACATACCAGTCATCAAGTCAGTCCAGTCAAAGTTTCCGTCGTTCAGCGGGGAATGTATACAGTTCA | This study |
| STAR Target | Regulatory RNA | CCAGTCATCAAGTCAGTCCAGTCAAAGTTTCCGTCGTTCAGCGGGGAATGTATACAGTTCATGTATATATTCCCCGCTTTTTTTTTT | 8 |
| Deoptimised STAR Target | Regulatory RNA | CCAGTCATCAAGTCAGTCCAGTCAAAGTTTCCCTGGTTCAGCGGGGAATGTATACAGTTCATGTATATATTCCCCGCTTTTTTTTTT | This study |
| Buffer 3 | Non-coding region | GGATCCTTACTCGAGAAAAAAAAAACCCCGCTTCGGCGGGGTTTTTTTTTTCTGGACTGCAGGCTTCCTCGCTCAC | This study |

### Supplementary Table 3: Plasmids used in this study

| Figure | Plasmid | Description | Source |
|---|---|---|---|
| Figure 2D | pJBL5939 | J23119 – STAR Target – $RBS_{RFP}$ – mRFP1 – TrrnB – p15A – CamR | [8] |
| | pAB317 | « $LLL_1$ » : $P_{tet}$ – B0034 – luxR – B0015 – AraC – $P_{araBAD}$ – STAR – t500 – Buffer 3 – $P_{lux}$ – anti-STAR – t500 – ColE1 – SpecR | This study |
| | pAB300 | « $LLL_2$ » : $P_{tet}$ – B0034 – luxR – B0015 – AraC – $P_{araBAD}$ – STAR_toehold2 – t500 – Buffer 3 – $P_{lux}$ – anti-STAR_toehold2 – t500 – ColE1 – SpecR | This study |
| | pAB545 | « $LLL_3$ » : $P_{tet}$ – B0030 – luxR – B0015 – AraC – $P_{araBAD}$ – STAR_toehold2 – t500 – Buffer 3 – $P_{lux}$ – anti-STAR_toehold2 – t500 – ColE1 – SpecR | This study |
| Figure 2E-F | pJBL5939 | J23119 – STAR Target – $RBS_{RFP}$ – mRFP1 – TrrnB – p15A – CamR | [8] |
| | pAB300 | « $LLL_2$ » : $P_{tet}$ – B0034 – luxR – B0015 – AraC – $P_{araBAD}$ – STAR_toehold2 – t500 – Buffer 3 – $P_{lux}$ – anti-STAR_toehold2 – t500 – ColE1 – SpecR | This study |
| | pAB401 | « LLR » : $P_{tet}$ – B0034 – rpaR – B0015 – AraC – $P_{araBAD}$ – STAR_toehold2 – t500 – Buffer 3 – $P_{lux}$ – anti-STAR_toehold2 – t500 – ColE1 – SpecR | This study |
| Figure 3B | pAB300 | « $LLL_2$ » : $P_{tet}$ – B0034 – luxR – B0015 – AraC – $P_{araBAD}$ – STAR_toehold2 – t500 – Buffer 3 – $P_{lux}$ – anti-STAR_toehold2 – t500 – ColE1 – SpecR | This study |
| | pAB368 | J23119 – STAR Target – $RBS_{Bujard}$ – eforRed – TrrnB – p15A – CmR | This study |
| Figure 3C | pAB300 | « $LLL_2$ » : $P_{tet}$ – B0034 – luxR – B0015 – AraC – $P_{araBAD}$ – STAR_toehold2 – t500 – Buffer 3 – $P_{lux}$ – anti-STAR_toehold2 – t500 – ColE1 – SpecR | This study |
| | pAB517 | J23100 – STAR Target – $RBS_{Bujard}$ – VioB-mCherry – L3S2P55 – p15A – CamR | This study |
| Figure 3D | pAB401 | $P_{tet}$ – B0034 – rpaR – B0015 – AraC – $P_{araBAD}$ – STAR_toehold2 – t500 – Buffer 3 – $P_{lux}$ – anti-STAR_toehold2 – t500 – ColE1 – SpecR | This study |
| | pAB550 | J23114 – STAR Target – $RBS_{CtrE}$ – CtrE – $RBS_{CtrB}$ – CtrB – $RBS_{CtrI}$ – CtrI – $RBS_{CtrY}$ – CtrY – L3S2P55 – p15A – CamR | This study |
| Figure 3E | pAB300 | « $LLL_2$ » : $P_{tet}$ – B0034 – luxR – B0015 – AraC – $P_{araBAD}$ – STAR_toehold2 – t500 – Buffer 3 – $P_{lux}$ – anti-STAR_toehold2 – t500 – ColE1 – SpecR | This study |
| | pAB205 | $P_{RAND14}$ – RBS8B – dCas9 – rrnB T1 – J23119 – STAR Target 6 – HH – gRNA(LeuLp, End-0: A→C) – HDV – p15A – AmpR | This study |
| Figure 4B-C | pAB317 | « $LLL_1$ » : $P_{tet}$ – B0034 – luxR – B0015 – AraC – $P_{araBAD}$ – STAR – t500 – Buffer 3 – $P_{lux}$ – anti-STAR – t500 – ColE1 – SpecR | This study |
| | pAB537 | $P_{tet}$ – B0034 – rpaR – B0015 – AraC – $P_{rhaBAD}$ – STAR_toehold2 – t500 – Buffer 3 – $P_{lux}$ – anti-STAR_toehold2 – t500 – ColE1 – SpecR | This study |
| | pAB518 | J23100 – STAR Target 6 – $RBS_{Bujard}$ – VioB-sfYFP – L3S2P55 – p15A – CamR | This study |
| | pAB519 | J23100 – STAR Target 6 – $RBS_{Bujard}$ – VioB-sfCFP – L3S2P55 – p15A – CamR | This study |
| Figures 5B1-2<br>Figure 5C | pAB300 | « $LLL_2$ » : $P_{tet}$ – B0034 – luxR – B0015 – AraC – $P_{araBAD}$ – STAR_toehold2 – t500 – Buffer 3 – $P_{lux}$ – anti-STAR_toehold2 – t500 – ColE1 – SpecR | This study |
| | pAB537 | $P_{tet}$ – B0034 – rpaR – B0015 – AraC – $P_{rhaBAD}$ – STAR_toehold2 – t500 – Buffer 3 – $P_{lux}$ – anti-STAR_toehold2 – t500 – ColE1 – SpecR | This study |
| | pAB518 | J23100 – STAR Target 6 – $RBS_{Bujard}$ – VioB-sfYFP – L3S2P55 – p15A – CamR | This study |
| | pAB519 | J23100 – STAR Target 6 – $RBS_{Bujard}$ – VioB-sfCFP – L3S2P55 – p15A – CamR | This study |
| Figure 5B3<br>Figure 5C | pAB317 | « $LLL_1$ » : $P_{tet}$ – B0034 – luxR – B0015 – AraC – $P_{araBAD}$ – STAR – t500 – Buffer 3 – $P_{lux}$ – anti-STAR – t500 – ColE1 – SpecR | This study |
| | pAB537 | $P_{tet}$ – B0034 – rpaR – B0015 – AraC – $P_{rhaBAD}$ – STAR_toehold2 – t500 – Buffer 3 – $P_{lux}$ – anti-STAR_toehold2 – t500 – ColE1 – SpecR | This study |
| | pAB518 | J23100 – STAR Target 6 – $RBS_{Bujard}$ – VioB-sfYFP – L3S2P55 – p15A – CamR | This study |
| | pAB519 | J23100 – STAR Target 6 – $RBS_{Bujard}$ – VioB-sfCFP – L3S2P55 – p15A – CamR | This study |
| Figure 5B4<br>Figure 5C<br>Figure 5D<br>Figure 5E<br>Figure 5F | pAB317 | « $LLL_1$ » : $P_{tet}$ – B0034 – luxR – B0015 – AraC – $P_{araBAD}$ – STAR – t500 – Buffer 3 – $P_{lux}$ – anti-STAR – t500 – ColE1 – SpecR | This study |
| | pAB401 | « LLR » : $P_{tet}$ – B0034 – rpaR – B0015 – AraC – $P_{araBAD}$ – STAR_toehold2 – t500 – Buffer 3 – $P_{lux}$ – anti-STAR_toehold2 – t500 – ColE1 – SpecR | This study |
| | pAB518 | J23100 – STAR Target 6 – $RBS_{Bujard}$ – VioB-sfYFP – L3S2P55 – p15A – CamR | This study |
| | pAB519 | J23100 – STAR Target 6 – $RBS_{Bujard}$ – VioB-sfCFP – L3S2P55 – p15A – CamR | This study |
| Supplementary Figure 2B-D | pAB420 | $P_{tet}$ – B0034 – luxR – L3S2P55 – $P_{lux}$ – dBroccoli – L3S2P21 – ColE1 – SpecR | This study |
| Supplementary Figure 3C-D | pAB420 | « Lux Receiver » : $P_{tet}$ – B0034 – luxR – L3S2P55 – $P_{lux}$ – dBroccoli – L3S2P21 – ColE1 – SpecR | This study |

| | | | |
|---|---|---|---|
| | pAB421 | « Rpa Receiver » : $P_{tet}$ – B0034 – luxR – L3S2P55 – $P_{lux}$ – dBroccoli – L3S2P21 – ColE1 – SpecR | This study |
| Supplementary Figure 4B-D | pJBL5939 | J23119 – STAR Target – $RBS_{RFP}$ – mRFP1 – TrrnB – p15A – CamR | [8] |
| | pAB161 | $P_{tet}$ – B0034 – luxR – B0015 – J23100 – STAR toehold0 – t500 – $P_{lux}$ – anti-STAR toehold0 – t500 – ColE1 – SpecR | This study |
| | pAB232 | $P_{tet}$ – B0034 – luxR – B0015 – J23100 – STAR toehold1 – t500 – $P_{lux}$ – anti-STAR toehold1 – t500 – ColE1 – SpecR | This study |
| | pAB233 | $P_{tet}$ – B0034 – luxR – B0015 – J23100 – STAR toehold2 – t500 – $P_{lux}$ – anti-STAR toehold2 – t500 – ColE1 – SpecR | This study |
| | pAB234 | $P_{tet}$ – B0034 – luxR – B0015 – J23100 – STAR toehold3 – t500 – $P_{lux}$ – anti-STAR toehold3– t500 – ColE1 – SpecR | This study |
| | pAB235 | $P_{tet}$ – B0034 – luxR – B0015 – J23100 – STAR toehold4– t500 – $P_{lux}$ – anti-STAR toehold4 – t500 – ColE1 – SpecR | This study |
| | pAB236 | $P_{tet}$ – B0034 – luxR – B0015 – J23100 – STAR toehold5 – t500 – $P_{lux}$ – anti-STAR toehold5 – t500 – ColE1 – SpecR | This study |
| Supplementary Figure 5B | pJBL5939 | J23119 – STAR Target – $RBS_{RFP}$ – mRFP1 – TrrnB – p15A – CamR | [8] |
| | pAB300 | « $LLL_2$ » : $P_{tet}$ – B0034 – luxR – B0015 – AraC – $P_{araBAD}$ – STAR_toehold2 – t500 – Buffer 3 – $P_{lux}$ – anti-STAR_toehold2 – t500 – ColE1 – SpecR | This study |
| | pAB317 | « $LLL_1$ » : $P_{tet}$ – B0034 – luxR – B0015 – AraC – $P_{araBAD}$ – STAR – t500 – Buffer 3 – $P_{lux}$ – anti-STAR – t500 – ColE1 – SpecR | This study |
| Supplementary Figure 5C | pJBL5939 | J23119 – STAR Target – $RBS_{RFP}$ – mRFP1 – TrrnB – p15A – CamR | [8] |
| | pAB271 | AraC – $P_{araBAD}$ – STAR toehold2 – t500 – ColE1 – SpecR | This study |
| | pAB298 | $P_{tet}$ – B0034 – luxR – B0015 – $P_{lux}$ – anti-STAR toehold2 – t500 – ColE1 – SpecR | This study |
| | pAB300 | « $LLL_2$ » : $P_{tet}$ – B0034 – luxR – B0015 – AraC – $P_{araBAD}$ – STAR_toehold2 – t500 – Buffer 3 – $P_{lux}$ – anti-STAR_toehold2 – t500 – ColE1 – SpecR | This study |
| | pAB317 | « $LLL_1$ » : $P_{tet}$ – B0034 – luxR – B0015 – AraC – $P_{araBAD}$ – STAR – t500 – Buffer 3 – $P_{lux}$ – anti-STAR – t500 – ColE1 – SpecR | This study |
| | pAB303 | AraC – $P_{araBAD}$ – STAR – t500 – ColE1 – SpecR | This study |
| | pAB304 | $P_{tet}$ – B0034 – luxR – B0015 – $P_{lux}$ – anti-STAR – t500 – ColE1 – SpecR | This study |
| Supplementary Figure 6B | pJBL5939 | J23119 – STAR Target – $RBS_{RFP}$ – mRFP1 – TrrnB – p15A – CamR | [8] |
| | pAB262 | J23119 – STAR Target – STAR Target – $RBS_{RFP}$ – mRFP1 – TrrnB – p15A – CamR | This study |
| | pAB127 | $P_{tet}$ – B0034 – luxR – B0015 – $P_{lux}$ – STAR toehold0 – t500 – ColE1 – SpecR | This study |
| Supplementary Figure 6B | pJBL5939 | J23119 – STAR Target – $RBS_{RFP}$ – mRFP1 – TrrnB – p15A – CamR | [8] |
| | pAB262 | J23119 – STAR Target – STAR Target – $RBS_{RFP}$ – mRFP1 – TrrnB – p15A – CamR | This study |
| | pAB161 | $P_{tet}$ – B0034 – luxR – B0015 – J23100 – STAR toehold0 – t500 – $P_{lux}$ – anti-STAR toehold0 – t500 – ColE1 – SpecR | This study |
| Supplementary Figure 7B | pJBL5939 | J23119 – STAR Target – $RBS_{RFP}$ – mRFP1 – TrrnB – p15A – CamR | [8] |
| | pAB300 | « $LLL_2$ » : $P_{tet}$ – B0034 – luxR – B0015 – AraC – $P_{araBAD}$ – STAR_toehold2 – t500 – Buffer 3 – $P_{lux}$ – anti-STAR_toehold2 – t500 – ColE1 – SpecR | This study |
| | pAB401 | « LLR » : $P_{tet}$ – B0034 – rpaR – B0015 – AraC – $P_{araBAD}$ – STAR_toehold2 – t500 – Buffer 3 – $P_{lux}$ – anti-STAR_toehold2 – t500 – ColE1 – SpecR | This study |
| Supplementary Figure 8B-C | pAB300 | « $LLL_2$ » : $P_{tet}$ – B0034 – luxR – B0015 – AraC – $P_{araBAD}$ – STAR_toehold2 – t500 – Buffer 3 – $P_{lux}$ – anti-STAR_toehold2 – t500 – ColE1 – SpecR | This study |
| | pAB317 | « $LLL_1$ » : $P_{tet}$ – B0034 – luxR – B0015 – AraC – $P_{araBAD}$ – STAR – t500 – Buffer 3 – $P_{lux}$ – anti-STAR – t500 – ColE1 – SpecR | This study |
| Supplementary Figure 9B-D | pAB300 | « $LLL_2$ » : $P_{tet}$ – B0034 – luxR – B0015 – AraC – $P_{araBAD}$ – STAR_toehold2 – t500 – Buffer 3 – $P_{lux}$ – anti-STAR_toehold2 – t500 – ColE1 – SpecR | This study |
| | pAB517 | J23100 – STAR Target – $RBS_{Bujard}$ – VioB-mCherry – L3S2P55 – p15A – CamR | This study |
| Supplementary Figure 10B-E | pAB401 | « LLR » : $P_{tet}$ – B0034 – rpaR – B0015 – AraC – $P_{araBAD}$ – STAR_toehold2 – t500 – Buffer 3 – $P_{lux}$ – anti-STAR_toehold2 – t500 – ColE1 – SpecR | This study |
| | pAB550 | J23114 – STAR Target – $RBS_{CtrE}$ – CtrE – $RBS_{CtrB}$ – CtrB – $RBS_{CtrI}$ – CtrI – $RBS_{CtrY}$ – CtrY – L3S2P55 – p15A – CamR | This study |
| | B0034_mKate | J23106 – B0034 – mKate – B0015 – pMB1 – CamR | [1] |

| | | | |
|---|---|---|---|
| Supplementary Figure 11E | pAB81 | $P_{RAND14}$ – RBS8B – dCas9 – rrnB T1 – $P_{araBAD}$ – HH – gRNA(LeuLp) – HDV – p15A – AmpR | This study |
| Supplementary Figure 12 | pAB58 | $P_{RAND14}$ – RBS8B – dCas9 – rrnB T1 – AraC - $P_{araBAD}$ – No Target gRNA – p15A – AmpR | This study |
| | pAB60 | $P_{RAND14}$ – RBS8B – dCas9 – rrnB T1 – AraC - $P_{araBAD}$ – gRNA(HisLp) – p15A – AmpR | This study |
| | pAB61 | $P_{RAND14}$ – RBS8B – dCas9 – rrnB T1 – AraC – $P_{araBAD}$ – gRNA(LeuLp) – p15A – AmpR | This study |
| Supplementary Figure 13 | pAB58 | $P_{RAND14}$ – RBS8B – dCas9 – rrnB T1 – AraC - $P_{araBAD}$ – No Target gRNA – p15A – AmpR | This study |
| | pAB96 | $P_{RAND14}$ – RBS8B – dCas9 – rrnB T1 – J23119 – STAR Target 6 – HH – gRNA(LeuLp) – HDV – p15A – AmpR | This study |
| | pAB205 | $P_{RAND14}$ – RBS8B – dCas9 – rrnB T1 – J23119 – STAR Target 6 – HH – gRNA(LeuLp, End-0: A→C) – HDV – p15A – AmpR | This study |
| Supplementary Figure 14 | pAB96 | $P_{RAND14}$ – RBS8B – dCas9 – rrnB T1 – J23119 – STAR Target 6 – HH – gRNA(LeuLp) – HDV – p15A – AmpR | This study |
| | pAB205 | $P_{RAND14}$ – RBS8B – dCas9 – rrnB T1 – J23119 – STAR Target 6 – HH – gRNA(LeuLp, End-0 mutation) – HDV – p15A – AmpR | This study |
| | pAB206 | $P_{RAND14}$ – RBS8B – dCas9 – rrnB T1 – J23119 – STAR Target 6 – HH – gRNA(LeuLp, End-1 mutation) – HDV – p15A – AmpR | This study |
| | pAB208 | $P_{RAND14}$ – RBS8B – dCas9 – rrnB T1 – J23119 – STAR Target 6 – HH – gRNA(LeuLp, End-3 mutation) – HDV – p15A – AmpR | This study |
| | pAB127 | $P_{tet}$ – B0034 – luxR – B0015 – AraC – $P_{araBAD}$ – STAR_toehold0 – t500 – SpecR | This study |
| Supplementary Figure 16 | pAB252 | $P_{tet}$ – B0034 – luxR – L3S2P55 – $P_{lux}$ – RBSc33 – sfGFP – L3S2P21 – ColE1 – SpecR | This study |
| | pAB409 | AraC – $P_{araBAD}$ – RBSc33 – sfGFP – L3S2P21 – ColE1 – SpecR | This study |
| | pAB410 | $P_{rhaBAD}$ – RBSc33 – sfGFP – L3S2P21 – ColE1 – SpecR | This study |
| Supplementary Figure 17B-C | pAB300 | « $LLL_2$ » : $P_{tet}$ – B0034 – luxR – B0015 – AraC – $P_{araBAD}$ – STAR_toehold2 – t500 – Buffer 3 – $P_{lux}$ – anti-STAR_toehold2 – t500 – ColE1 – SpecR | This study |
| | pAB518 | J23100 – STAR Target 6 – $RBS_{Bujard}$ – VioB-sfYFP – L3S2P55 – p15A – CamR | This study |
| Supplementary Figure 17E-F | pAB401 | « LLR » : $P_{tet}$ – B0034 – rpaR – B0015 – AraC – $P_{araBAD}$ – STAR_toehold2 – t500 – Buffer 3 – $P_{lux}$ – anti-STAR_toehold2 – t500 – ColE1 – SpecR | This study |
| | pAB519 | J23100 – STAR Target 6 – $RBS_{Bujard}$ – VioB-sfCFP – L3S2P55 – p15A – CamR | This study |
| Supplementary Figure 18B-F | pAB300 | « $LLL_2$ » : $P_{tet}$ – B0034 – luxR – B0015 – AraC – $P_{araBAD}$ – STAR_toehold2 – t500 – Buffer 3 – $P_{lux}$ – anti-STAR_toehold2 – t500 – ColE1 – SpecR | This study |
| | pAB537 | $P_{tet}$ – B0034 – rpaR – B0015 – AraC – $P_{rhaBAD}$ – STAR_toehold2 – t500 – Buffer 3 – $P_{lux}$ – anti-STAR_toehold2 – t500 – ColE1 – SpecR | This study |
| | pAB518 | J23100 – STAR Target 6 – $RBS_{Bujard}$ – VioB-sfYFP – L3S2P55 – p15A – CamR | This study |
| | pAB519 | J23100 – STAR Target 6 – $RBS_{Bujard}$ – VioB-sfCFP – L3S2P55 – p15A – CamR | This study |
| Supplementary Figure 19B-E | pAB399 | $P_{tet}$ – B0034 – luxR – B0015 – $P_{rhaBAD}$ – STAR_toehold2 – t500 – Buffer 3 – $P_{lux}$ – anti-STAR_toehold2 – t500 – ColE1 – SpecR | This study |
| | pAB401 | « LLR » : $P_{tet}$ – B0034 – rpaR – B0015 – AraC – $P_{araBAD}$ – STAR_toehold2 – t500 – Buffer 3 – $P_{lux}$ – anti-STAR_toehold2 – t500 – ColE1 – SpecR | This study |
| | pAB518 | J23100 – STAR Target 6 – $RBS_{Bujard}$ – VioB-sfYFP – L3S2P55 – p15A – CamR | This study |
| | pAB519 | J23100 – STAR Target 6 – $RBS_{Bujard}$ – VioB-sfCFP – L3S2P55 – p15A – CamR | This study |
| Supplementary Figure 20B | pAB420 | « Lux Receiver » : $P_{tet}$ – B0034 – luxR – L3S2P55 – $P_{lux}$ – dBroccoli – L3S2P21 – ColE1 – SpecR | This study |
| | pAB421 | « Rpa Receiver » : $P_{tet}$ – B0034 – luxR – L3S2P55 – $P_{lux}$ – dBroccoli – L3S2P21 – ColE1 – SpecR | This study |

All plasmid maps will be made available on Zenodo: https://doi.org/10.5281/zenodo.7757475

### Supplementary Table 4: Model variables

| Variable | Description | Units |
|---|---|---|
| $Q_{11}$ | Concentration of Quorum sensing molecule 1 in cell 1 | nM |
| $Q_{21}$ | Concentration of Quorum sensing molecule 2 in cell 1 | nM |
| $S_1$ | Concentration of STAR in cell 1 | tr |
| $A_1$ | Concentration of Anti-STAR in cell 1 | tr |
| $T_1$ | Concentration of the Target RNA in cell 1 | tr |
| $Co_{SA1}$ | Concentration of STAR-Anti-STAR complex in cell 1 | co |
| $Co_{ST1}$ | Concentration of STAR-Target complex in cell 1 | co |
| $P_1$ | Concentration of Target Protein in cell 1 | nM |
| $Q_{22}$ | Concentration of Quorum sensing molecule 2 in cell 2 | nM |
| $Q_{12}$ | Concentration of Quorum sensing molecule 1 in cell 2 | nM |
| $S_2$ | Concentration of STAR in cell 2 | tr |
| $A_2$ | Concentration of Anti-STAR in cell 2 | tr |
| $T_2$ | Concentration of the Target RNA in cell 2 | tr |
| $Co_{SA2}$ | Concentration of STAR-Anti-STAR complex in cell 2 | co |
| $Co_{ST2}$ | Concentration of STAR-Target complex in cell 2 | co |
| $P_2$ | Concentration of Target Protein in cell 2 | nM |
| $Q_{1m}$ | Concentration of Quorum sensing molecule 1 in the culture medium | nM |
| $Q_{2m}$ | Concentration of Quorum sensing molecule 2 in the culture medium | nM |
| $C_1$ | Concentration of cell 1 in the culture medium | CFU $\mu m^{-3}$ |
| $C_2$ | Concentration of cell 2 in the culture medium | CFU $\mu m^{-3}$ |

### Supplementary Table 5: Model parameters

| Parameter | Description | Value | Units | Source |
|---|---|---|---|---|
| $\rho_o$ | Production rate of the quorum sensing molecules | 595 | nM $\mu m^3$ $h^{-1}$ | Fusco et al. 2002 |
| $\eta$ | Diffusion rate of the quorum sensing molecules | 120 | $\mu m^3$ $h^{-1}$ | Fusco et al. 2002 |
| $D$ | Degradation rate due to dilution | 2 | $h^{-1}$ | Fusco et al. 2002 |
| $y$ | Degradation rate for the RNA species | 13.86 | $h^{-1}$ | - |
| $y_P$ | Degradation rate for the target protein | 0 | $h^{-1}$ | - |
| $a_{So}$ | Production rate of STAR | 6000 | tr $h^{-1}$ | Gorochowskoi et al. 2020* |
| $a_{Ao}$ | Constant term for the production rate of Anti-STAR | 55.368 | tr $h^{-1}$ | Gorochowskoi et al. 2020* |
| $a_A$ | Coefficient for the quorum sensing dependent term of the production rate of Anti-STAR | 50000 | tr $h^{-1}$ | Gorochowskoi et al. 2020* |
| $a_{To}$ | Production rate of the Target RNA | 18000 | tr $h^{-1}$ | Gorochowskoi et al. 2020* |
| $\theta$ | Activation coefficient | 20 | nM | Fusco et al. 2002 |
| $k_{Co_{SA}+}$ | Complex binding rate for $C_A$ complex | 15 | co $tr^{-1}$ $h^{-1}$ | - |
| $k_{Co_{ST}+}$ | Complex binding rate for $C_T$ complex | 1.54 | co $tr^{-1}$ $h^{-1}$ | Fusco et al. 2002 |
| $k_{Co-}$ | Spontaneous unbinding rate of CA and CT complexes | 0.4032 | $co^{-1}$ tr $h^{-1}$ | Fusco et al. 2002 |
| $k_P$ | Production rate of the Target protein | 2.8448 | $co^{-1}$ nM $h^{-1}$ | You et. al. 2004 |
| $k_{C1}$ | Growth rate of cell 1 | 0.5 | $h^{-1}$ | - |
| $k_{C2}$ | Growth rate of cell 2 | 0.5 | $h^{-1}$ | - |
| $d_{B1}$ | Burden produced by target protein P in cell 1 | 0.001 | h $nM^{-1}$ | - |
| $d_{B2}$ | Burden produced by target protein P in cell 2 | 0.0034 | h $nM^{-1}$ | - |
| $D_C$ | Death rate for cell 1 and cell 2 | 0 | $h^{-1}$ | - |
| $V$ | Volume of culture medium | $2.0 \times 10^{11}$ | $\mu m^3$ | - |
| $C_{max}$ | Maximum supported cell concentration for the medium | 0.145 | CFU $\mu M^{-3}$ | Fusco et al. 2002 |

* *values adjusted from Gorochowski et. al. 2020 to compensate for different copy numbers of the plasmids used for the expression of STAR, Anti-STAR and the Target RNA.*

**Supplementary Note 1:** **Modelling and simulations**

The mathematical model developed to represent the system is described by the following set of ordinary differential equations. The equations can be divided into three sets based on the three distinct compartments, namely, within cell 1, within cell 2 and in the culture media. The variables are described in Supplementary Table 4 and the parameters are described in Supplementary Table 5.

In cell 1:

$$\frac{dQ_{11}}{dt} = \rho_o Q_{11} - \frac{\eta(VC_1 Q_{11} - Q_{1m})}{VC_1} - DQ_{11} \quad (1)$$

$$\frac{dQ_{21}}{dt} = \frac{\eta(Q_{2m} - VC_1 Q_{21})}{VC_1} - DQ_{21} \quad (2)$$

$$\frac{dS_1}{dt} = a_{So} + k_{C-}Co_{ST1} + k_{C-}Co_{SA1} - k_{Co_{ST}+}S_1T_1 - k_{Co_{SA}+}S_1A_1 - (y + D)S_1 \quad (3)$$

$$\frac{dA_1}{dt} = a_{Ao} + \frac{a_A Q_{21}^2}{Q_{21}^2 + \theta^2} + k_{Co-}Co_{SA1} - k_{Co_{SA}+}S_1A_1 - (y + D)A_1 \quad (4)$$

$$\frac{dT_1}{dt} = a_{To} + k_{Co-}Co_{T1} - k_{Co_{ST}+}S_1T_1 - (y + D)T_1 \quad (5)$$

$$\frac{dCA_1}{dt} = k_{Co_{SA}+}S_1A_1 - k_{Co-}Co_{SA1} - (y + D)Co_{SA1} \quad (6)$$

$$\frac{dCT_1}{dt} = k_{Co_{ST}+}S_1T_1 - k_{Co-}Co_{ST1} - k_P Co_{ST1} - (y + D)Co_{ST1} \quad (7)$$

$$\frac{dP_1}{dt} = k_P Co_{ST1} - (y_P + D)Co_{ST1} \quad (8)$$

In cell 2:

$$\frac{dQ_{22}}{dt} = \rho_o Q_{22} - \frac{\eta(VC_2 Q_{22} - Q_{2m})}{VC_2} - DQ_{22} \quad (9)$$

$$\frac{dQ_{12}}{dt} = \frac{\eta(Q_{1m} - VC_2 Q_{12})}{VC_2} - DQ_{12} \quad (10)$$

$$\frac{dS_2}{dt} = a_{So} + k_{Co-}Co_{ST2} + k_{Co-}Co_{SA2} - k_{Co_{ST}+}S_2T_2 - k_{Co_{SA}+}S_2A_2 - (y + D)S_2 \quad (11)$$

$$\frac{dA_2}{dt} = a_{Ao} + \frac{a_A Q_{12}^2}{Q_{12}^2 + \theta^2} + k_{Co-}Co_{SA2} - k_{Co_{SA}+}S_2A_2 - (y + D)A_2 \quad (12)$$

$$\frac{dT_2}{dt} = a_{To} + k_{Co-}Co_{ST2} - k_{Co_{ST}+}S_2T_2 - (y + D)T_2 \quad (13)$$

$$\frac{dCA_2}{dt} = k_{Co_{SA}+}S_2A_2 - k_{Co-}Co_{SA2} - (y + D)Co_{SA2} \quad (14)$$

$$\frac{dCT_2}{dt} = k_{Co_{ST}+}S_2T_2 - k_{Co-}Co_{ST2} - k_P Co_{ST2} - (y + D)Co_{ST2} \quad (15)$$

$$\frac{dP_2}{dt} = k_P Co_{ST2} - (y_P + D)Co_{ST2} \quad (16)$$

In the culture media:

$$\frac{dQ_{1m}}{dt} = \eta(VC_1Q_{11} + VC_2Q_{12} - 2Q_{1m}) - DQ_{1m} \quad (17)$$

$$\frac{dQ_{2m}}{dt} = \eta(VC_2Q_{22} + VC_1Q_{21} - 2Q_{2m}) - DQ_{2m} \quad (18)$$

$$\frac{dC_1}{dt} = \frac{k_{C1}C_1(1-\frac{C_1+C_2}{C_{max}})}{1+d_{B1}k_P Co_{ST1}} - D_C C_1 \quad (19)$$

$$\frac{dC_2}{dt} = \frac{k_{C2}C_2(1-\frac{C_1+C_2}{C_{max}})}{1+d_{B2}k_P Co_{ST2}} - D_C C_2 \quad (20)$$

Equations (1), (2), (9) and (10) describe the production and diffusion of the quorum sensing molecules within the cells. Equations (3) and (11) describe the STAR production within the cells. Due to the difference in the hybridization energies of the STAR-Anti-Star complex and the STAR-Target RNA complex the rates of binding are taken to be different, while the rate of unbinding of the complex is assumed to be the same for simplicity. Equations (4) and (12) describe the production of anti-STAR within the cells dependent on the quorum sensing molecules. The Hill coefficient for the activation of Anti-STAR production by the quorum sensing molecule is assumed to be 2. Equations (5) and (13) describe the production of the Target RNA within the cells. Equations (6), (7), (14) and (15) describe the STAR-Anti-STAR and STAR-Target RNA complex formation. Equations (8) and (16) describe the production of the burdensome target protein within the cell. Equations (17) and (18) describe the diffusion of the quorum sensing molecules into and out of the culture media. Equations (19) and (20) describe the cell concentrations within the culture medium.
